# Integrated metabolomic, molecular networking, and genome mining analyses uncover novel angucyclines from *Streptomyces* sp. RO-S4 strain isolated from Bejaia Bay, Algeria

**DOI:** 10.1101/2021.12.21.473593

**Authors:** Rima Ouchene, Didier Stien, Juliette Segret, Mouloud Kecha, Alice M. S. Rodrigues, Carole Veckerlé, Marcelino T. Suzuki

## Abstract

Multi-omic approaches have recently made big strides towards the effective exploration of microorganisms and accelerating the discovery of new bioactive compounds. We combined metabolomic, molecular networking, and genomic-based approaches to investigate the metabolic potential of the *Streptomyces* sp. RO-S4 strain isolated from the polluted waters of Bejaia Bay in Algeria. Antagonistic assays against *methicillin-resistant Staphylococcus aureus* with RO-S4 organic extracts showed an inhibition zone of 20 mm by the agar diffusion method, and its minimum inhibitory concentration was 16 μg/mL. A molecular network was created using GNPS and annotated through the comparison of MS/MS spectra against several databases. The predominant compounds in the RO-S4 extract belonged to the angucyclines family. Three compounds were annotated as known metabolites, while all the others were putatively new to Science. Notably, all compounds had fridamycin-like aglycones, and several of them had a lactonized D ring analogous to that of urdamycin L. The whole genome of *Streptomyces* RO-S4 was sequenced to identify the biosynthetic gene cluster (BGC) encoding for these angucyclines, which yielded a draft genome of 7,497,846 bp with 72.4% G+C content. Subsequently, a genome mining analysis revealed 19 putative biosynthetic gene clusters, including a grincamycin-like BGC with a high similarity to that of *Streptomyces* sp. CZN-748 previously reported to also produce mostly open fridamycin-like aglycones. As the ring-opening process leading to these compounds is still not defined, we performed comparative analysis with other angucycline BGCs and advanced some hypotheses to explain the ring-opening and lactone formation, possibly linked to the uncoupling between the activity of GcnE and GcnM homologues in the RO-S4 strain. The combination of metabolomic and genomic approaches greatly improved the interpretation of the metabolic potential of the RO-S4 strain.

## Introduction

The emergence of novel mechanisms of antimicrobial resistance is increasing and spreading worldwide, posing a challenge to mankind. The World Health Organization has stated that antibiotic resistance will be one of the biggest threats to human health in the future^1^. Multidrug-resistant organisms have become common not only in hospital settings but also in the wide community settings, suggesting that reservoirs of antibiotic-resistant bacteria are present outside hospitals (reviewed by Munita and Arias)^2^. This antibiotic resistance crisis has been attributed to overuse and inappropriate use of these drugs, as well as the lack of antimicrobial drug development by the pharmaceutical industry due to reduced economic incentives and difficult regulatory requirements^3,4^. Methicillin-resistant *Staphylococcus aureus* (MRSA) is the most common cause of nosocomial infections as it is very capable of developing antibiotic resistance^5,6^. Many challenges are faced by laboratories and clinicians in the diagnosis and treatment of MRSA infections, some of which were highlighted by Edwards and coworkers^7^. It is thus clear that the search for new bioactive compounds to combat antimicrobial resistance is a research priority.

Marine environments represent a largely unexplored source for the isolation of new microorganisms^8^. They display a unique combination of environmental conditions and organisms with distinct metabolic capabilities to adapt and thrive^9,10^. A large number of bioactive compounds have been isolated from marine organisms^11,12,13^, particularly Actinobacteria, which have been a main source of natural products in the past^14,15,16^. Among the latter, the *Streptomyces* genus is well known for its ability to produce a wide range of bioactive metabolites as well as antibacterial, anticancer, antifungal, antiparasitic, and immunosuppressive agents^17,18^, representing the most prolific source of bioactive metabolites that have been approved for clinical use, notably as antibiotics^19^.

Traditionally, activity-guided fractionation of metabolite extracts, followed by purification and characterization of metabolites, has commonly been used for natural product research, but this approach often leads to the isolation of already known molecules. More recently, significant developments in genetics, genomics, and data analysis have greatly changed natural product research, leading to a new era in the emerging field of systems biology. Consequently, new avenues were opened for the discovery of novel compounds from actinomycetes (e.g.^20,13,21,22,23,16^). Interestingly, metabolomics and genomics approaches have proven to be efficient and promising tools for defining phenotypes in a dynamic context, with the potential to reduce rediscovery rates^24,25,26^, and several tools have been designed for this purpose, as reported by Caesar and colleagues^27^. These approaches have been successfully applied to study the chemical diversity of marine bacteria and to uncover novel bioactive molecules^28,29,30^, despite the challenges encountered due to the complexity of biological matrices^31^. More recently, molecular networking, a tandem mass spectrometry (MS/MS) data organizational approach, has been introduced in the field of drug discovery^32^, and the combination of system analyses involving multi-omics data and genome-scale, metabolic network models has greatly contributed to exploring bioactive Actinobacteria^33^, and have great potential to accelerate natural product discovery.

Here, we investigated the secreted metabolome of *Streptomyces* sp. RO-S4 in the quest for novel antimicrobial compounds against antibiotic-multi-resistant *S. aureus* (MRSA). For this purpose, we used a metabolomic approach based on ultra-performance high resolution tandem mass spectrometry (UPLC-HRMS/MS) followed by the creation of a molecular network using the Global Natural Product Social Molecular Networking (GNPS) analysis. These analyses were combined with a genomic analysis to refine and further annotate the structural hypothesis generated, and conversely, to understand the biosynthesis of the major angucyclines produced by this strain.

## Results

### Isolation and antimicrobial assays of the RO-S4 strain

The RO-S4 strain was isolated from the bay of Bejaia City in Algeria. It grew well on the M2 medium, showing substrate growth typical for *Streptomyces* strains, with a brown powdery aspect and producing a dark-brown pigment (Fig. 1a). Antimicrobial activity was first evaluated against the MRSA strain by the agar diffusion method. It exhibited antagonistic activity against this bacterium with an inhibition zone estimated at 20 mm (Fig. 1b). The 16S rRNA gene sequence indicated that the strain belongs to the *Streptomyces* genus, with 99.79% identity to *Streptomyces albogriseolus* NRRL B-1305 (T). The MIC of the ethyl acetate extract produced by the RO-S4 strain was measured by the broth microdilution method on a 96-well plate. A MIC value of 16 µg/mL was observed against the MRSA ATCC 43300 strain.

**Figure 1.**
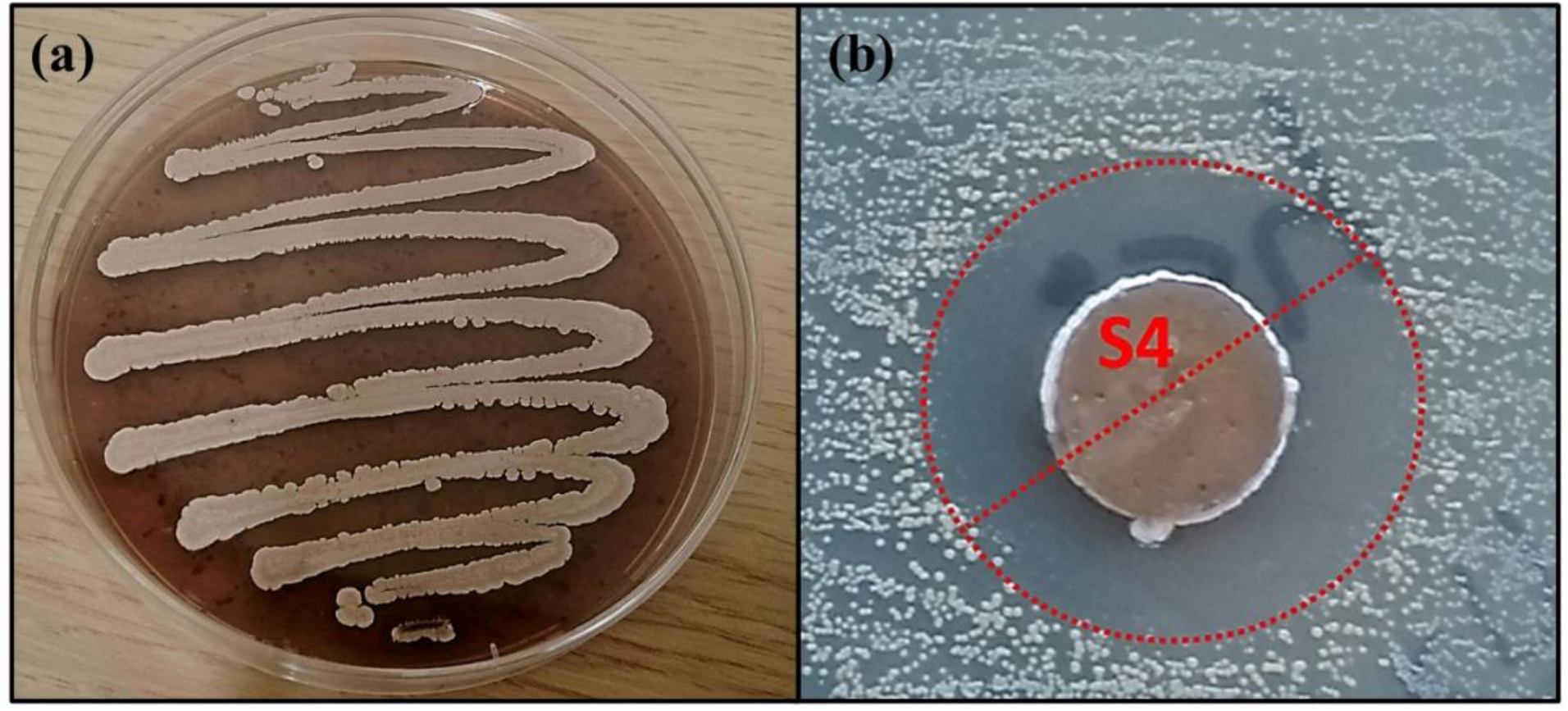
**(a):** The morphological appearance of Streptomyces sp. RO-S4 strain grown on M2 medium for 12 days at 28 °C. **(b)**: MRSA inhibitory potential of the RO-S4 strain evaluated by the agar diffusion method.

### Untargeted metabolomic analysis and Molecular Networking

The metabolomic profile of the active ethyl acetate (EtOAc) crude extract was investigated using UPLC-HRMS. An examination of the MS and collision induced MS/MS (MS^2^ hereafter) spectra of the main constituents of the mixture indicated that most metabolite molecular ions fragmented to yield a product at *m/z* 487.1600 corresponding to the formula C_25_H_27_4O_10_ ^+^ (Calcd. 487.1599). The same ion was also detected as a protonated molecular ion corresponding to compound **1** (Table 1). Compound **1** was annotated either as aquayamycin or as fridamycin A or B by various dereplication tools. A comparison with experimental spectra from the MoNA database confirmed that compound **1** was fridamycin A or its diastereomer fridamycin B. The MS^2^ spectra of fridamycin A and aquayamycin are very similar (Fig. S3, Supporting Information). Nevertheless, two fragment ions are diagnostic. These are the ions at *m/z* 347.09 and 427.14, the relative intensities of which are very low in fridamycin A when compared to those of aquayamycin.

**Table 1.**
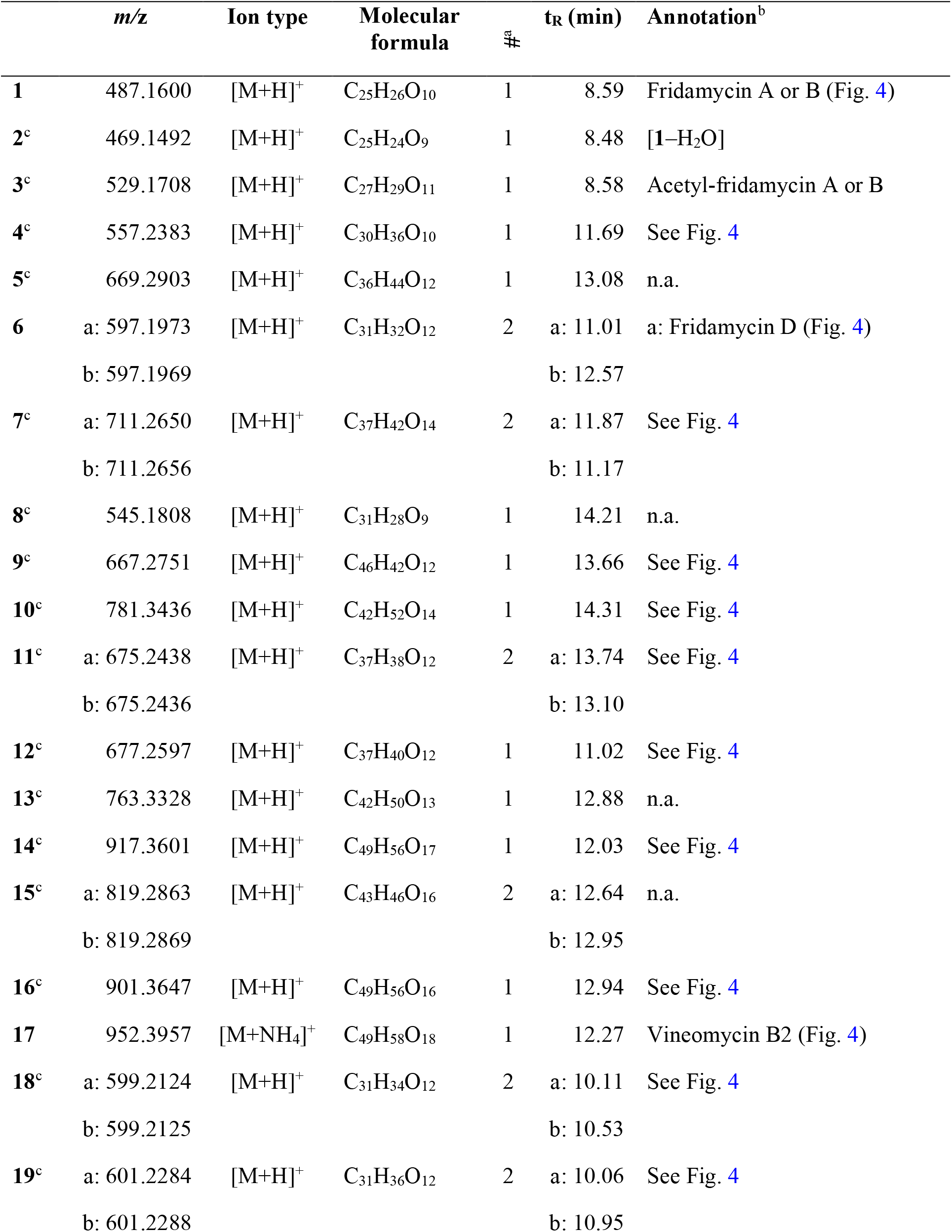
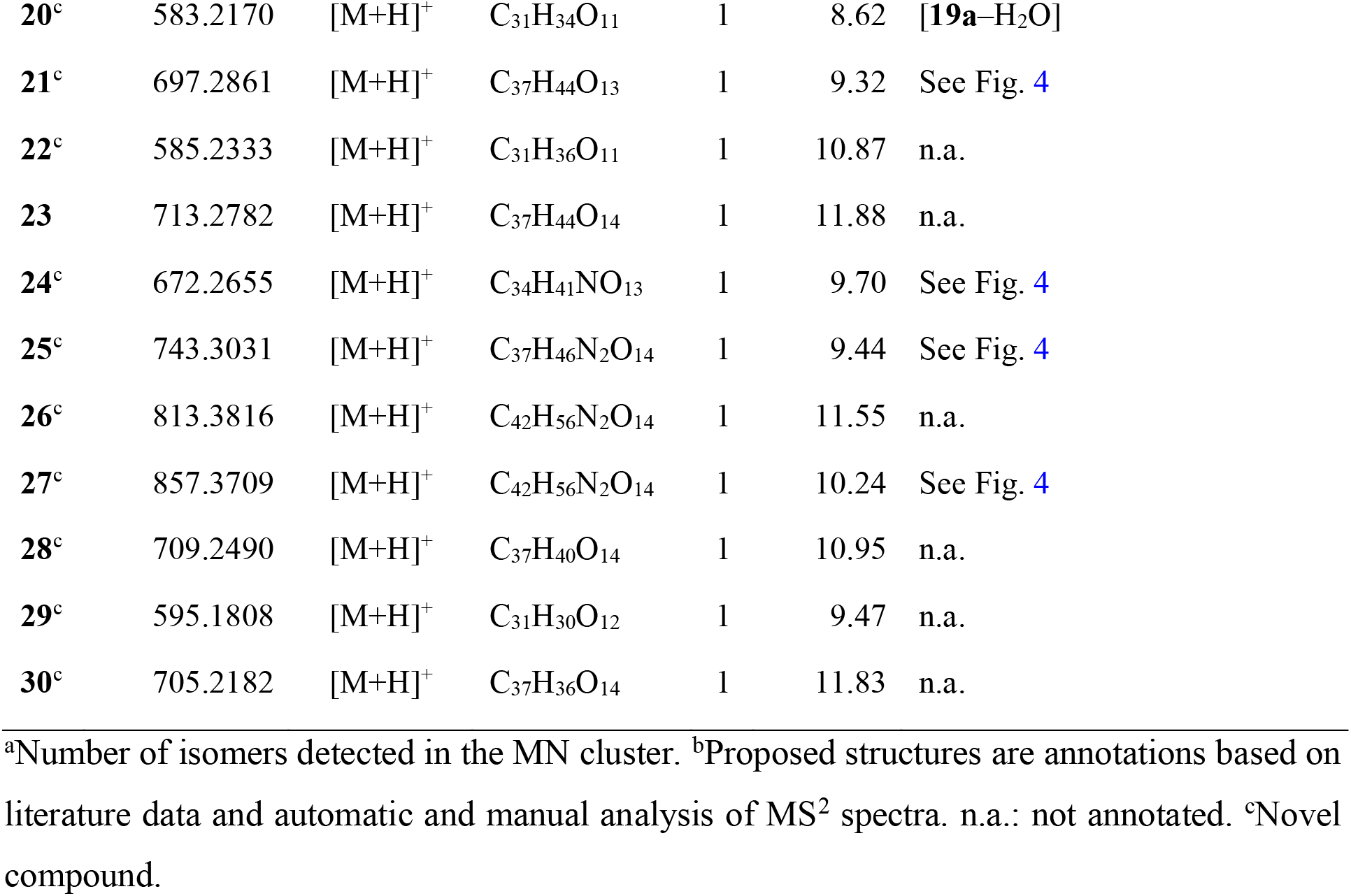
Annotated angucyclines in the metabolomic profile of strain RO-S4

As mentioned above, the [**1**+H]^+^ ion was also produced as a fragment resulting from in-source fragmentation of many metabolites in the profile. The MS^2^ spectra of all ions leading to a 487.1599 fragment clustered in the same node in the molecular network (MN). The MS^2^ spectrum obtained for the protonated molecular ion of compound **1** was compared to all other MS^2^ spectra of equimassic ions found whenever other more complex metabolites were fragmented in source. The fragmentation patterns were all similar, confirming that many metabolites in the MN were fridamycin A or B analogues. It was deduced that the strain biosynthesizes the central fridamycin core and then adds various substituents to generate its diverse products. Notably, the pentacyclic aquayamycin subunit was not detected in any of the annotated metabolites. This observation was supported by the presence of a biosynthetic gene cluster very close to that of *Streptomyces* sp. CNZ-748^34^ which also produces a majority of fridamycin-like compounds.

The parameters for MN were set to construct the best representative network containing all fridamycin analogs (Fig. 2). The MN was constituted of three groups of ions. In the first group, the annotation was propagated from **1** as follows. Compound **2** is a dehydrofridamycin based on its molecular formula and MS^2^ spectrum. Compound **3** molecular formula was C_27_H_28_O_11_, which might be annotated as an acetyl-fridamycin A or B, while the position of the acetyl group could not be inferred from MS^2^. Compound **4** whose molecular formula was C_30_H_36_O_10_ (*m/z* for [M+H]^+^ 669.2903, calcd. 669.2905) was a fridamycin bearing a C_5_H_10_ substituent. A fridamycin isopentyl ester was thought to be a reasonable putative structure based on the biosynthetic considerations below. Compound **5** could not be annotated more precisely than just with its molecular formula, but its MS^2^ spectrum is also one of a fridamycin analog. These considerations indicated that **5** was new to Science.

**Figure 2.**
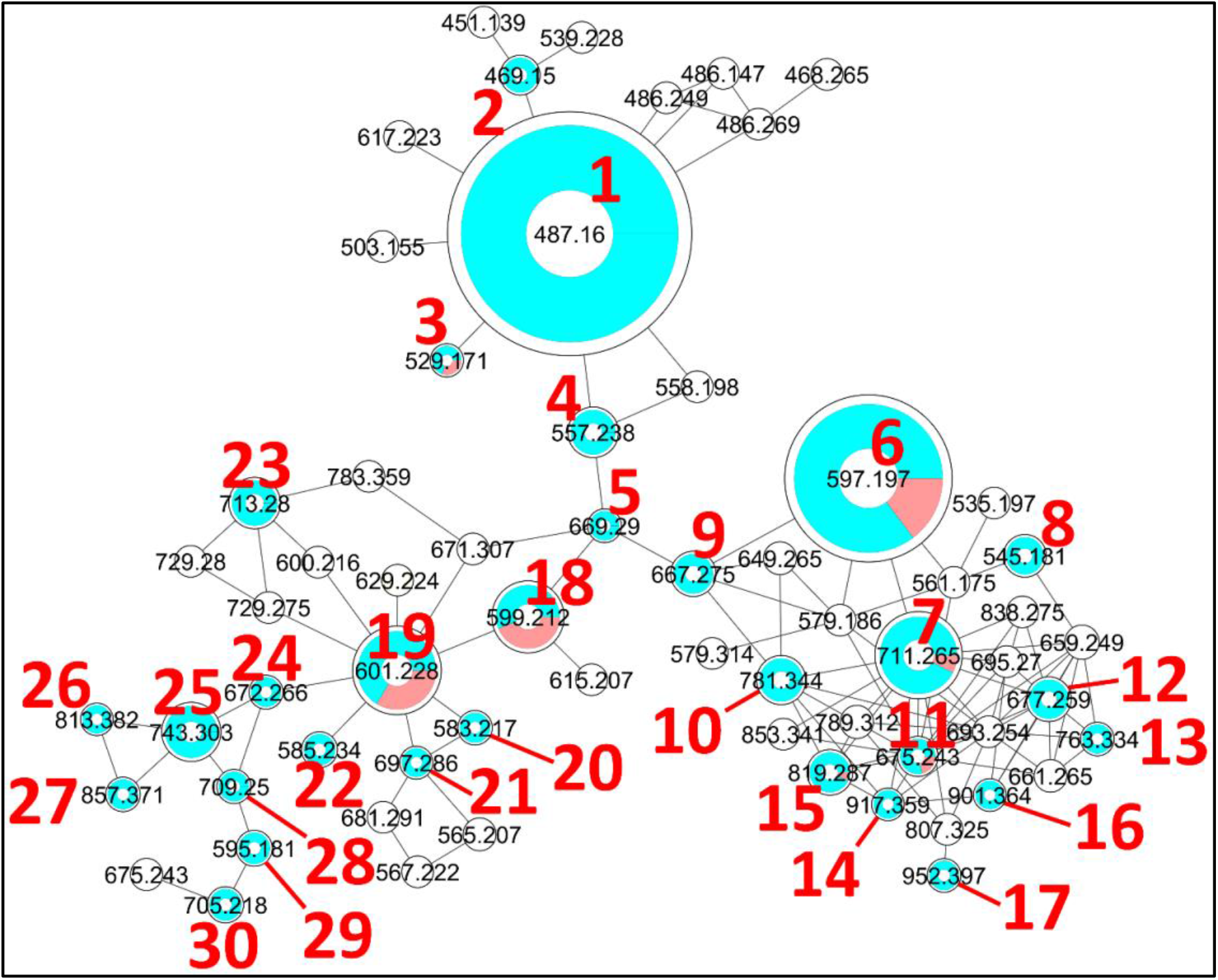
Molecular network generated with the GNPS Molecular Networking tool. The diameter of the nodes represents the total extracted ion chromatogram integration of the corresponding ion peak(s) and the blue/red pie chart represents the proportions of each isomer in the cluster. The colorless nodes are clusters of MS2 spectra of ions produced by in-source fragmentation of diverse compounds and were thus neither annotated nor integrated (integration was set to 0). Only protonated molecular ions were considered for integration measurement.

In the second part of the fridamycin MN, compound **6a** protonated molecular ion was at *m/z* 597.1973, corresponding to the formula C_31_H_33_O_12+_ (Calcd. *m/z* 597.1967). The formula and fragmentation pattern were consistent with the annotation of **6a** as fridamycin D^35^. Annotation as fridamycin D was also supported by Sirius. An isomer of **6a** (**6b**) was also detected in smaller relative proportions in the strain’s metabolomic profile. Another major constituent of the strain metabolome was compound **7a**, whose protonated molecular ion at *m/z* 711.2645, corresponded to the formula C_37_H_43_O_14+_ (Calcd. 711.2647). The molecular formula indicated that angucycline **7a** may be vineomycin C^36^. It was annotated as vineomycin C by Sirius as well, although with 57% confidence. In the collision-induced MS^2^ spectrum of **7a** protonated molecular ion, the rhodinose (or its distereoisomer amicetose) and the aculose oxonium ions were present at *m/z* 115.0756 and 111.0443 (Table S1), respectively, while the presence of the fridamycin A aglycone was ascertained based on the common fridamycin A fragment ions (Fig. S16, Supporting Information). Nonetheless, the first fragmentation steps in the MS^2^ spectrum suggested that the sugar sequence might be different to that of vineomycin C. The protonated molecular ion lost both water (*m/z* forC_37_ H_41_ O_13_ ^+^ 693.2258) and a C_6_H_12_O_3_ group (*m/z* for C_31_ H_31_O_11_ ^+^ 597.1938) which could only be assigned to the rhodinose/amicetose moiety. Hence, the rhodinose could not be placed in between the aglycone and the aculose, as in vineomycin C, and compound **7a** must be considered new to Science, unless the published structure of vineomycin C requires revision. Possible annotations are reported in Fig. 3. These alternative proposals could account for the preferential fragmentation observed in the MS^2^ spectrum. The proximity of **7a** with fridamycin D (**6a)** in the MN suggested that the most probable annotation might be the one in which aculose underwent a Michael addition, as in **6a**.

**Figure 3.**
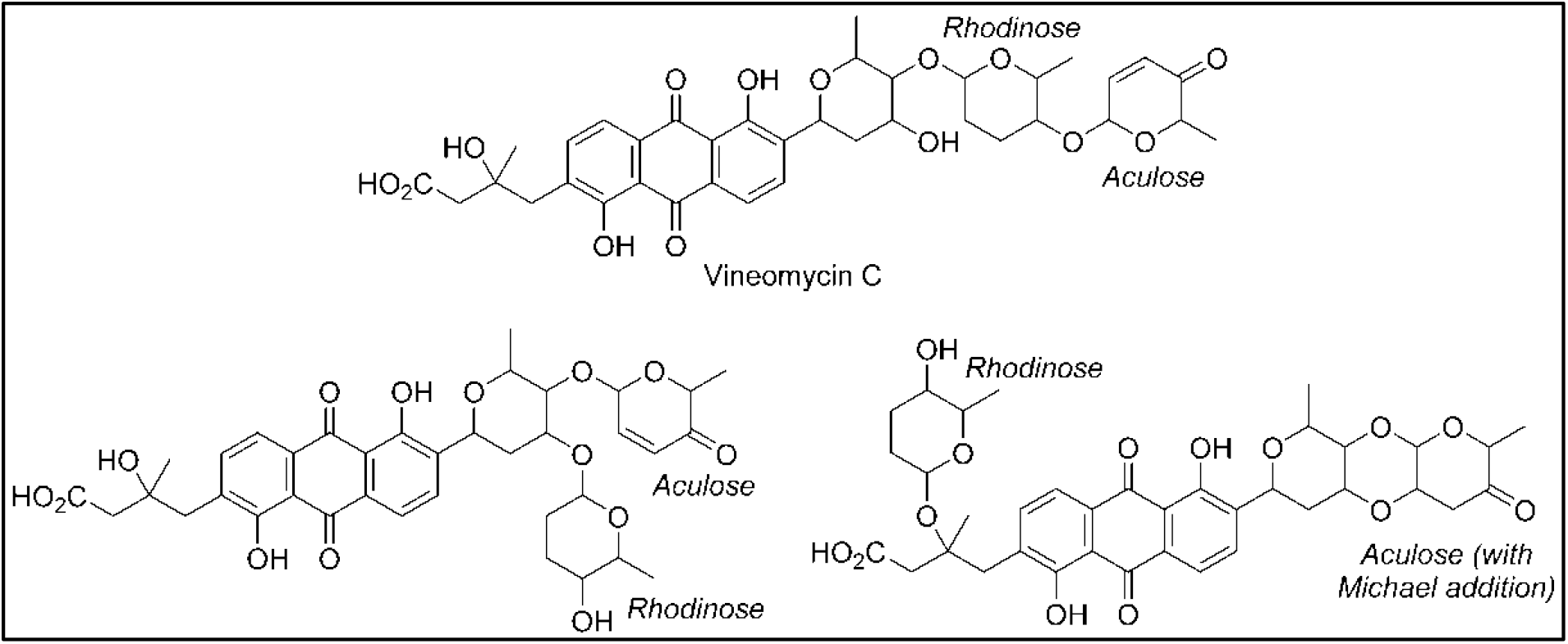
Published structure of vineomycin C and putative structures for compound **6a**. The sugar groups annotated as rhodinose are either rhodinose or amicetose (indistinguishable stereoisomers in MS).

A minor isomer of **7b** also appeared in the MN under the same cluster. Based on the same considerations as for **7a, 7b** could not be annotated as vineomycin C either. It was therefore new to Science. Next to both **6** and **7**, compound **8** protonated molecular ions at *m/z* 545.1808 corresponded to the formula C_31_H_29_O_9+_ (Calcd. 545.1806). The formula suggested that **8** might be marangucycline B^37^. Sirius also annotated this compound as marangucycline B, although with 58% confidence only. Marangucycline B is an analog of fridamycin D with a modified aglycone. The presence of an aculose subunit has been confirmed by MS^2^ (*m/z* 111.0442), but the fragmentation pattern was not attributable to marangucycline B. Hence, while this compound is probably not marangucycline B it could not be annotated further. Angucycline **9** was also new to Science. Its protonated molecular ion at *m/z* 667.2753 corresponded to the formula C_36_H_43_O_12+_ (Calcd. 667.2749). In the MS^2^ spectrum, the fragmentation from the protonated molecular ion to product at *m/z* 579.1851 corresponded to the loss of a neutral fragment the formula of which was C_5_H_12_O (Fig. S22, Supporting Information). This group has been annotated above as an isopentanol, probably linked to the carboxylic acid moiety. A carbon monoxide loss was also detected from *m/z* 561.1750 to 533.1812, indicating that the carboxylic acid side chain of the aglycone should be present. The MS^2^ spectrum also showed a hydrated aculose oxonium ion at *m/z* 129.0549 and its dehydrated form at *m/z* 111.0443. For all these reasons, compound **9** was annotated as shown in Table 1. The protonated molecular ion of compound **10** was found at *m/z* 781.3438, a mass that corresponds to the formula C_42_H_53_O_14+_ (Calcd. 781.3430). An isopentanol, a rhodinose/amicetose, and an aculose were pointed out in the fragmentation spectrum of the parent ion. Rhodinose and aculose oxonium ions were also detected at *m/z* 115.0754 and 111.0442, respectively, showing that **10** should be annotated as the isopentanol ester of **7a/b**. The formula of compound **11a** protonated molecular ion was found to be C_42_H_53_O_14+_ (exp. *m/z* 675.2438, calcd. 675.2436). This corresponded to the grincamycin H molecular formula^38^. Nonetheless, the MS^2^ spectrum indicated that **11a** successively lost rhodinose/amicetose and aculose, and therefore could not be annotated as grincamycin H. The mass of the protonated aglycone after oses fragmentation was *m/z* 451.1388, corresponding to a doubly dehydrated fridamycin A. We hypothesized that one hydroxyl group of the olivose side chain might be dehydrated, and that the second H_2_O loss might be explained by a ring closing of the lactone as in urdamycin L^39^, a possibility supported by the analysis of the biosynthetic gene cluster discussed below. Compound **11b** was presumably a minor diasteroisomer of **11a** due to the high similarity of their respective MS^2^ spectra. Compound **12** protonated molecular ion at *m/z* 677.2597 indicated the formula C_37_H_41_O_12+_ (Calcd. 677.2593). The MS^2^ fragmentation pattern showed the successive losses of rhodinose/amicetose, and aculose. The masses of the fragments generated by cleavage of the aglycone were all shifted by 2 Da relative to those of compound **11a**, indicating that the two additional hydrogens were in the center of the aglycone. Thus, it could be deduced that **12** was likely the hydroquinone form of **11a**. Compound **13** is a minor metabolite and could not be annotated with reasonable confidence. The molecular formula of compound **14** was C_49_H_56_O_17_ (exp. *m/z* 917.3601 ([M+H]^+^), calcd. 917.3590). The MS^2^ spectrum indicated a loss of an aculosyl-rhodinose (or aculosyl-amicetose) moiety. Then the product at *m/z* 675.2443 lost the neutral group C_6_H_8_O (a dehydro-rhodinose), indicating that the aculosyl-rhodinose moiety was linked to another rhodinose. Then the aglycone ion at *m/z* 451.1398 was produced by the loss of another aculose. The molecular weight of this aglycone ion (fridamycin A -2 H_2_O) along with the proximity of other lactonic aglycones in the MN spoke in favor of **14** also being a lactone derivative, as reported in Table 1. The fragmentation spectrum of compound **15** was not very clear and **15** could not be annotated with enough confidence. At *m/z* 901.3653, compound **16** protonated molecular ion indicated the formula C_49_ H_57_ O_16_ ^+^ (calcd. 901.3641). In MS^2^, the ion **16**+H^+^ lost an aculosyl-rhodinose group to give the product at *m/z* 659.2499, which then lost dehydro-rhodinose. Further dehydration produced an ion at *m/z* 545.1790 which again lost aculose to yield the aglycone ion at *m/z* 435.1471. This aglycone was doubly dehydrated compared to fridamycin A, indicating that the *C*-olivosyl group in **16** may be dehydrated. The structure proposed in Table 1 appeared to be a reasonable hypothesis for **16**. The molecular formula of compound **17** is C_49_H_58_O_18_ (exp. *m/z* 952.3960 ([M+NH_4_]^+^), calcd. 952.3961). This molecular formula and sodium adduct fragmentation pattern in which two successive aculosyl-rhodinose losses were recorded were compatible with the annotation of **17** as vineomycin B2^40^.

In the third part of the fridamycin MN, compounds **18a** and **18b** were annotated as isomers with the molecular formula C_31_H_34_O_12_ (exp. *m/z* 599.2124 for [M+H]^+^, calcd. 599.2123). Both isomers fragmented extensively in the ESI source to produce the fridamycin A protonated molecular ion [**1**+H]^+^ losing C_6_H_8_O_2_, *i*.*e*., a dehydro-cinerulose A moiety. The cinerulose A oxonium ion was also detected in MS^2^, while the aglycone fragmentation was very similar to what was recorded for [**1**+H]^+^. Angucyclins **18a** and **18b** were therefore annotated as cinerulosyl-fridamycin A or B. The position of the cinerulosyl side chain was not determined and may not be identical for both isomers. Compounds **19a** and **19b** were annotated as isomers with the molecular formula C_31_H_36_O_12_ (exp. *m/z* 601.2284 for [M+H]^+^, calcd. 601.2279). Both isomers also fragmented extensively in the ESI source to produce the fridamycin A protonated molecular ion [**1**+H]^+^ losing C_6_H_10_O_2_, *i*.*e*., dehydro-rhodinose/amicetose subunit, the corresponding oxonium of which was also present in MS^2^. As mentioned above, the MS^2^ spectrum of the aglycone protonated ion formed by in-source fragmentation was identical to the one of [**1**+H]^+^, therefore confirming that **19a** or **b** should not be annotated as grincamycin L^41^. Instead, **19a** and **19b** were annotated as rhodinosyl-and/or amicetosyl-fridamycin A/B and should be considered as new to Science. Angucycline **19a** was one of the major constituents in the profile of the strain. Compound **20** molecular formula was C_31_H_34_O_11_ (exp. *m/z* 583.2170 for [M+H]^+^, calcd. 583.2174). Its MS^2^ spectrum showed a rhodinose/amicetose subunit and a dehydrated protonated aglycone. Therefore, compound **20** was annotated as a dehydro-**19a**. Compound **21** molecular formula was C_37_H_45_O_13_ (exp. *m/z* 697.2861 for [M+H]^+^, calcd. 697.2855). Its MS^2^ spectrum revealed the successive loss of two rhodinose/amicetose subunits, yielding a dehydrated protonated aglycone. It was thus annotated as shown in Table 1. Compound **22** molecular formula was C_31_H_37_O_11_ (exp. *m/z* 585.2333 for [M+H]^+^, calcd. 585.2330). Its MS^2^ spectrum showed the loss of one rhodinose/amicetose subunit, yielding a protonated deoxy-aglycone. The structure of the aglycone could not be readily inferred from the MS^2^ spectrum and compound **22** could not be annotated further. Compound **23**’s molecular formula was C_37_H_44_O_14_ (exp. *m/z* 713.2782 for [M+H]^+^, calcd. 713.2804). A rhodinose/amicetose was visible in MS, but the MS^2^ spectrum was impure and further annotation was not possible. Compound **24** molecular formula was C_34_H_41_NO_13_ (exp. *m/z* 672.2655 for [M+H]^+^, calcd. 672.2652). Its MS^2^ spectrum showed the loss of 1 rhodinose/amicetose subunit yielding a product at *m/z* 558.1983 (C_28_H_32_NO_11_^+^), which then successively lost water, CH_2_O_2_ (formic acid or H_2_O+CO), and C_2_H_5_N to generate a protonated didehydro-fridamycin A at *m/z* 451.1393. This fragmentation pattern was compatible with **24** being an alanine amide of **19**, as shown in Table 1. The presence of an alanine subunit was also supported by the fragment ion at *m/z* 90.0550, corresponding to an alaninium ion^42^. Compound **25** molecular formula was C_37_H_46_N_2_O_14_ (exp. *m/z* 743.3026 for [M+H]^+^, calcd. 743.3022). In-source fragmentation indicated the successive loss of a dehydro-rhodinose/amicetose group and a C_6_H_10_N_2_O_2_ subunit. The MS^2^ spectrum highlighted the loss of one dehydro-rhodinose/amicetose and two alanine subunits to generate the protonated didehydro-fridamycin A at *m/z* 451.1393 (see Fig. S64, Supporting information). Overall, compound **25** could be annotated with high confidence as shown in Table 1. Neighboring minor compound **26** in the MN could not be annotate. Compound **27** was an analog of **25** with one rhodinose/amicetose subunit more, the presence of which could be ascertained by examination of both in-source fragmentation scheme and the collision-induced MS^2^ spectrum. Compound **28** molecular formula was C_37_H_40_O_14_ (exp. *m/z* 709.2490 for [M+H]^+^, calcd. 709.2491). The MS^2^ spectrum clearly indicated that the protonated molecular ion lost both dehydro-rhodinose/amicetose and a C_6_H_8_O_4_ neutral fragment, therefore confirming that this compound was new and should not be annotated as saprolmycin B^43^. However, the annotation remained ambiguous and **28** was not annotated further. Compounds **29** and **30** were analogs of **28**; **29** did not have the rhodinose/amicetose subunit, while **30** had an aculose in place of the rhodinose/amicetose. All the annotated compounds’ MS^2^ spectra were provided in the supporting information (Figs S1-S74, Supporting Information), and a list of sugars potentially linked to the angucyclins is provided in Table S1.

### Whole genome sequencing

The whole genome of *Streptomyces* sp. RO-S4 was sequenced using the Illumina Novaseq technology. The complete genome consisted of 7,497,846 bp with 72.4% G+C content. The closest genome to *Streptomyces* sp. RO-S4 sequenced from a type strain was that of *S. althioticus* JCM 4344 (assembly GCA_014649355; 90.73% ANI), and that of *S. tendae* strain 139 (CP04395; 95.84%) for a non-type strain. The genome of *Streptomyces albogriseolus* NRRL B-1305, the closest strain based on 16S rRNA gene similarity was not available at the time of analysis.

### Secondary metabolite biosynthetic gene clusters of *Streptomyces* sp. RO-S4

The genome of the RO-S4 strain was analyzed by the antibiotics & Secondary Metabolite shell (antiSMASH) to determine its putative biosynthetic capabilities. A total of 19 putative biosynthetic gene clusters were annotated (Table S2, supporting information), including three types of polyketide synthases BGCs [Type 1 (T1PKS), Type 2 (T2PKS), and type 2 (T3PKS)] polyketide synthases, class I Lanthipeptides, a Lassopeptide, a Ribosomally Synthesized and Post-Translationally Modified Peptide (RiPP), an Ectoine, a Terpene, Phenazines, and a Butyrolactone BGC. In addition, four hybrid clusters were recovered that were composed of 1) T2PKS, Oligosaccharide Phenazine, Siderophore, 2) RiPP-like, betalactam, Terpene, 3) Two hybrid Non-ribosomal Peptide Synthase (NRPS), and T1PKS. The analysis showed that 17 out of the 19 identified BGCs showed high content similarity with known BGCs, five of which (**3, 5, 7, 13**, and **17**) showed 100% content similarity with known BGCs. Two clusters (Cluster **10** and **16**) were annotated as orphan BGCs for which no homologous gene clusters could be identified, suggesting that they could be responsible for the biosynthesis of novel natural products or natural products with no characterized BGCs. Many of these clusters are known to encode genes linked to the production of biologically active natural compounds, such as antibiotics. Notably, we have recovered a T2PKS BGC very similar to those linked to the biosynthesis of angucycline compounds, consistent with the metabolomic analysis.

### Description of the Angucycline Biosynthetic Gene Cluster

AntiSMASH analysis using an unannotated genomic DNA sequence revealed that Cluster 2 in Contig ROS4_2 of the assembly displayed a high synteny (> 65% common genes) to that of BGCs linked to several angucycline compounds, such as grincamycin (97%), saprolmycin E (83%), saquayamycin A (75%), landomycin A (71%), and saquayamycin Z (67%). All these compounds share a common tetracyclic angular benz[a]anthraquinone aglycone. Due to the predominance of tricyclic aglycones (“open” aglycone(s) hereafter) among the major metabolites of RO-S4, we performed an in-depth analysis of Cluster 2.

Cluster 2 contains genes putatively involved in the biosynthesis and modification of the aglycone core. A typical set of genes responsible for an angucycline core assembly named “minimal PKS” has been identified supporting the synthesis of angucycline like molecules by his cluster. This included three genes: a ketoacyl synthase α (LRR80_00487), a ketoacyl synthase β/chain length factor CLF (LRR80_00488) and an acyl carrier protein (ACP) (LRR80_004989). Two possible cyclase genes (LRR80_00486 and LRR80_00491) were also annotated, which are likely responsible for the polyketide chain cyclization into the benz[a]anthracene structure. In addition, this cluster harbors two genes encoding oxygenase enzymes (LRR80_00485 and LRR80_00492) probably involved in the modification of the aglycone and possibly in the lactonization and opening of the angular aglycone cycle (see discussion below). (Keto)reductase-coding genes, including LRR80_00490 and LRR80_00470, were annotated to exhibit a high degree of sequence similarity to known enzymes involved in the modification of aromatic polyketides. Three genes (LRR80_00495, LRR80_00496, and LRR80_00498) likely associated with the glycosylation steps showed high similarity to genes coding glycosyltransferases (GTs) in other angucyclines. All the annotated genes involved in the BGC of Cluster 2 and their homologs are listed in Table S3.

The closest BGCs to RO-S4 Cluster-2 are those of the grincamycin-producing *Streptomyces lusitanus* SCSIO LR32 (Gcn LR32)^44^, and *Streptomyces* sp. CZN-748 (Gcn CZN-748^34^ graciously provided by the authors). We performed a synteny analysis comparing the three BGCs and notably, when PROKKA^45^-annotations were used for RO-S4 and CZN-748, the *gcnM* ORF was absent in these strains, as previously described by Shang and co-workers^34^. In contrast, the BGC annotated by AntiSMASH from genomic sequences identified these ORFs. This difference could be explained by a possible tRNA_ala_ in the region coding the *gcnM* orthologs (Fig. S75, Supporting Information). In addition, the comparison between the BGC of LR32 identified two missing genes (*gcnU* and *gcnT*) in the BGC of ROS4_2, whereas the region downstream of gcnS8 was not present in the available BGC of strain CZN-748 (Fig. 5). Cluster 2 of RO-S4 showed near complete synteny and a higher average amino acid identity (99.8 % for 28 common ORFs) to the grincamycin BGC of CZN-748, which fits the observation that both strains produce a majority of “open” aglycone angucyclines, whereas LR32 (93.2 % average amino acid identity for 28 common ORFs) produces primarily tetracyclic angucyclines, as previously noted by Shang and co-workers^34^.

**Figure 4.**
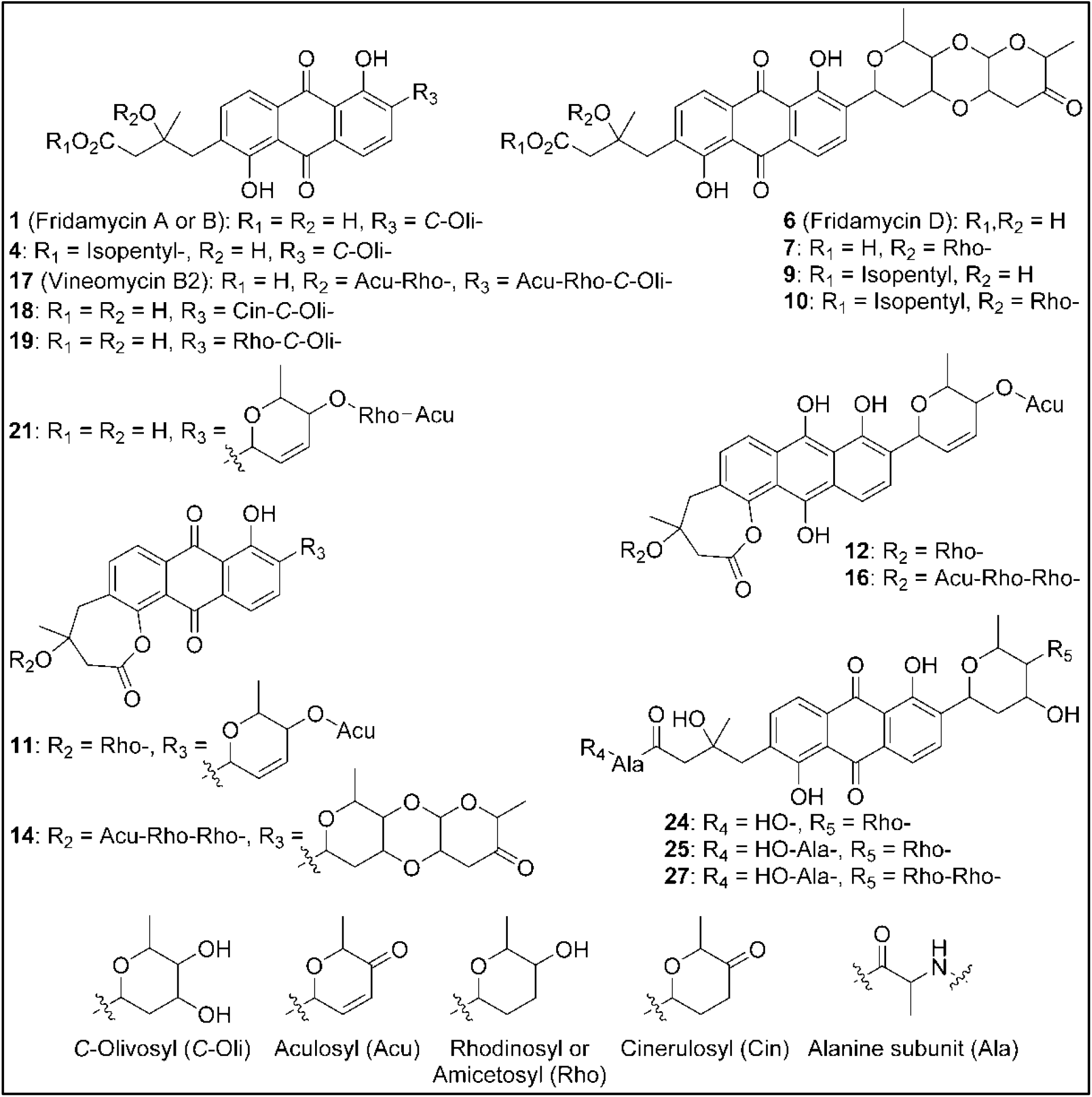
Annotated metabolites from Strain RO-S4. Stereocenters are intentionally drawn as undefined. Only the raw formula of the substituents can be inferred from the mass spectra. Their developed formulas and relative positions are putative.

**Figure 5.**
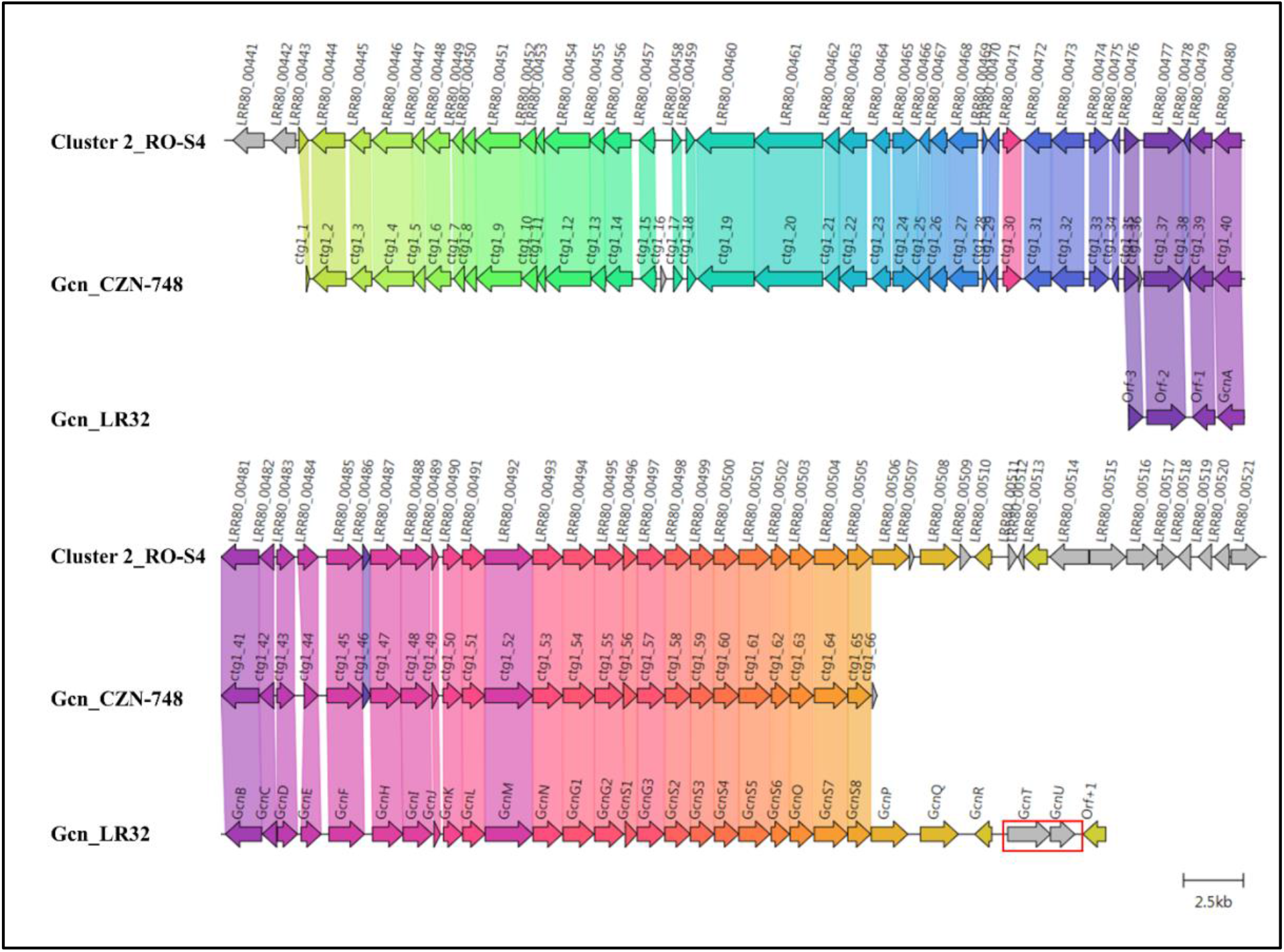
Comparison between Cluster 2 of Streptomyces sp. RO-S4, the Grincamycin Gene Cluster of Streptomyces lusitanus SCSIO LR32, and the BGC of Grincamycin-producing Streptomyces sp. CNZ-748. Gene neighborhoods representative of the compared BGCs are shown aligned with an arbitrary color scheme using clinker to highlight the conserved genes. Missing genes in RO-S4 compared to LR-32 were highlighted in red.

## Discussion

Here we report the use of a combined genomic-metabolomic approach to investigate the antagonistic potential of the *Streptomyces* sp. RO-S4 strain isolated from a polluted marine environment. Based on 16S rRNA gene sequencing and genomic analysis, the strain belongs to the genus *Streptomyces*, but it was not possible to assign it to a species. RO-S4 extracts show inhibitory activity against MRSA with a MIC of 16 µg/mL.

Metabolomic analyses of the crude extract produced by the RO-S4 strain using mass-spectrometry-based molecular networking revealed diverse angucycline derivatives as dominant products, which have mostly (but not exclusively) been linked to the *Streptomyces* genus^46,47^. Angucyclines represent the largest group of type 2 PKS natural products produced by actinobacteria, and they show diverse pharmacological activities including cytotoxicity, antitumor, antibacterial, and antiviral properties^48,49^.

We have demonstrated that many of the compounds identified by our untargeted metabolomic analysis are novel to Science, and this high diversity of novel molecules can be explained by the ability of HRMS to highlight minor compounds even though some of the major metabolites are also new to Science. Our annotation of the RO-S4 angucycline-like compounds was further supported by genomic analysis.

The combination of metabolomic and genome analysis provides some interesting insights into the biosynthesis of angucyclines. Cluster 2 shows high content, synteny, and sequence similarity to previously described BGCs of grincamycin-like products and is certainly responsible for producing the structures predicted by the MN analysis. Since the RO-S4 strain and *Streptomyces* CNZ-748 produce primarily tricyclic glycosylated structures, while LR32 and many other strains produce tetracyclic angucyclines, we were interested in possible enzymatic processes that could lead to the biosynthesis of these tricyclic aglycons^34^. Several studies have indicated that oxygenase complexes are required for cyclic C-C bond cleavage, in particular Baeyer-Villiger type oxygenases^50^. In the case of RO-S4 products, we hypothesized that this reaction would take place by an oxidation of the C12b–C1 single bond of the aglycone (an UWM6-like molecule), prior or after glycosylation, and a subsequent hydrolysis of the lactone (Figure 6). We have therefore focused the analysis on putative Baeyer-Villiger mono-oxidases (BVMOs) in Cluster 2 of the RO-S4 strain.

**Figure 6.**
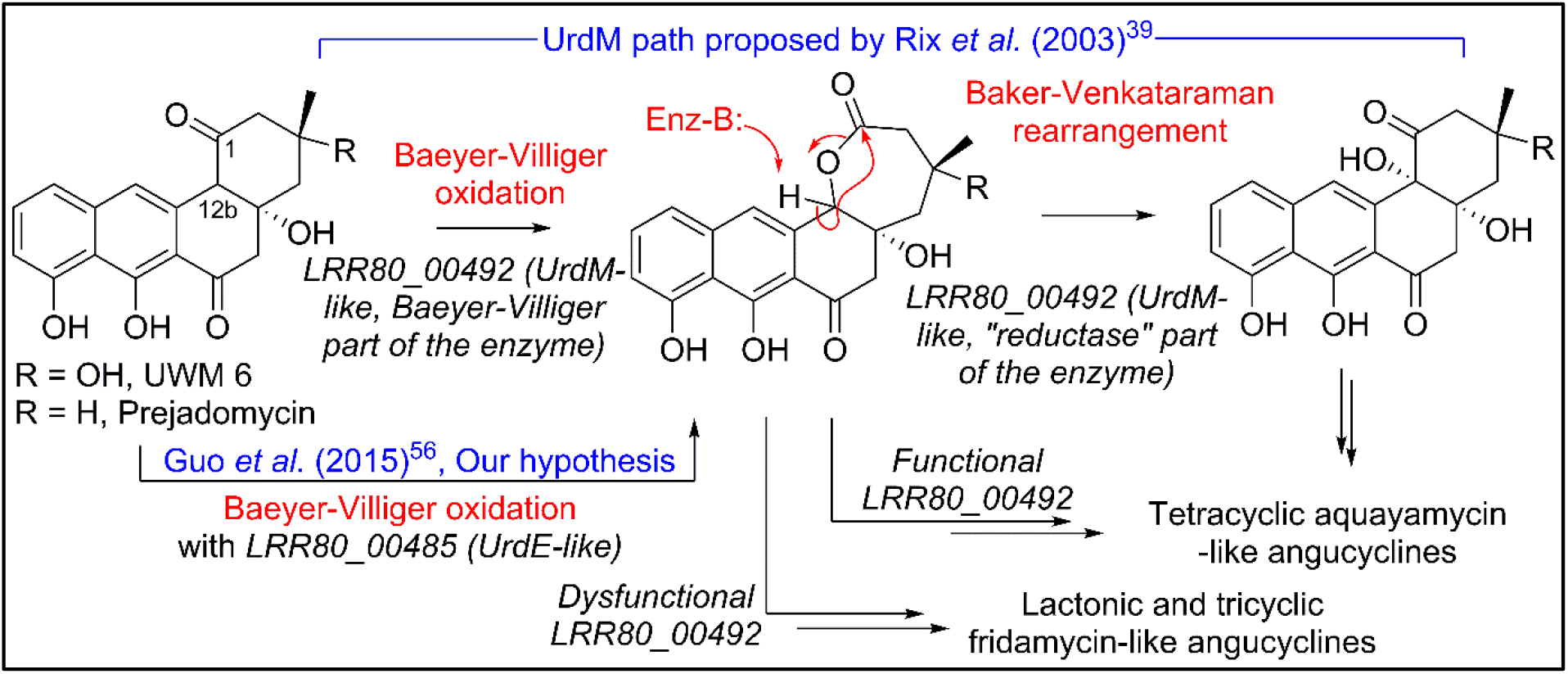
Hypotheses for conversion of UWM 6 or prejadomycin into angucyclines.

The observation that – as in the case of the grincamycin BGC of strain CZN-748 – annotation with PROKKA^45^ failed to identify an ORF downstream of the T2PKS synthases and the cyclase putatively involved in the generation of the angular cycle (the region coding for GcnM in *S. lusitanus* LR32^44^), suggested that this ORF could be involved in the ring opening process. Since AntiSMASH identified an ORF both in RO-S4 (LRR80_00492) and CNZ-748 (CTG1-52) when a genomic sequence was used as the query, we determined that PROKKA failed to annotate the ORF since its Aragorn^51^ step identified a putative tRNA_ala_ (Fig. S75, Supporting Information) in its complementary strand. However, the facts that the putative tRNA is in the reverse strand, that it contains mismatches in the side harpins, and that 4 more canonical tRNA_ala_ are coded in the genome, suggest that this tRNA_ala_ could have been misidentified. We performed an RNA fold analysis^52^ that identified a highly probable and low-entropy hairpin loop that could potentially affect the translation of this ORF (Fig. S76, Supporting Information) and possibly decreases the production of the coded protein.

The LRR80_00492 ORF codes a hybrid FAD-dependent oxidase-reductase (GcnM in the grincamycin BGC) homologous to UrdM, that has been linked to C12b hydroxylation in the biosynthesis of urdamycins by *S. fradiae* TÜ 2717^53,39^. Furthermore, a mutant with an in-frame deletion of the reductase domain of UrdM produced small amounts of urdamycin-L, a product containing an oxygen between C12b and C1, leading to the hypothesis that UrdM is involved in the C12b–C1 bond oxidation and subsequent lactone Baker-Venkataraman rearrangement leading to the tetracyclic skeleton of aquayamycin-like angucyclins (Fig. 6)^39^. However, in this model, low levels, or absence of LRR80_00492 due to the secondary structure described above would not lead to fridamycin-like aglycones as those in RO-S4 (Table 1), and another BVMO should be responsible for the oxidation of the C12b–C1 bond in **UWM 6** (or other intermediates) leading to compounds **11, 12** and, **16** and fridamycin-like aglycones.

The BGC of RO-S4 and of all grincamycin-producing strains contains a second FAD-dependent putative BVMO product of LRR80_00485 (GcnE in the grincamycin BGC) that is homologous to FAD-dependent monooxygenases involved in angucycline modifications (e.g. UrdE, PgaE, BexE, CabE; Fig S77 and S78, Supporting Information). In earlier studies, UrdE was hypothesized to directly hydroxylate different positions of the aglycon (C6, C12, C12b)^53,54,55^ in urdamycin biosynthesis, but more recent evidence have suggested that its homologue PgaE might also oxidize the C12b–C1 bond of prejadomycin leading to the tricyclic aglycons of gaudimycins D and E^56^. *In vitro* assays using enzymes heterologous expressed in *E. coli* also showed that PgaE/CabE oxidizes UWM6 and is dependent on the PgaM_red_ homologue CabV to complete the hydroxylation of UWM6 at C12b^57^.

Since in the RO-S4 BGC there are two possible FAD-dependent mono-oxidases that could be involved in oxidation of the C12b–C1 bond and subsequent ring opening of RO-S4 and CNZ-748, we attempted to compare the sequences of both ORFs to different enzymes oxidizing analogous cyclic compounds, including MtmOIV, shown to perform a Baeyer-Villiger oxidation and ring opening of premithramycin B to mithramycin. *Blastp* analyses indicated that the LRR80_00492 (UrdM like)/mtmOIV alignment was shorter and had a lower overall score, but a higher number of identical positions, whereas the LRR80_00485 (UrdE-like)/mtmOIV alignment was longer and had a higher total score, but with fewer identical positions. Phylogenetic analyses including UrdE homologues and the oxidase portion of UrdM homologues separated these oxidases into two groups, with maximum likelihood and distance methods showing that LRR80_00485 was slightly closer to MtmOIV than LRR80_00492, and in a subclade including PgaE (Figs. S77 and S78, Supporting Information).

In aggregate, these results lead to different possibilities that could be tested in the future using genetic modifications of the different ORFs in the anguclyline BGC of the RO-S4 strain or CNZ-748. Hypothesis 1): the product of LRR80_00492 would be solely responsible for oxidation of the C12b–C1 bond. This hypothesis is supported by the prediction that this enzyme has several AAs unique to RO-S4 and CNZ-748. On the other hand, since the reductase portion of the enzyme is present, one would expect that tetracyclic angucyclines with a hydroxylated C12b would be produced. Hypothesis 2): the most likely hypothesis based on our work on the RO-S4 strain is that the LRR80_00492 ORF is inactive due to the presence of a tRNA*ala* in the coding region or that its translation is affected by secondary structure, in which case LRR80_00485 would generate the lactone via a Baeyer-Villiger oxidation, possibly allowing for a later opening of the ring. Other alternative hypotheses that could be related to the ring opening are: Hypothesis 3): that the LRR80_00492 ORF would be partly transcribed due to the mRNA secondary structure, which would allow its oxidase portion to be transcribed but not the reductase, much like the case with the *urdM* partial knockout mutant that produces urdamycin L^39^ and is responsible for C12b–C1 single bond oxidation, and Hypothesis 4): another BVMO enzyme not in the BGC could be responsible for ring opening. The genome of RO-S4 codes for another enzyme with a slightly higher *blastp* score when queried with MtmOIV. That ORF is present in a hybrid NRPS-T1PKS hybrid BGC similar to that of polyoxypeptin A BGC^58^. However, the function of that homologue (ORF4) in the polyoxypeptin A BGC has not yet been described.

In addition to these oxygenases, three genes are presumed to code for glycosyltransferase (GT) enzymes, which show similarities to known GTs involved in angucycline biosynthesis. The first, RO-LRR80_00495, shows high homology to GcnG1 (96.98%^44^), sqnG1 (88.14%^59^), sprGT1 (86.74%^60^), and SchS10 (81.63%^61^). The second (LRR80_00496) is closely related to GcnG2 (92.46%^44^), sprGT2 (83.92%^60^), sqnG2 (81.91%^59^), and schS9 (74.74%^61^). The third GT (LRR80_00498) is most similar to enzymes involved in the glycosylation of D-olivose at the C9 position of several angucycline-like molecules [ex. SchS7 in the Sch-47554 biosynthetic gene cluster^61^; SprGT3 in saprolmycin biosynthesis^60^; UrdGT2 in urdamycin biosynthesis in *Streptomyces fradiae* Tü 2717^62^, among others], supporting well the predicted structures in Table 1, all of which are predicted to have been glycosylated with an olivose at the C9-position. SchS9 and SchS10 are thought to be *O*-glycosyltransferases involved in the biosynthesis of Sch-47554^61^. Genetic studies using heterologous expression and targeted gene disruption have shown that SchS7 attaches D-amicetose at C-9 and SchS9 further extends the saccharide chain, while SchS10 attaches L-aculose at the C-3 position^63^. The SqnGT1-G3 are glycotransferases involved in the biosynthesis of saquayamycin A in *Streptomyces sp*. KY40-1. According to genetic experimentation, sqnG2 was identified as catalyzing both *O*- and *C*-glycosylations^59^.

Hence, based on similarities between known GTs and the sugars annotated by metabolomic analysis, we hypothesize that a D-olivose is added to C9 by LRR80_00498 and further *O*-glycosylated with rhodinose/amicetose by one of the three glycosylases, and in some cases (ex. compound **9**), experiencing a Michael addition, or further glycosylation as in compounds **19** and **27** (and in previously described angucyclines). Interestingly, all compounds glycosylated at position C3 in RO-S4 appear to have a rhodinose/amicetose as the sugar moiety, as do all grincamycins. It is worth noting that the composition and varying lengths of the oligosaccharide chains in angucyclines have a considerable impact on their biological potential, as previously reported by Elshahawi *et al*.^64^.

In addition to the rare lactonized D cycle, we also detected other modifications that, to the best of our knowledge, have so far not been shown for fridamycin-like molecules, including alanyl amidation(s) (compounds **25**-**28**). Amide modifications have been previously shown in fridamycin G^65^, and fridamycin I produced by *Actinokineospora spheciospongiae*^66^. Fridamycin G contains an ethanolamine moiety and was produced heterologously, and the amidation was attributed to a process linked to the host, since the BGC source (*S. cyanogenus* S136) does not produce it. Fridamycin I contains a benzylamine moiety, the biosynthetic origin of which was not discussed. As compounds **25**-**28** contain multiple alanine moieties, we speculate that fridamycin-like molecules could undergo peptide-like elongation with aminoacids (alanine or L-*p-*hydroxyphenylglycine in the case of fridamycin I), perhaps via an NRPS in a manner analogous to what has been described in the biosynthesis of actinomycin-D^67^. The presence of an *hpgT* homologue – the enzyme linked to L-*p-*hydroxyphenylglycine production in actinobacteria^68^ – in the *Actinokineospora spheciospongiae*, supports this hypothesis.

## Conclusions

Our combination of untargeted UPLC-HRMS/MS metabolomic, molecular networking, and genomic analyses generated many structural and biosynthetic hypotheses for targeted structural determination and genetic manipulation, possibly using recently developed gene editing approaches (e.g^69,70^). This approach does not substitute traditional isolation-NMR structural analyses and genetic manipulations, which ultimately will be needed to confirm these hypotheses. However, as it yields structural information of many minor compounds linked to BGCs in a relatively fast manner, it can streamline and accelerate pipelines of discovery of new drugs, biosynthetic pathways, and enzymes, and hopefully inspire the discovery to novel antibiotics effective against multi-resistant microorganisms.

## Methods

### Isolation of the RO-S4 strain

The RO-S4 strain was isolated from polluted seawater that was collected from the coastline of Bejaia City (36°43’55.2”N5°04’37.9”E), Algeria on August 2017. It was isolated after filtration onto a 0.22 µm pore size membrane filter as described by^71^ and laid onto solid M2 medium prepared according to Jensen *et al*.^72^ with small modifications, which consisted of starch (10 g), casein bovine milk (1 g), microbiological agar (18 g), and natural 100% of seawater (1 L), and 1 mL (per liter of final medium) of trace salt solution that was prepared according to Shirling and Gottlieb^73^.

### Molecular identification of the RO-S4 isolate

The initial molecular identification of the RO-S4 strain was based on the 16S rRNA gene sequence. Genomic DNA was extracted from the grown strain using the Wizard® Genomic DNA Purification Kit (Promega, USA) according to the manufacturer’s instructions. PCR and sequencing were realized as previously described ^74^, utilizing universal primers recommended for bacteria 27F mod: 5’AGRGTTTGATCMTGGCTCAG 3’ and 1492R mod: 5’TACGGYTACCTTGTTAYGACTT 3’. The PCR product was purified with a purification kit (Promega, USA), and then sequenced by the dideoxy termination reaction using an AB3130 DNAxl sequencer. The obtained 16S rRNA sequence was identified by comparison to the EZBiocloud database (https://www.ezbiocloud.net/) recommended by Yoon and co-workers^75^. The RO-S4 16S rRNA gene sequence was deposited in the GenBank database under the accession number (MW448345)

### Antimicrobial Assays

The antibacterial potential of RO-S4 isolate was evaluated against *methicillin-resistant Staphylococcus aureus* (MRSA) ATCC 43300 by the agar diffusion method^76^. 8 mm diameter agar cylinders of the RO-S4 strain (M2 medium, incubation for 14 days at 28 °C) were inserted into Muller Hinton plates previously seeded with the targeted bacterium at 10^7^ UFC/mL. The plates were placed for 2 h at 4 °C and antibacterial activity was estimated by measuring the inhibition zone around the agar disc after incubation of the plates for 24 h at 37 °C.

### Culture strain and the production of raw extract

The production of bioactive compounds by the selected strain was carried out by agar surface fermentation (ASF), according to Nkanga and Hagedorn^77^. Briefly, the RO-S4 isolate was initially grown on M2 agar plates. After 14 days, the mycelium layers were peeled off and extracted overnight in ethyl acetate (EtOAc), covering the entire surface, and the ethyl acetate extract was drawn. Subsequently, the organic extract was concentrated under vacuum with a rotary evaporator at 40 °C and then stored at -80 °C until further analysis. A control, uninoculated medium, was extracted with the same protocol.

### Minimum Inhibitory Concentration (MIC) of the RO-S4 crude extract

The minimal inhibitory concentration (MIC) of the RO-S4 EtOAc extract was evaluated against the MRSA ATCC 43300 strain by the broth microdilution method as recommended by the Clinical and Laboratory Standards Institute^78^. The assays were performed in serial dilutions (in triplicate) in 96-well plates. Briefly, the EtOAc extract was diluted in DMSO and tested at different concentrations ranging from 256 to 0.5 *µ*g/mL. The targeted bacterium culture was prepared in Muller Hinton broth at 2·10^5^ UFC/mL. Afterward, 10 µl of the test bacterial culture was pipetted into each well. The last column (column 12) with no inoculum served as a sterility control, while wells that were not treated with the crude extract served as a negative control (column 11). The final volume of each well was adjusted to 100 µL. The microplate was shaken gently, then incubated for 24 h at 37 °C. Inhibition was evaluated as well, where the growth medium appeared clear, indicating that the test extract prevented the growth or killed the bacteria.

### UHPLC-HRMS Profiling

The protocol for high-resolution Full MS data dependent MS^2^ analyses was adapted from previous reports^79,80^. Here, crude bacterial and culture medium (M2) extracts were dissolved in MeOH at a concentration of 1.5 mg/mL. Pure methanol injections were used as blanks for metabolomics. In HPLC, the solvent system was a mixture of water (solution A) with increasing proportions of acetonitrile (solution B), both solvents modified with 0.1% formic acid. Here, the gradient was as follows: 5% B 5 min before injection, then from 1 to 12 min, a linear increase of B up to 100%, followed by 100% B for 8 min.

### MS/MS Molecular Networking analysis and Spectra Annotation

The molecular network was constructed using the Global Natural Product Social networking (GNPS) platform available at: (https://gnps.ucsd.edu) as recommended by Wang and collaborators^32^, using the molecular networking (MN) tool. The MS^2^ data of the crude extract, solvent (blank) and culture media were converted from RAW to mzXML files using the Proteowizard MSConvert tool version (3.0.20104), then uploaded to GNPS. For MN construction, the precursor ion mass tolerance was set at 0.005 Da and the MS^2^ fragment ion tolerance was set at 0.01 Da. A network was created where edges were filtered to have a cosine score above 0.77 and 11 or more matched peaks. The maximum size of a molecular family was set at 85. The MS^2^ spectra in the network were searched against the ‘GNPS spectra library’. All matches between network and library spectra were required to have a score above 0.7 and at least 6 matched peaks. Visualization of the molecular network was performed in Cytoscape (3.8.0) which allowed its visualization as a network of nodes and edges^81^. Redundancies and adducts were cleared manually. In our Fig. 2, node numbers are consensus parent masses, node size is linked to the relative molecular ion intensity based on peak area measured from the extracted ion chromatogram. Peak areas were measured automatically with the FreeStyle Genesis algorithm, sometimes modified manually if found unfitting, and then pasted manually into the Cytoscape table. This information was also used to create pie charts in which each portion represents the relative peak area of different isomers included in the same node (each GNPS node is a cluster of MS^2^ spectra that may come from different isomeric protonated molecular ions). For nodes gathering protonated molecular ions and in-source fragments of higher molecular weight compounds, only protonated molecular ion integrations were included for peak area information. Any ion present in the network solely due to in-source fragmentation was given the arbitrary extracted ion intensity of 0 and was white in the network. Furthermore, spectra of interest were manually annotated using different databases and tools, including Sirius^82^, Metfrag^83^ available at (https://msbi.ipb-halle.de/MetFrag/), Pubchem, Sci-Finder, and Mass Bank of North America (MoNa, https://mona.fiehnlab.ucdavis.edu/). Detailed spectral data are provided in supporting information (Figs. S1-S74, Supporting Information), and the raw spectral files are available (See Data Availability).

### Whole Genome Sequencing and Assembly of *Streptomyces sp*. RO-S4 strain

Genomic DNA was isolated from 50 mL of RO-S4 grown in M2 broth medium for 12 days at 28 °C with shaking (150 rpm/min). The DNA was extracted using the Bacteria Genomic DNA Extraction Kit (Promega, United States) according to the manufacturer’s instructions. Illumina whole genome sequencing was performed by the Genotoul facility in Toulouse, France. Briefly, libraries (Truseq nano HT, Ilumina) were constructed using 200 ng of purified DNA and sequenced on a Novaseq 6000 sequencer (Illumina), generating 93 million paired 150 base pair (bp) reads. The entire dataset was assembled using SPAdes (v3.14.0)^84^ with the option “careful”. The assembly was manually curated to remove contigs with low (<500) coverage and low (<55%) G+C content. Since a gene cluster of interest was truncated in a contig, we manually extended it using blast searches against raw reads and re-assembly of reads and of a downstream contig using the *gap4* tool of the Staden package (http://staden.sourceforge.net). This final assembly was auto-annotated using PROKKA v. 1.14.6^45^ using *Streptomyces* sp. Vc74B-19 protein descriptions and the annotation was manually curated prior to final submission to 1) add the ORF corresponding to GcnM in *Streptomyces* sp. LR32, 2) remove partial rRNA genes and possible adapters in contig ends. The genomic sequence was compared to that of close strains based on the 16S rRNA gene analysis described above, relatives (type strains) and genomes in the NCBI GenBank, based on top *blastp* hits using the five housekeeping genes described by Antony-Babu and coworkers^85^. ANI values were calculated using the OrthoANI tool available through the EZbiocloud server (https://www.ezbiocloud.net/tools/orthoani).

### Prediction of secondary metabolite biosynthetic gene clusters (BGCs) in the RO-S4 genome

The sequenced genome (DNA sequence as input) of the RO-S4 strain was analyzed for the prediction of secondary metabolites and biosynthetic gene clusters (BGCs) using the genome mining tool antiSMASH^86^, version 6.0.1. available through (https://antismash.secondarymetabolites.org) using the “relaxed” option. Synteny plots of the RO-S4 angucycline BGC was performed using Clinker v 0.0.21^87^. We also submitted the PROKKA annotated genbank formatted files using different annotation parameters. All results as well as descriptions of annotation pipelines are available (see data availability below).

### Phylogenetic analysis of UrdE and UrdM homologues

Phylogenetic analysis of UrdE and UrdM homologues in RO-S4 and in the BGCs of other angucyclines and the ring-opening Baeyer-Villiger monooxygenase MtmOIV in the mithramycin BGC was performed using SeaView Version 4.6 and Mega11^88^. Briefly, amino-acid sequences were aligned using muscle and a mask created using the gblocks options “allow smaller blocks positions” and “allow gap positions” followed by manual curation of the mask and a tree were generated by maximum likelihood with the GT+F model and gamma distribution using MEGA11 based on 359 homologous. A neighbor-joining tree was constructed using the JTT substitution model. The robustness of both trees was evaluated by bootstrap analysis using 100 replicates. The raw alignments, the regions used for reconstruction and results of model testing are available (see Data Availability).

## Acknowledgements

We thank the Bio2Mar platform (http://bio2mar.obs-banyuls.fr) for providing technical support and access to instrumentation. This work benefited from access to the Observatoire Océanologique de Banyuls-sur-Mer, an EMBRC-France and EMBRC-ERIC site.

## Author Contributions

R.O., D.S, and M.S. conceived and designed the experiments. R.O. and K.M. collected biological material. R.O., A.S.R., C.V. performed the experiments. D.S. and J.S. performed the metabolomics analysis. R.O and M.T.S the genomic analysis. R.O., D.S., and M.T.S. wrote the original draft and prepared tables and figures. All authors read and approved the final version.

## Additional Information

### Data availability

The draft genome sequence was deposited in the GenBank database under accession number JAJQKZ000000000. Raw Illumina reads have been deposited in the SRA under accession SRR17084181. The sequence analysis pipeline plus discussions regarding angucycline biosynthesis, AntiSMASH results and the files used for the phylogenetic analysis are available through are available through github (github.com/suzumar/ROS4_manus).

