## Supplemental Tables and Figures for "Integrated metabolomic, molecular networking, and genome mining analyses uncover novel angucyclines from *Streptomyces* sp. RO-S4 strain isolated from Bejaia Bay, Algeria"

**Table of content**

*Table S1. List and mass spectrometry annotation key of sugars constituting the angucycline ose moieties* 4
*Figure S1. MS spectrum of compound 1.* 5
*Figure S2. MS/MS spectrum of an ion at m/z 487.1600 [(1+H)$^+$].* 5
*Figure S3. Compared experimental MS/MS spectrum of ion at m/z 487.1600 [(1+H)$^+$] with: (a) MS$^2$ spectrum of protonated fridamycin and (b) MS$^2$ spectrum of protonated aquayamycin.* 6
*Figure S3. MS spectrum of compound 2.* 6
*Figure S4. MS/MS spectrum of ion at m/z 469.1496 [(2+H)$^+$].* 6
*Figure S5. MS spectrum of compound 3.* 7
*Figure S6. MS/MS spectrum of ion at m/z 529.1708 [(3+H)$^+$].* 7
*Figure S7. MS spectrum of compound 4.* 7
*Figure S8. MS/MS spectrum of ion at m/z 557.2378 [(4+H)$^+$].* 7
*Figure S9. MS spectrum of compound 5.* 8
*Figure S10. MS/MS spectrum of ion at m/z 669.2903 [(5+H)$^+$].* 8
*Figure S11. MS spectrum of compound 6a.* 8
*Figure S12. MS/MS spectrum of ion at m/z 597.1973 [(6a+H)$^+$].* 8
*Figure S13. MS spectrum of compound 6b.* 9
*Figure S14. MS/MS spectrum of ion at m/z 597.1967 [(6b+H)$^+$].* 9
*Figure S15. MS spectrum of compound 7a.* 9
*Figure S16. MS/MS spectrum of ion at m/z 711.2650 [(7a+H)$^+$].* 9
*Figure S17. MS spectrum of compound 7b.* 10
*Figure S18. MS/MS spectrum of ion at m/z 711.2656 [(7b+H)$^+$].* 10
*Figure S19. MS spectrum of compound 8.* 10
*Figure S20. MS/MS spectrum of ion at m/z 545.1808 [(8+H)$^+$].* 10
*Figure S21. MS spectrum of compound 9.* 11
*Figure S22. MS/MS spectrum of ion at m/z 667.2751 [(9+H)$^+$].* 11
*Figure S23. MS spectrum of compound 10.* 11
*Figure S24. MS/MS spectrum of ion at m/z 781.3436 [(10+H)$^+$].* 11

*Figure S25. MS spectrum of compound 11a.* ____ *12*
*Figure S26. MS/MS spectrum of ion at m/z 675.2438 [(11a+H)$^+$].* ____ *12*
*Figure S27. MS spectrum of compound 11b.* ____ *12*
*Figure S28. MS/MS spectrum of ion at m/z 675.2436 [(11b+H)$^+$].* ____ *12*
*Figure S29. MS spectrum of compound 12.* ____ *13*
*Figure S30. MS/MS spectrum of ion at m/z 677.2597 [(12+H)$^+$].* ____ *13*
*Figure S31. MS spectrum of compound 13.* ____ *13*
*Figure S32. MS/MS spectrum of ion at m/z 763.3328 [(13+H)$^+$].* ____ *13*
*Figure S33. MS spectrum of compound 14.* ____ *14*
*Figure S34. MS/MS spectrum of ion at m/z 917.3601 [(14+H)$^+$].* ____ *14*
*Figure S35. MS spectrum of compound 15a.* ____ *14*
*Figure S36. MS/MS spectrum of ion at m/z 819.2863 [(15a+H)+].* ____ *14*
*Figure S37. MS spectrum of compound 15b.* ____ *15*
*Figure S38. MS/MS spectrum of ion at m/z 819.2869 [(15b+H)$^+$].* ____ *15*
*Figure S39. MS spectrum of compound 16.* ____ *15*
*Figure S40. MS/MS spectrum of ion at m/z 901.3653 [(16+H)$^+$].* ____ *15*
*Figure S41. MS spectrum of compound 17.* ____ *16*
*Figure S42. MS/MS spectrum of ion at m/z 952.3960 [(17+$NH_4$)$^+$].* ____ *16*
*Figure S43. MS/MS spectrum of ion at m/z 957.3506 [(17+Na)$^+$].* ____ *16*
*Figure S44. MS spectrum of compound 18a.* ____ *16*
*Figure S45. MS/MS spectrum of ion at m/z 599.2142 [(18a+H)$^+$].* ____ *17*
*Figure S46. MS spectrum of compound 18b.* ____ *17*
*Figure S47. MS/MS spectrum of ion at m/z 599.2125 [(18b+H)$^+$].* ____ *17*
*Figure S48. MS spectrum of compound 19a.* ____ *17*
*Figure S49. MS/MS spectrum of ion at m/z 601.2284 [(19a+H)$^+$].* ____ *18*
*Figure S50. MS spectrum of compound 19b.* ____ *18*
*Figure S51. MS/MS spectrum of ion at m/z 601.2288 [(19b+H)$^+$].* ____ *18*
*Figure S52. MS spectrum of compound 20.* ____ *18*
*Figure S53. MS/MS spectrum of ion at m/z 583.2170 [(20+H)$^+$].* ____ *19*
*Figure S54. MS spectrum of compound 21.* ____ *19*
*Figure S55. MS/MS spectrum of ion at m/z 697.2861 [(21+H)$^+$].* ____ *19*
*Figure S56. MS spectrum of compound 22.* ____ *19*
*Figure S57. MS/MS spectrum of ion at m/z 585.2333 [(22+H)$^+$].* ____ *20*
*Figure S58. MS spectrum of compound 23.* ____ *20*
*Figure S59. MS/MS spectrum of ion at m/z 713.2782 [(23+H)$^+$].* ____ *20*
*Figure S60. MS spectrum of compound 24.* ____ *20*
*Figure S61. MS/MS spectrum of ion at m/z 672.2655 [(24+H)$^+$]* ____ *21*
*Figure S62. MS spectrum of compound 25.* ____ *21*
*Figure S63. MS/MS spectrum of ion at m/z 743.3031 [(25+H)$^+$].* ____ *21*
*Figure S64. Fragmentation scheme for compound 25 from [M+H]$^+$ to the aglycone ion.* ____ *21*
*Figure S65. MS spectrum of compound 26.* ____ *22*
*Figure S66. MS/MS spectrum of ion at m/z 813.3816 [(26+H)$^+$].* ____ *22*
*Figure S67. MS spectrum of compound 27.* ____ *22*
*Figure S68. MS/MS spectrum of ion at m/z 857.3709 [(27+H)$^+$].* ____ *22*
*Figure S69. MS spectrum of compound 28.* ____ *23*
*Figure S70. MS/MS spectrum of ion at m/z 709.2490 [(28+H)$^+$].* ____ *23*
*Figure S71. MS spectrum of compound 29.* ____ *23*
*Figure S72. MS/MS spectrum of ion at m/z 595.1808 [(29+H)$^+$].* ____ *23*
*Figure S73. MS spectrum of compound 30.* ____ *24*
*Figure S74. MS/MS spectrum of ion at m/z 705.2182 [(30+H)$^+$].* ____ *24*

*Table S2. Summary of antiSMASH 6.0 secondary metabolites predicted from the RO-S4 strain* ______ *25*
*Table S3. Deduced functions of the ORFs in the ROS4_2 cluster 'Streptomyces sp. RO-S4 strain and their BLASTs. We have considered only the sequences that showed a homology >50%* ______ *26*
*Figure S75. tRNA$_{ala}$ predicted by Aragorn in the complementary strand of the LRR80_00492 ORF region.* ______ *34*
*Figure S76. RNA-fold predicted RNA secondary structure of the region predicted by Aragorn to code (in the complementary strand) a tRNAala in the LRR80_00492 ORF region gene.* ______ *35*
*Figure S77. Phylogenetic reconstruction (Neighbor Joining) of FAD dependent oxidases as described in the text. Where:* ______ *37*
*Figure S78. Phylogenetic reconstruction (Maximum Likelihood) of FAD dependent oxidases as described in the text.* ______ *39*

**Table S1.** List and mass spectrometry annotation key of sugars constituting the angucycline ose moieties

| Name | Neutral loss 1 | Neutral loss 2 | Oxonium cation |
|---|---|---|---|
| **Aculose** | 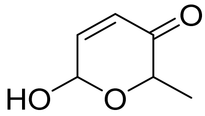 Exact Mass: 128.0473 | 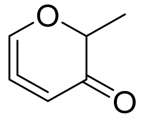 Exact Mass: 110.0368 | 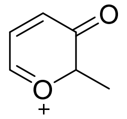 Exact Mass: 111.0441 |
| **Oxidized aculose ($C_6H_8O_4$)** | 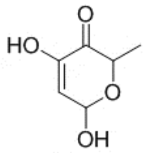 Chemical Formula: $C_6H_8O_4$ Exact Mass: 144.0423 | 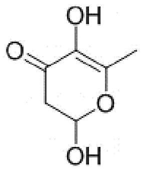 Chemical Formula: $C_6H_8O_4$ Exact Mass: 144.0423 | 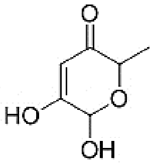 Chemical Formula: $C_6H_8O_4$ Exact Mass: 144.0423 |
| **Rhodinose or amicetose (diastereoisomers)** | 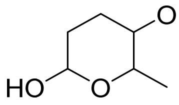 Exact Mass: 132.0786 | 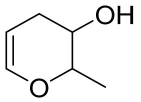 Exact Mass: 114.0681 | 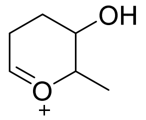 Exact Mass: 115.0754 |
| **Cinerulose A** | 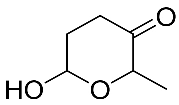 Exact Mass: 130.0630 | 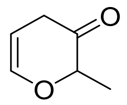 Exact Mass: 112.0524 | 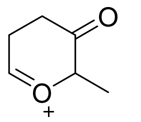 Exact Mass: 113.0597 |
| **Cinerulose B** | 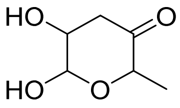 Exact Mass: 146.0579 | 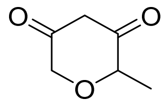 Exact Mass: 128.0473 | 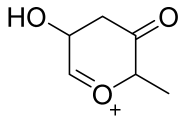 Exact Mass: 129.0546 |
| **Olivose** | 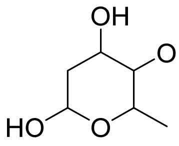 Exact Mass: 148.0736 | 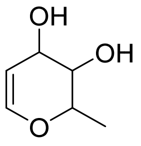 Exact Mass: 130.0630 | 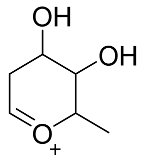 Exact Mass: 131.0703 |
| **Kerriose** | 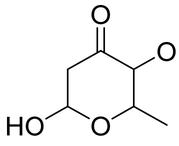 Exact Mass: 146.0579 | 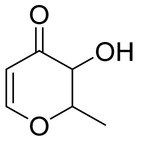 Exact Mass: 128.0473 | 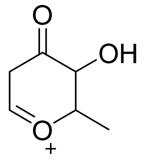 Exact Mass: 129.0546 |

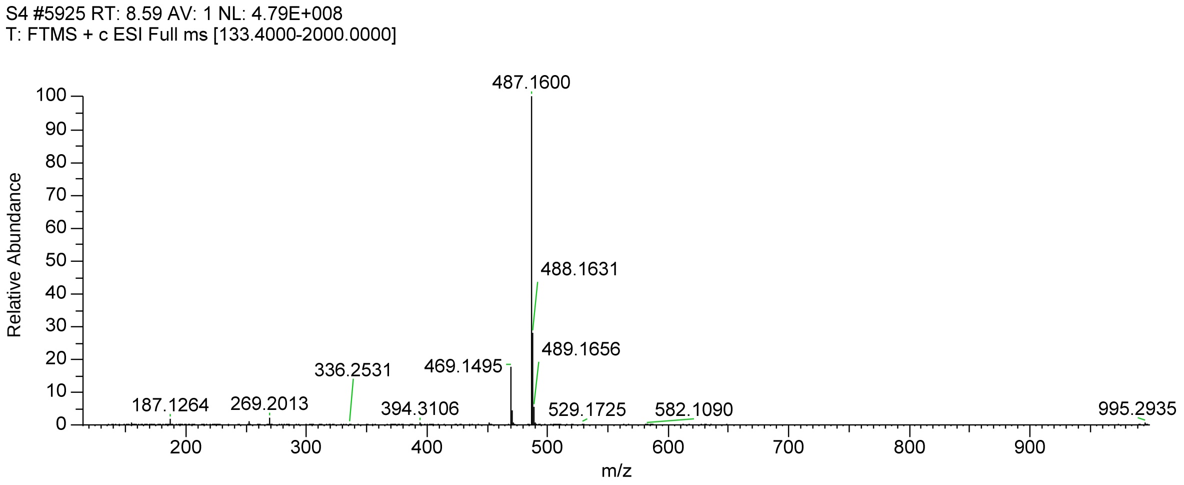


**Figure S1.** MS spectrum of compound **1.**

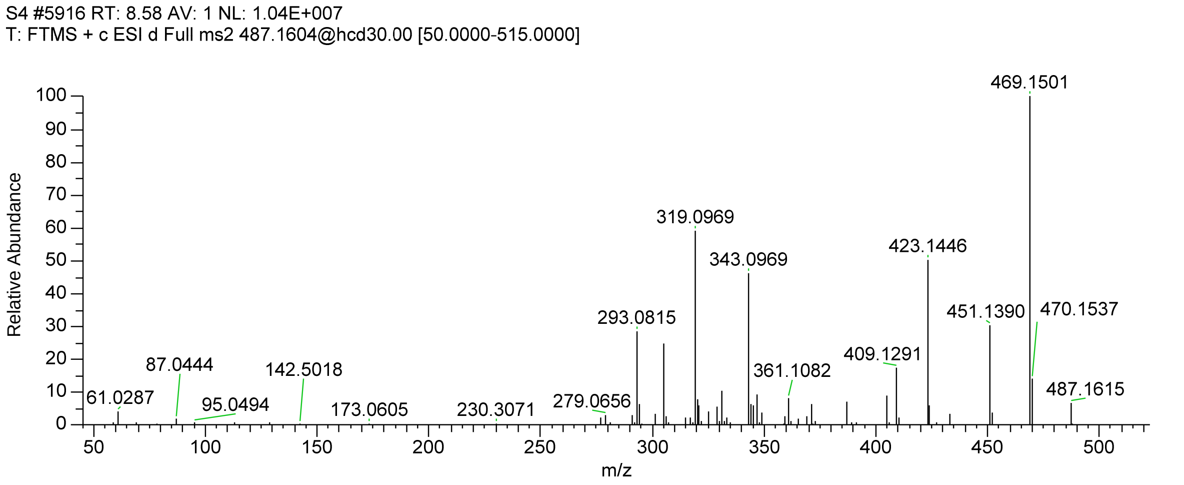


**Figure S2.** MS/MS spectrum of an ion at *m/z* 487.1600 [(**1**+H)$^+$].

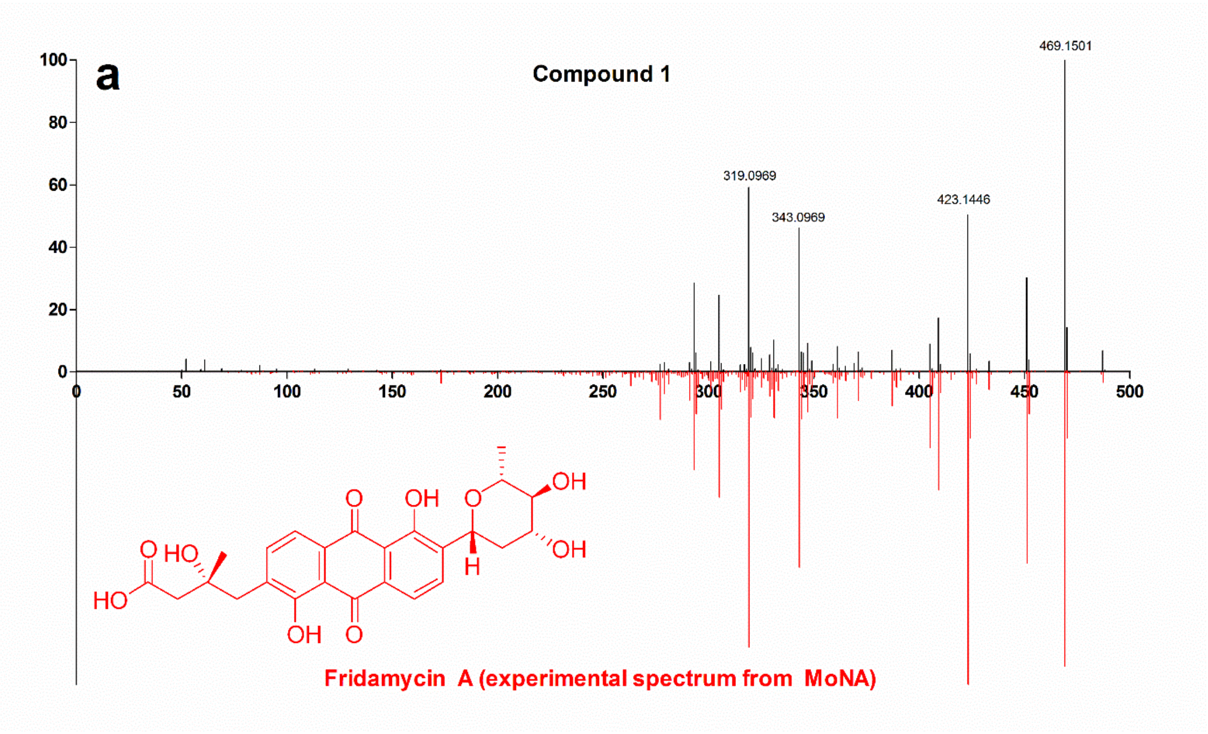

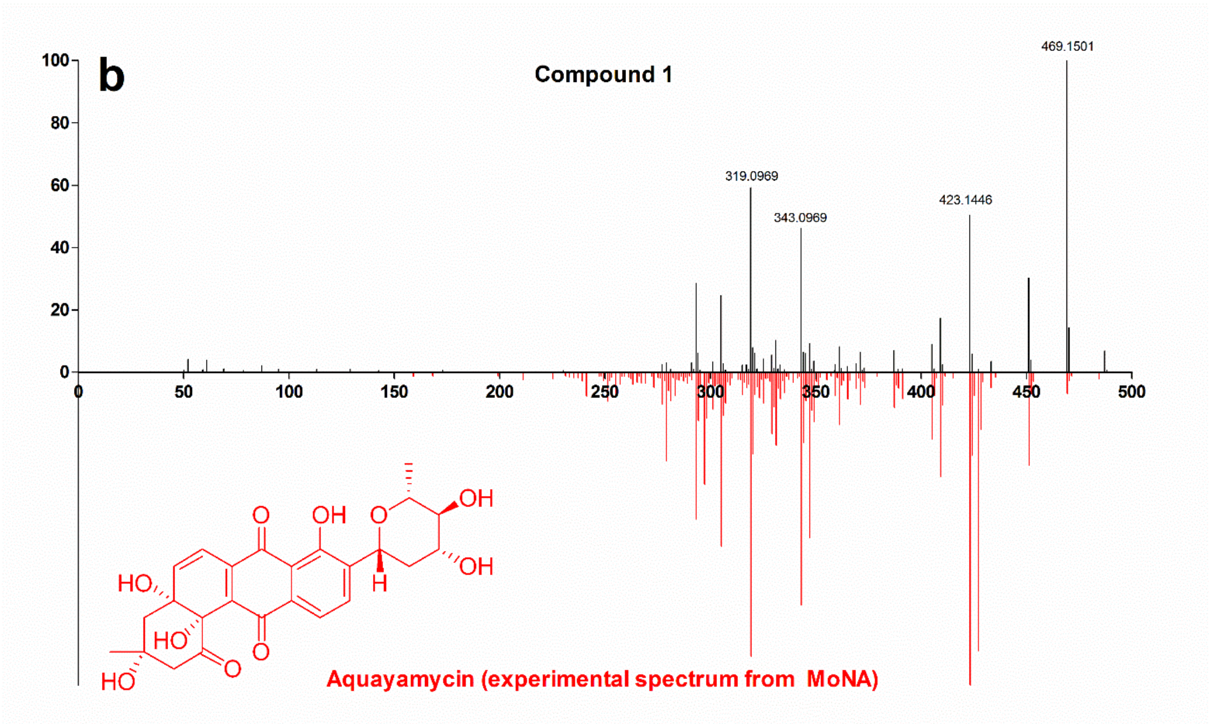


**Figure S3.** Compared experimental MS/MS spectrum of ion at *m/z* 487.1600 [(**1**+H)$^+$] with: (a) MS$^2$ spectrum of protonated fridamycin and (b) MS$^2$ spectrum of protonated aquayamycin.

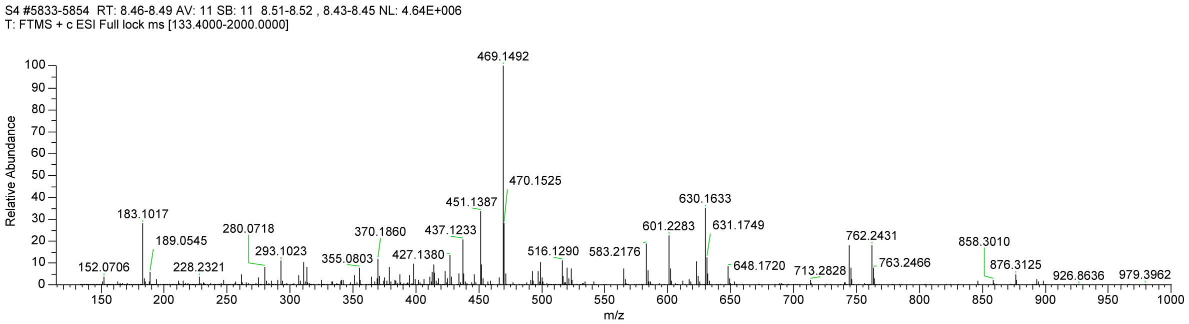


**Figure S3.** MS spectrum of compound **2**.

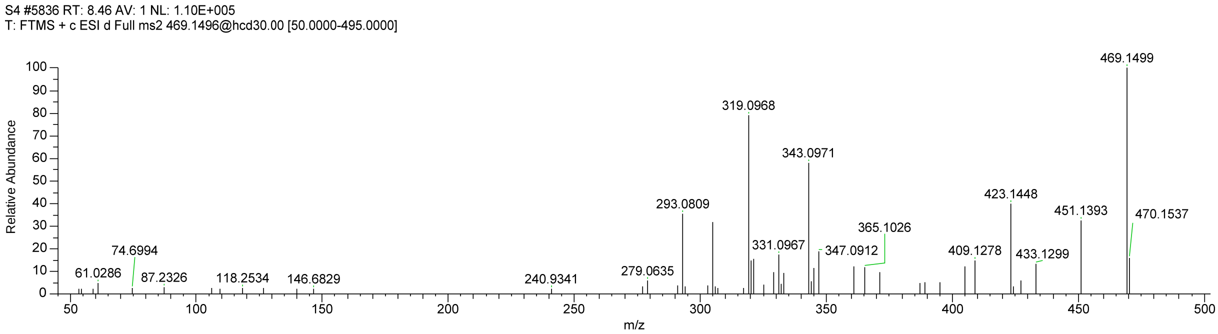


**Figure S4.** MS/MS spectrum of ion at *m/z* 469.1496 [(**2**+H)$^+$].

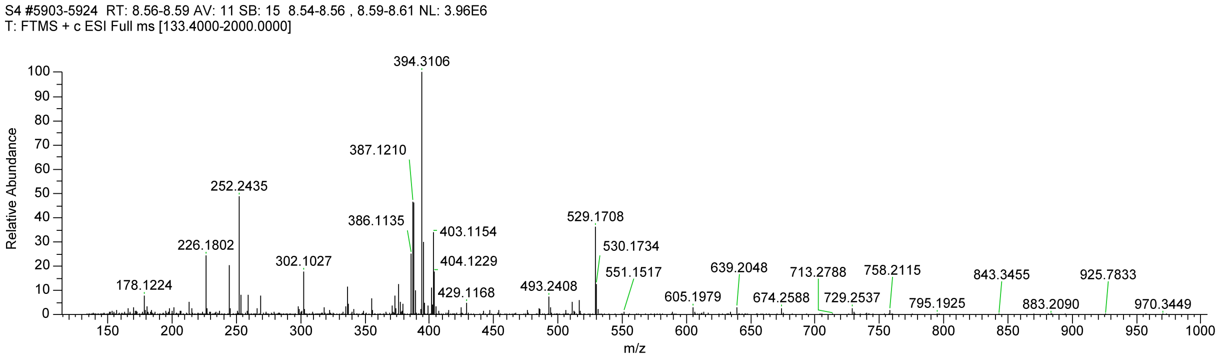


**Figure S5.** MS spectrum of compound **3**.

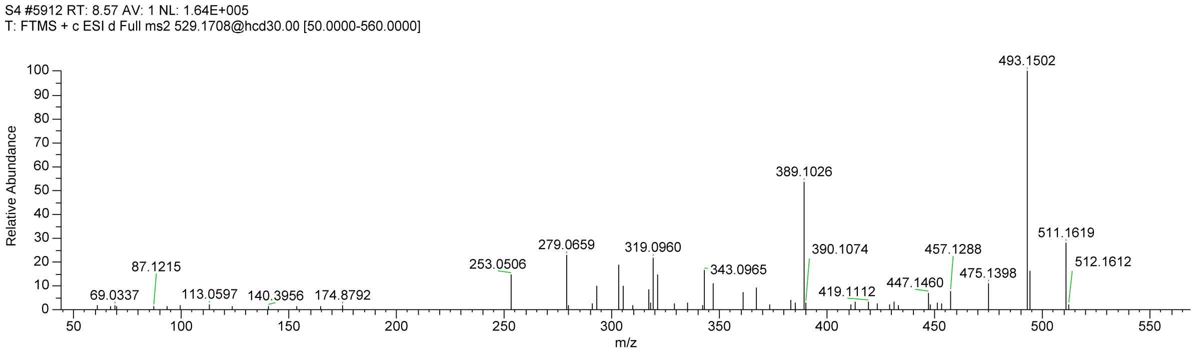


**Figure S6.** MS/MS spectrum of ion at *m/z* 529.1708 [(**3**+H)$^+$].

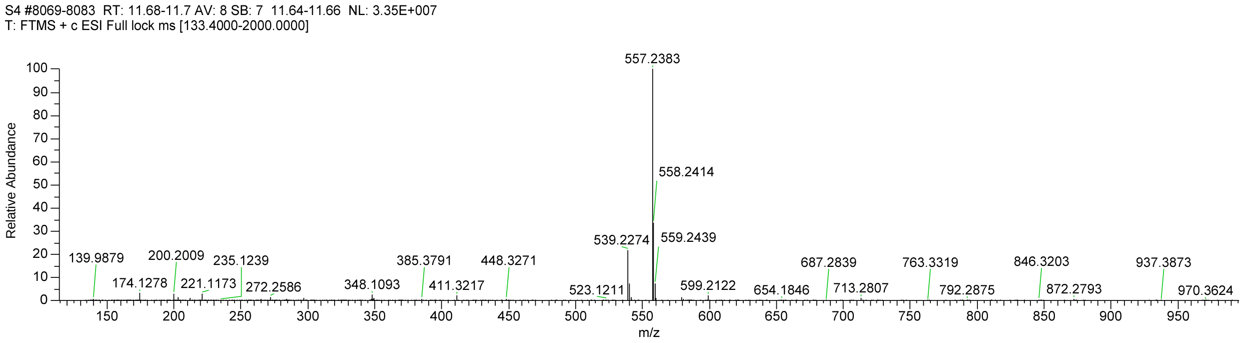


**Figure S7.** MS spectrum of compound **4**.

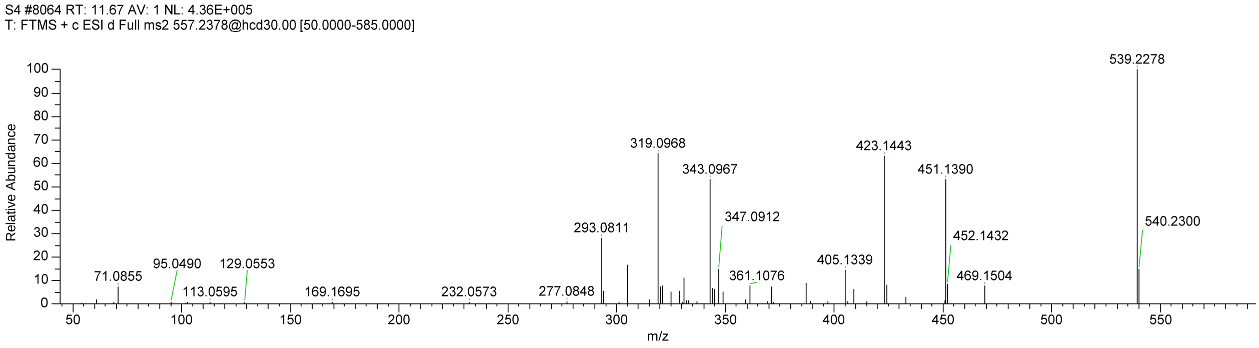


**Figure S8.** MS/MS spectrum of ion at *m/z* 557.2378 [(**4**+H)$^+$].

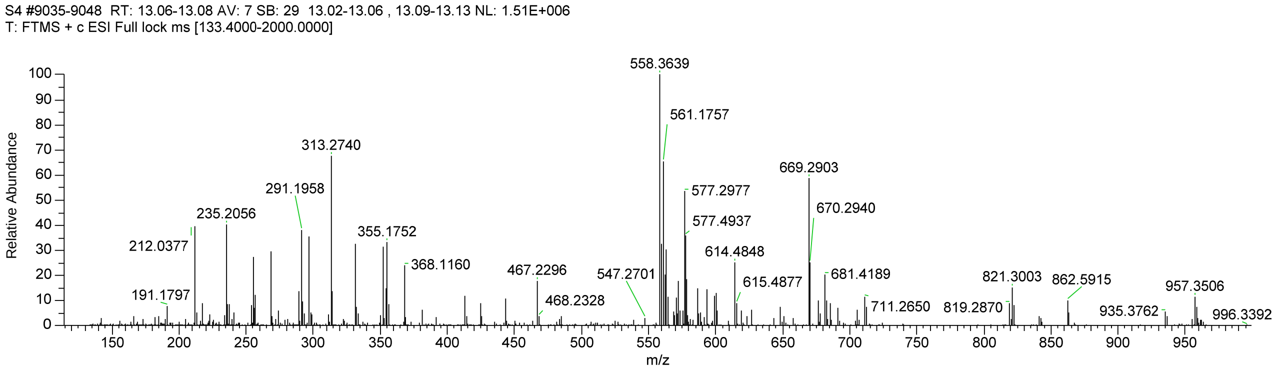


**Figure S9.** MS spectrum of compound **5**.

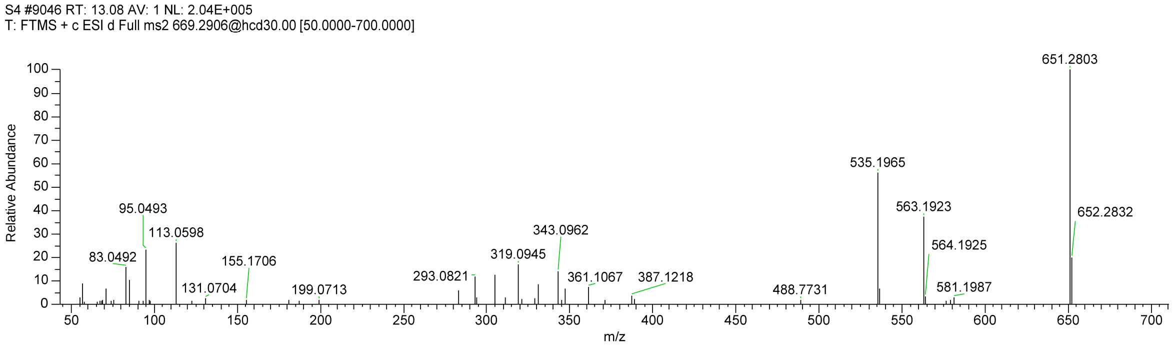


**Figure S10.** MS/MS spectrum of ion at *m/z* 669.2903 [(**5**+H)$^{+}$].

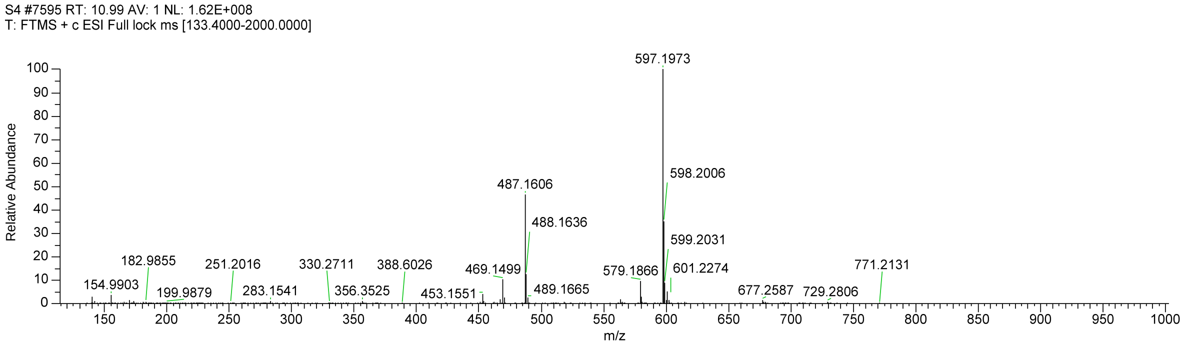


**Figure S11.** MS spectrum of compound **6a**.

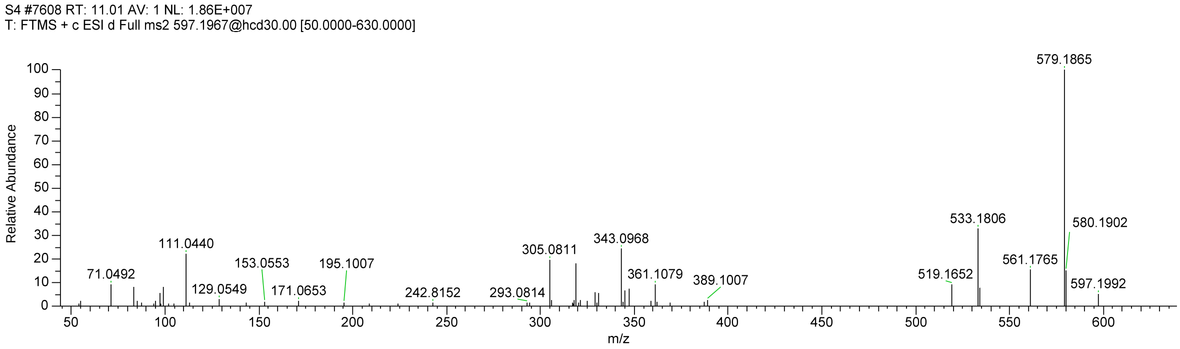


**Figure S12.** MS/MS spectrum of ion at *m/z* 597.1973 [(**6a**+H)$^{+}$].

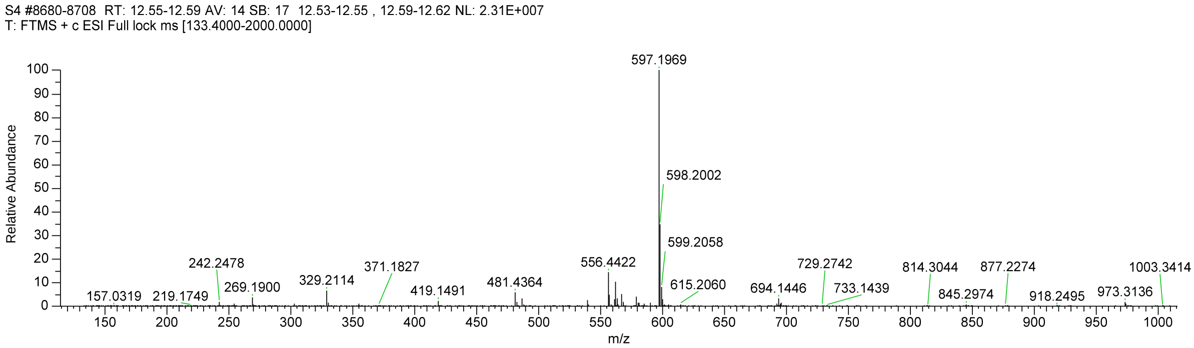


**Figure S13.** MS spectrum of compound **6b**.

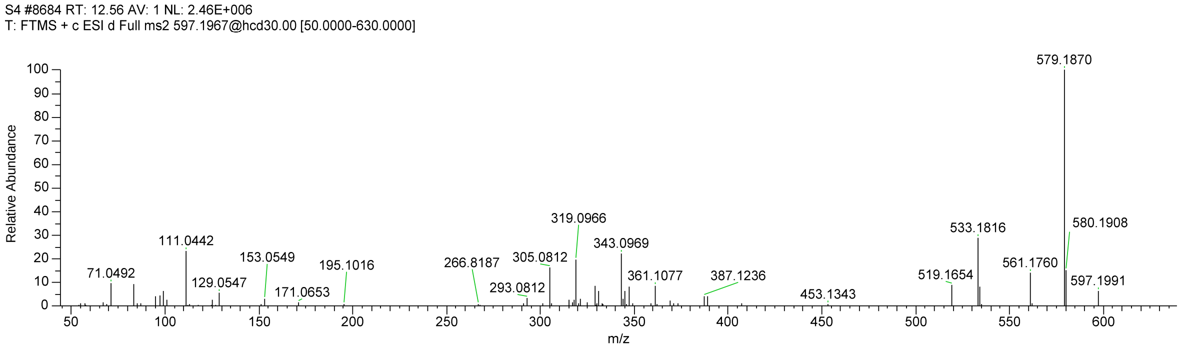


**Figure S14.** MS/MS spectrum of ion at *m/z* 597.1967 [(**6b**+H)$^{+}$].

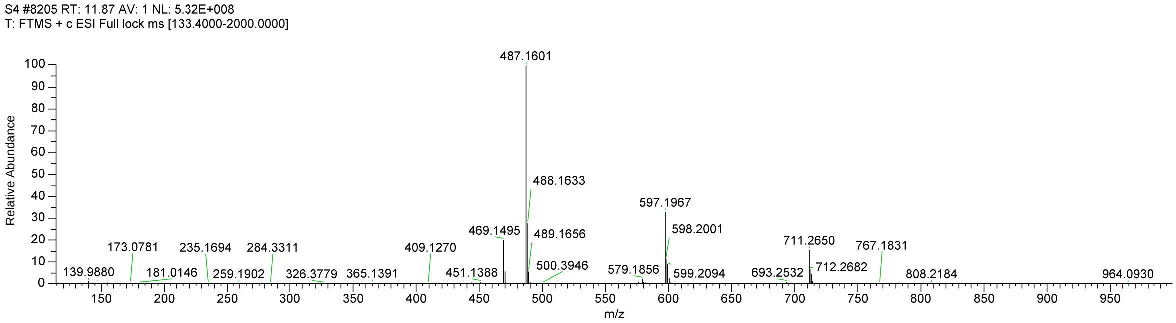


**Figure S15.** MS spectrum of compound **7a**.

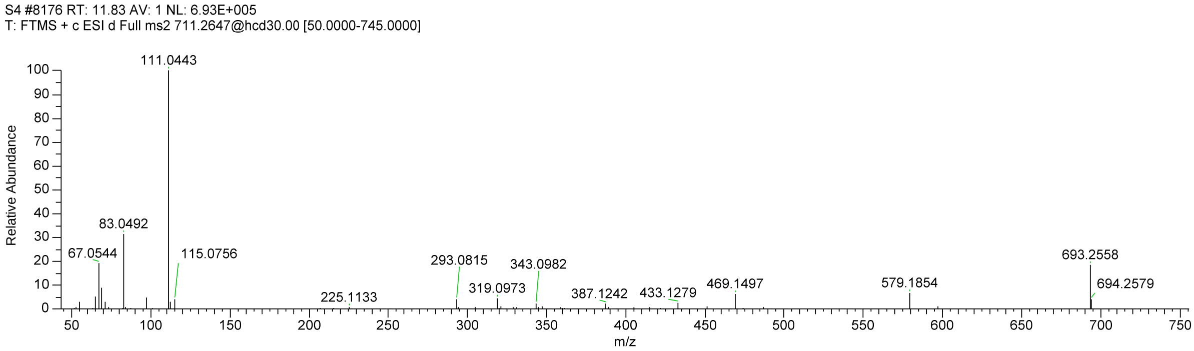


**Figure S16.** MS/MS spectrum of ion at *m/z* 711.2650 [(**7a**+H)$^{+}$].

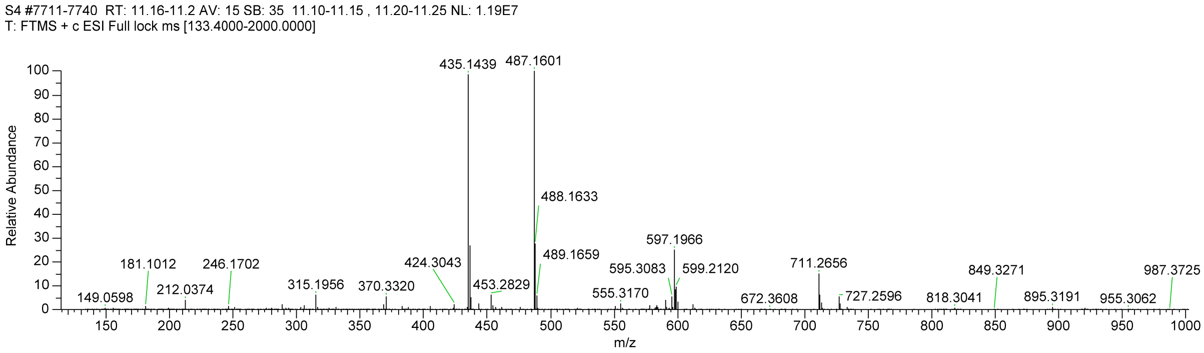


**Figure S17.** MS spectrum of compound **7b**.

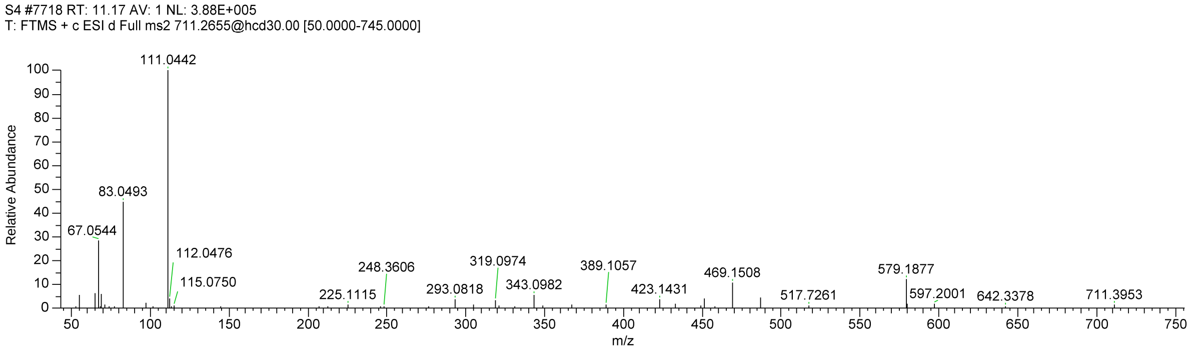


**Figure S18.** MS/MS spectrum of ion at *m/z* 711.2656 [(**7b**+H)$^+$].

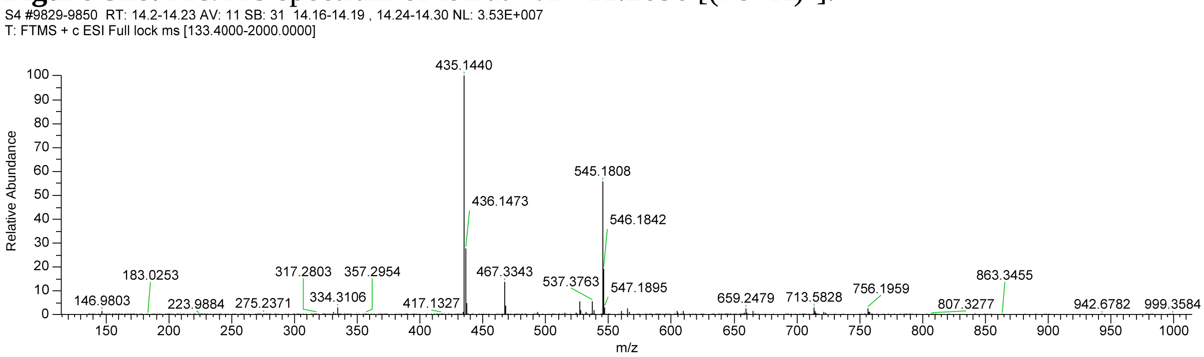


**Figure S19.** MS spectrum of compound **8**.

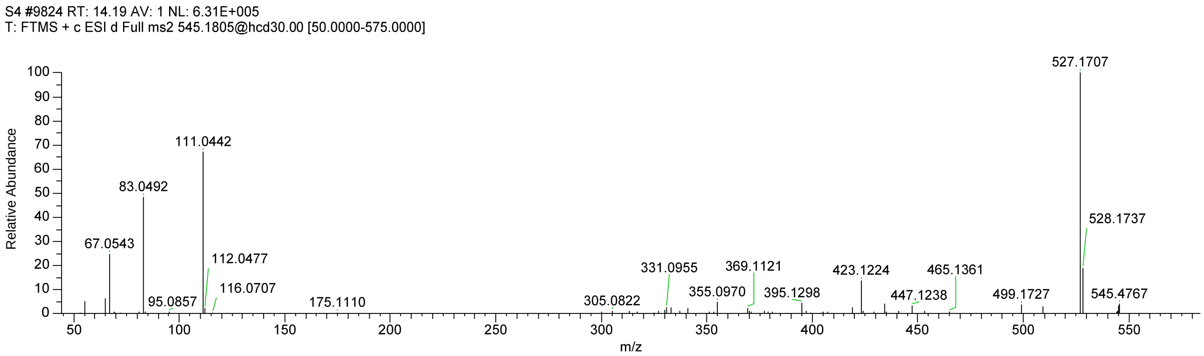


**Figure S20.** MS/MS spectrum of ion at *m/z* 545.1808 [(**8**+H)$^+$].

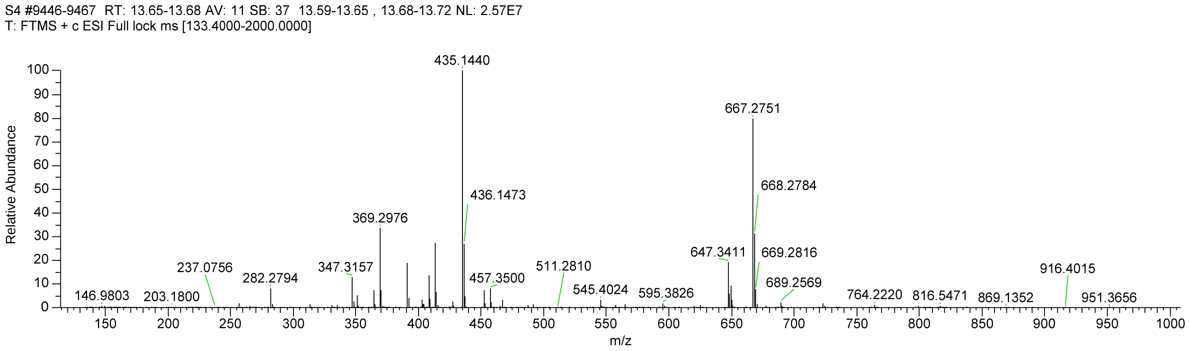


**Figure S21.** MS spectrum of compound **9**.

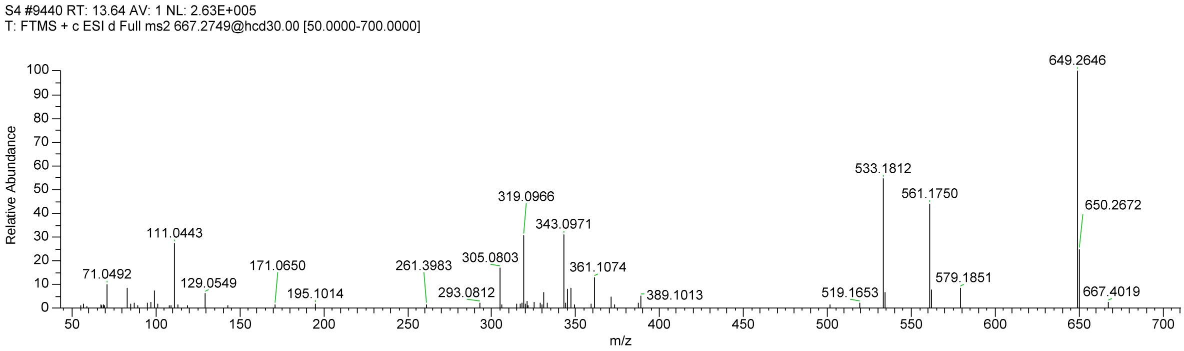


**Figure S22.** MS/MS spectrum of ion at *m/z* 667.2751 [(**9**+H)$^+$].

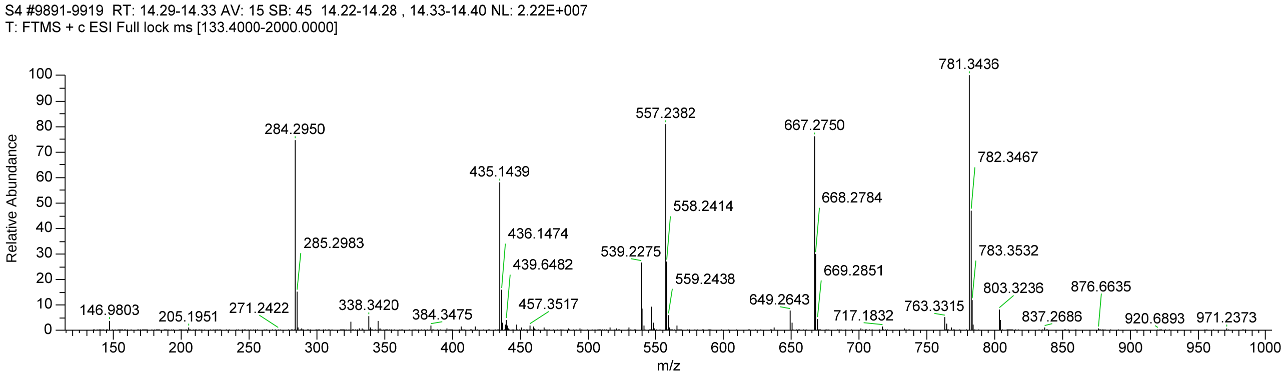


**Figure S23.** MS spectrum of compound **10**.

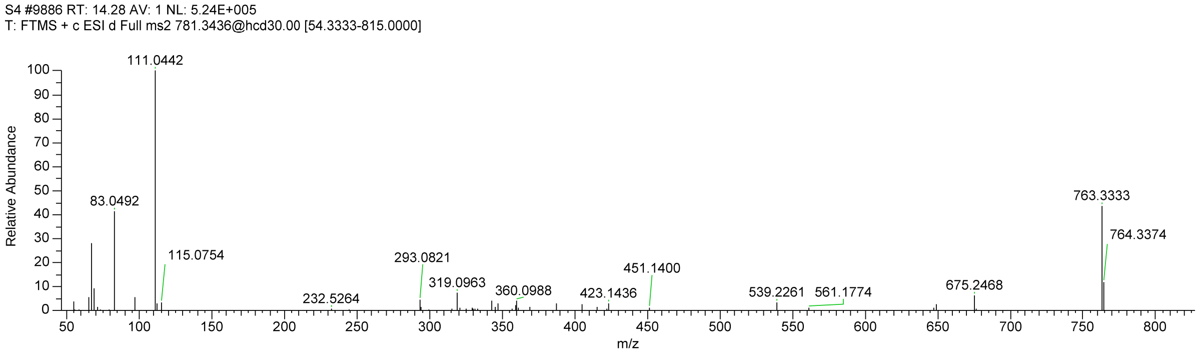


**Figure S24.** MS/MS spectrum of ion at *m/z* 781.3436 [(**10**+H)$^+$].

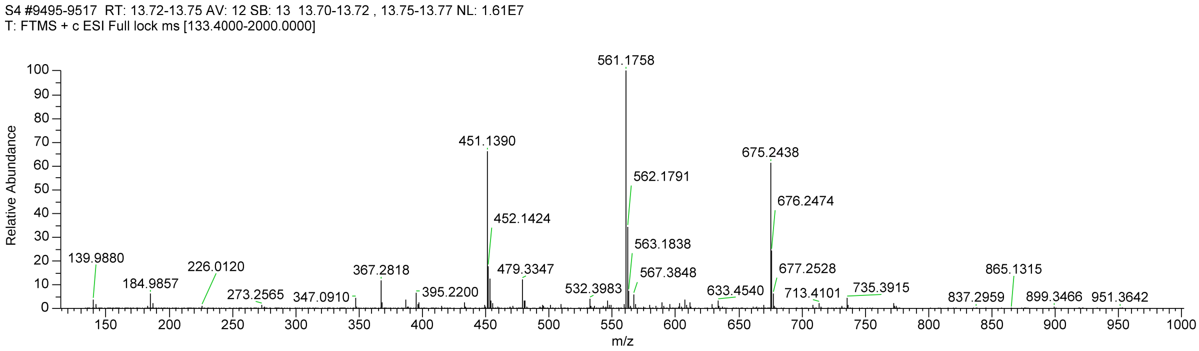


**Figure S25.** MS spectrum of compound **11a**.

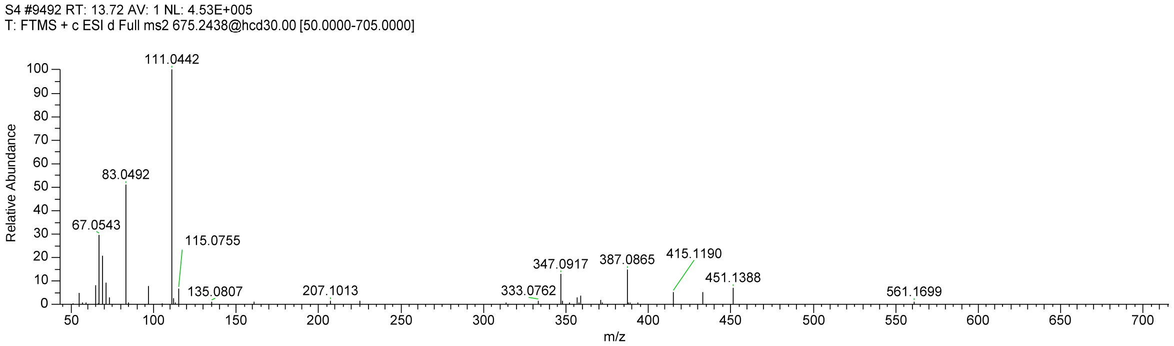


**Figure S26.** MS/MS spectrum of ion at *m/z* 675.2438 [(**11a**+H)$^+$].

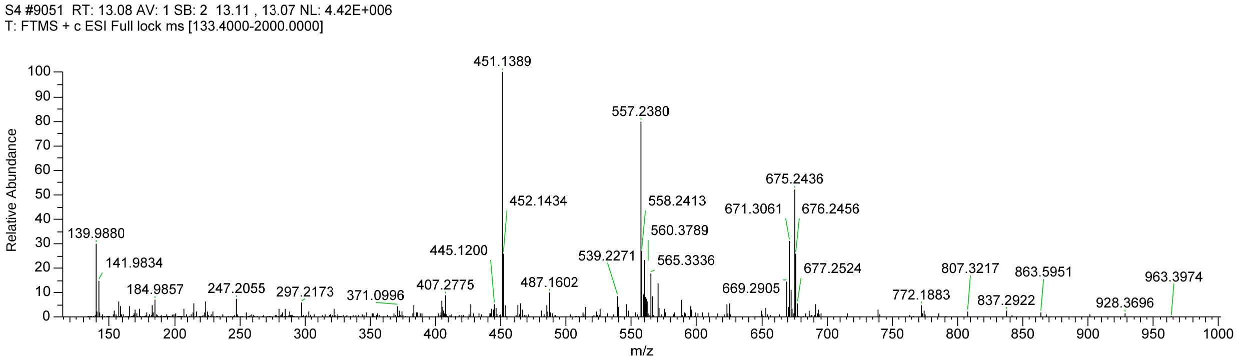


**Figure S27.** MS spectrum of compound **11b**.

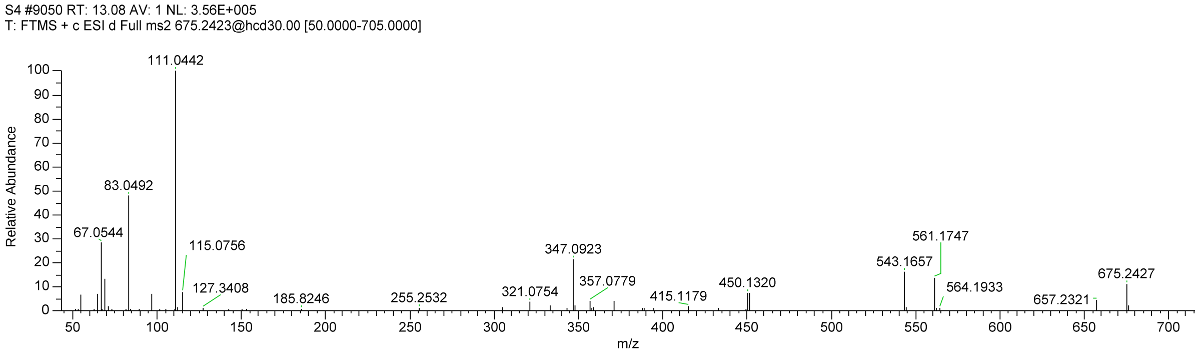


**Figure S28.** MS/MS spectrum of ion at *m/z* 675.2436 [(**11b**+H)$^+$].

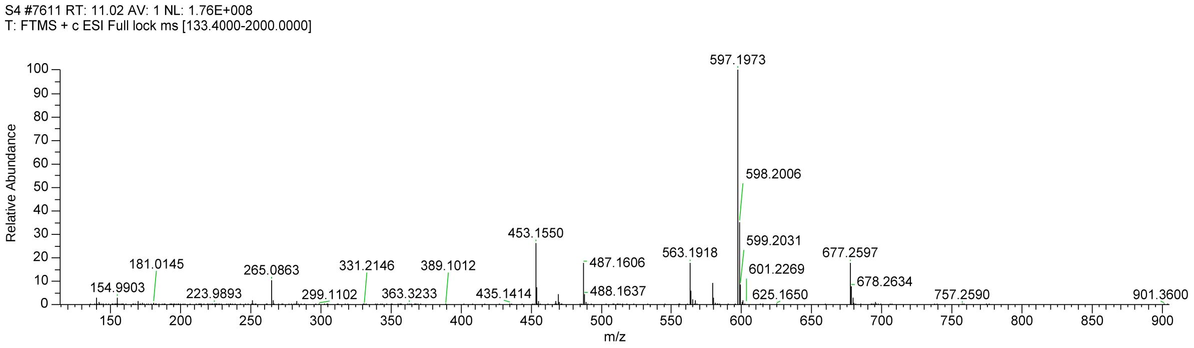


**Figure S29.** MS spectrum of compound **12**.

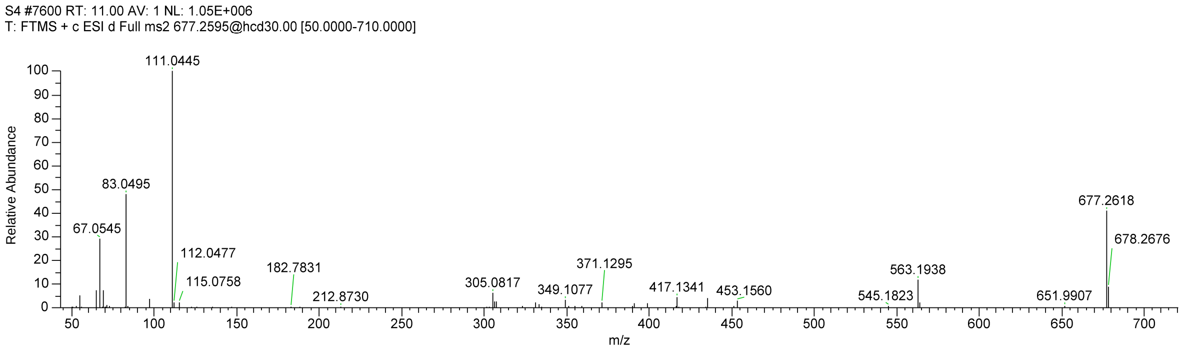


**Figure S30.** MS/MS spectrum of ion at *m/z* 677.2597 [(**12**+H)$^{+}$].

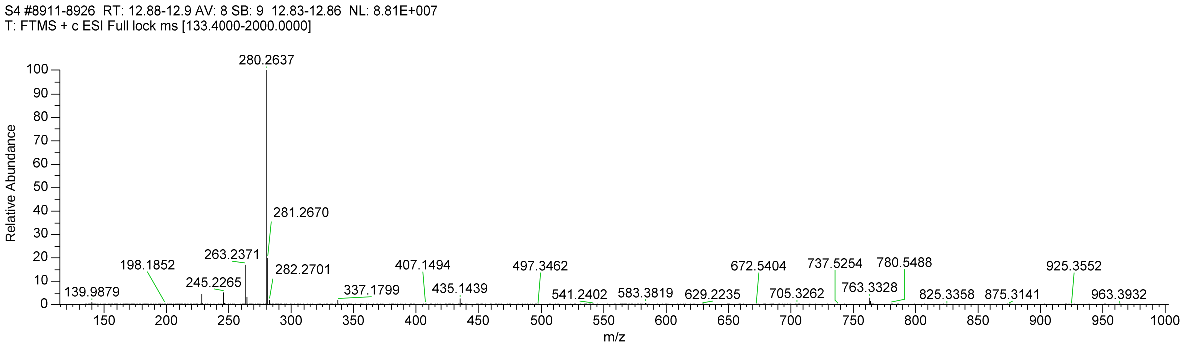


**Figure S31.** MS spectrum of compound **13**.

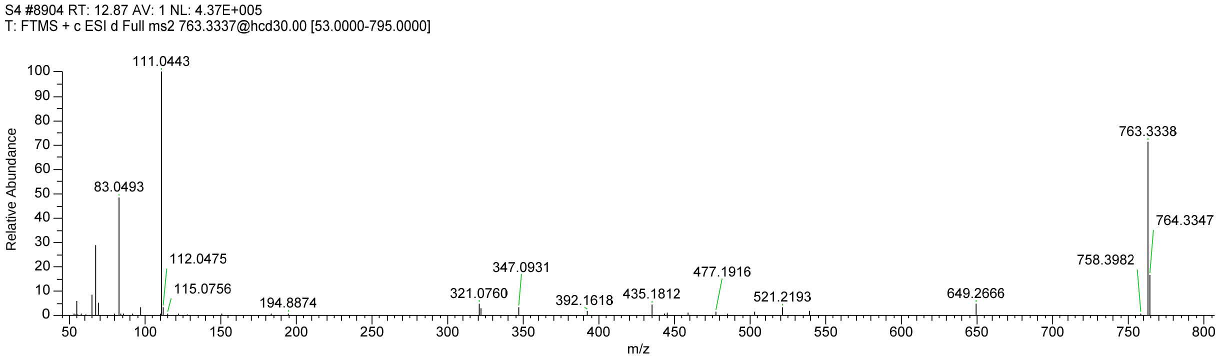


**Figure S32.** MS/MS spectrum of ion at *m/z* 763.3328 [(**13**+H)$^{+}$].

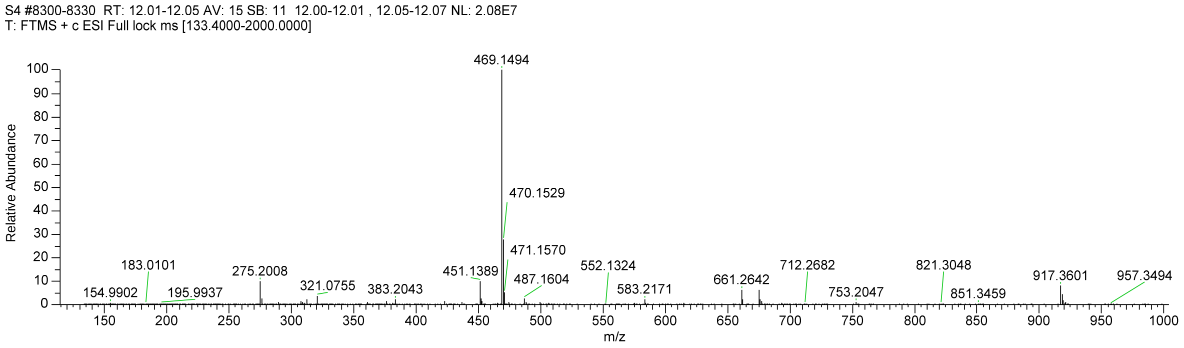


**Figure S33.** MS spectrum of compound **14**.

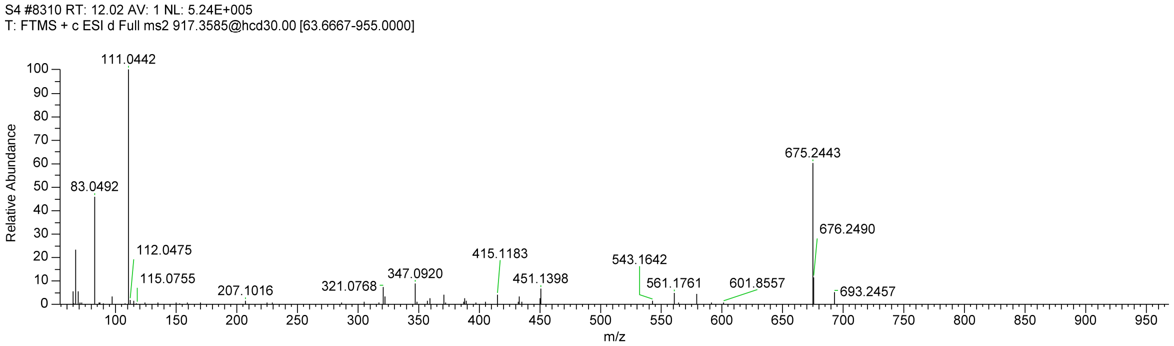


**Figure S34.** MS/MS spectrum of ion at *m/z* 917.3601 [(**14**+H)$^{+}$].

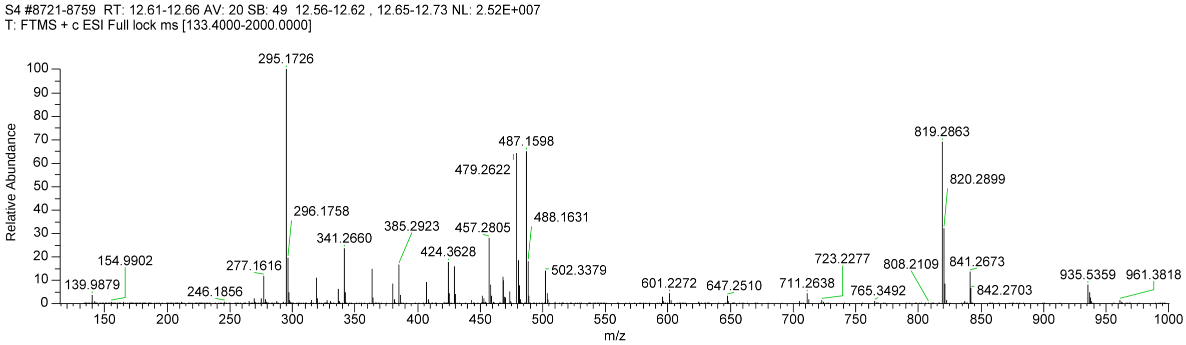


**Figure S35.** MS spectrum of compound **15a**.

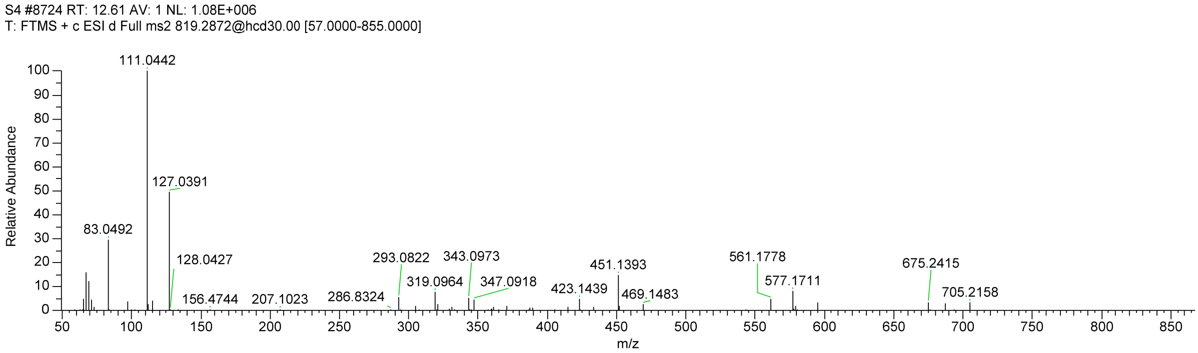


**Figure S36.** MS/MS spectrum of ion at m/z 819.2863 [(**15a**+H)+].

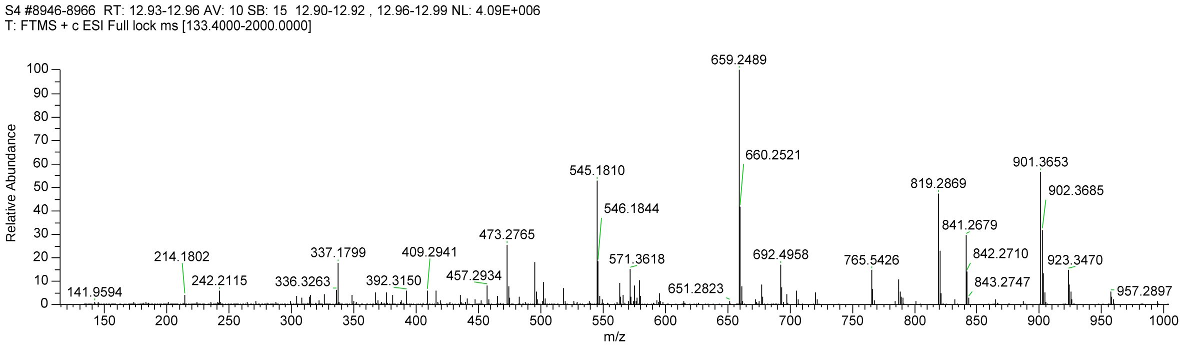


**Figure S37.** MS spectrum of compound **15b**.

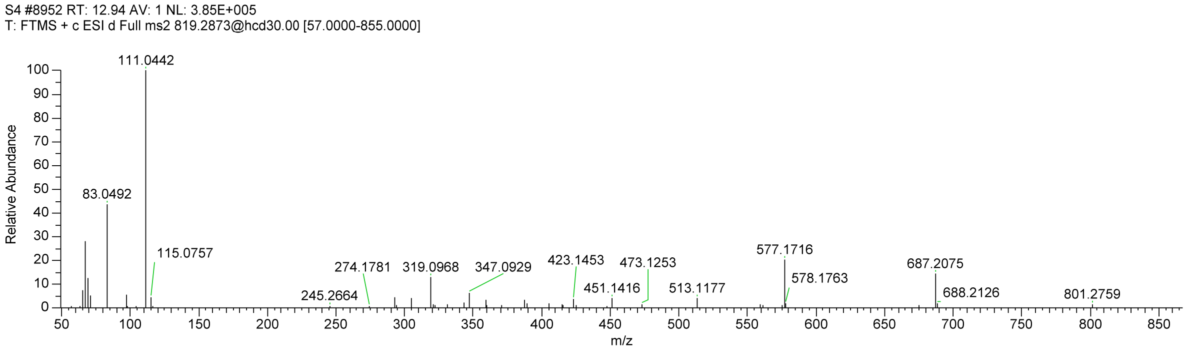


**Figure S38.** MS/MS spectrum of ion at *m/z* 819.2869 [(**15b**+H)$^+$].

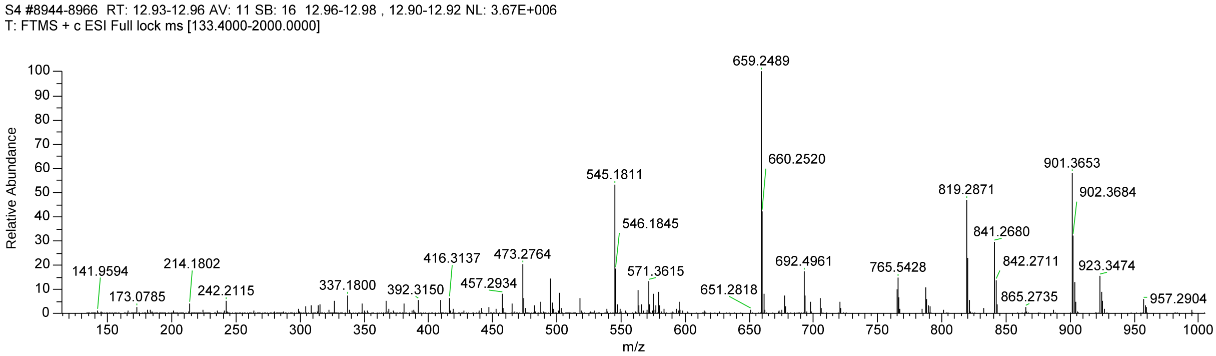


**Figure S39.** MS spectrum of compound **16**.

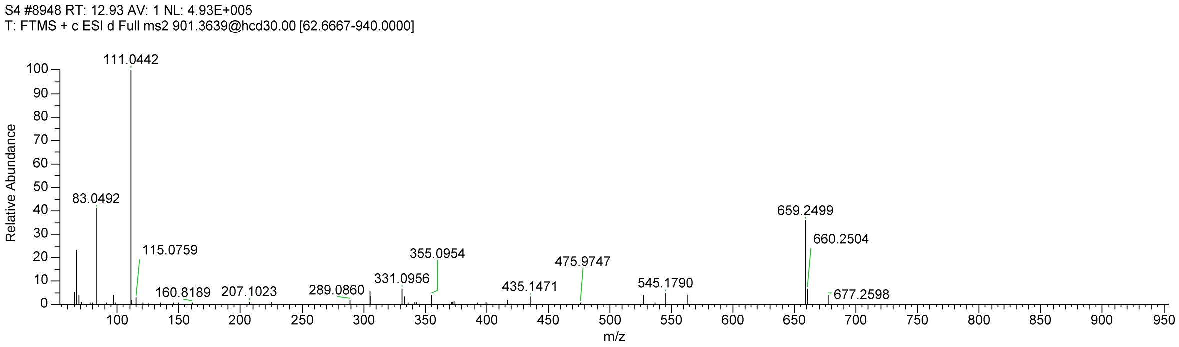


**Figure S40.** MS/MS spectrum of ion at *m/z* 901.3653 [(**16**+H)$^+$].

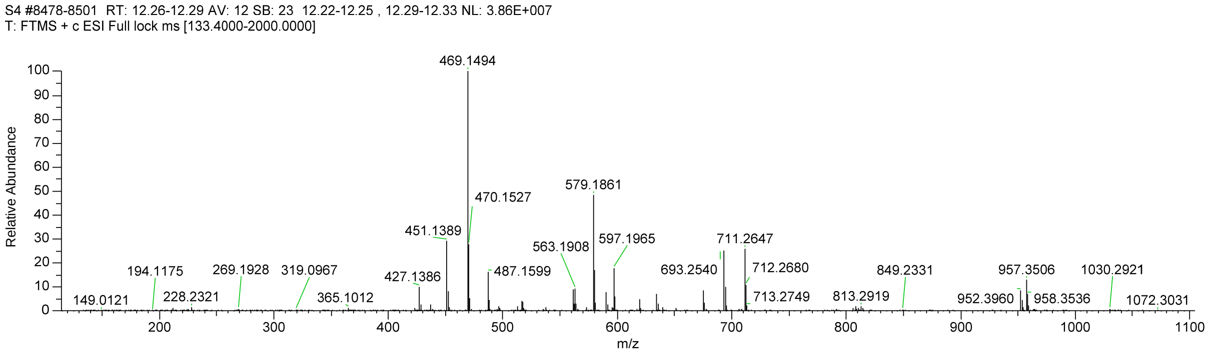


**Figure S41.** MS spectrum of compound **17**.

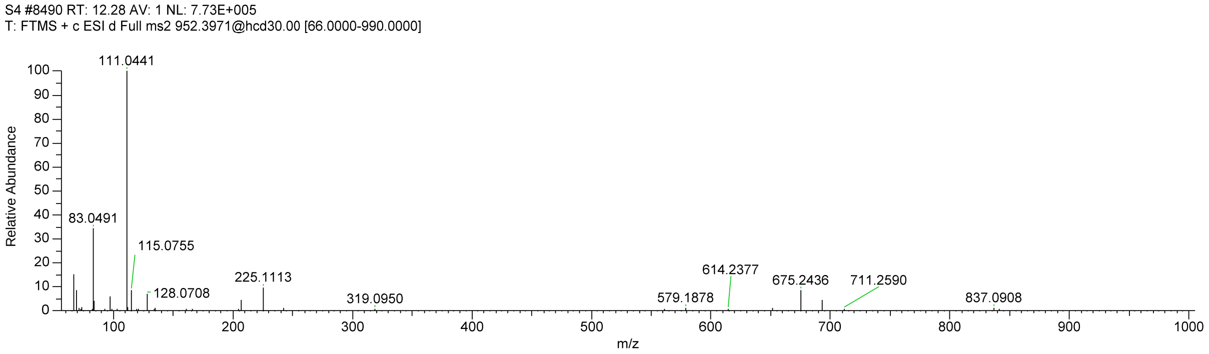


**Figure S42.** MS/MS spectrum of ion at *m/z* 952.3960 [(**17**+$NH_4$)$^+$].

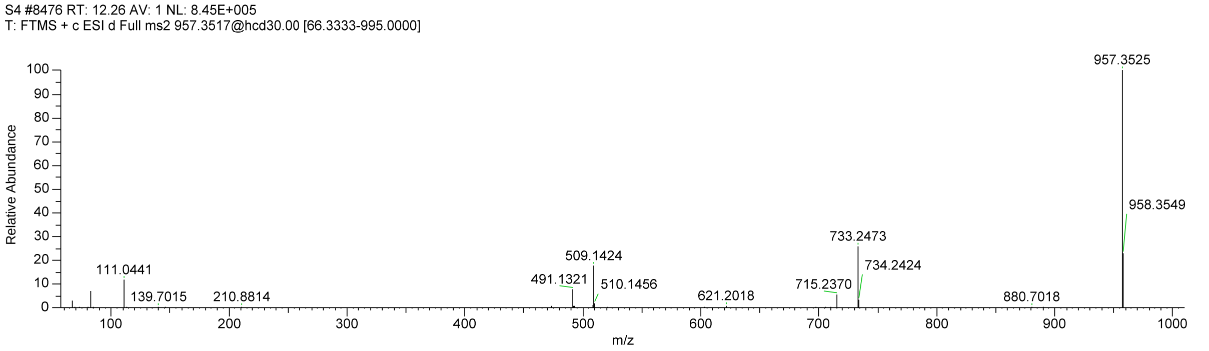


**Figure S43.** MS/MS spectrum of ion at *m/z* 957.3506 [(**17**+Na)$^+$].

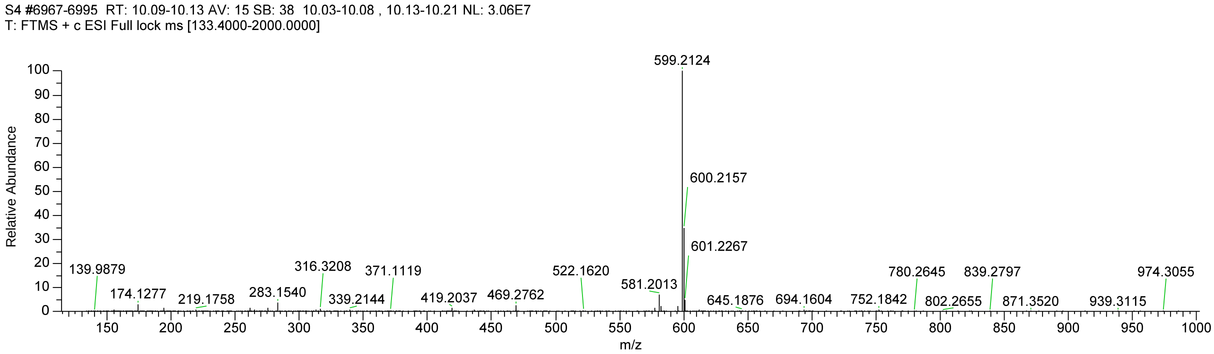


**Figure S44.** MS spectrum of compound **18a**.

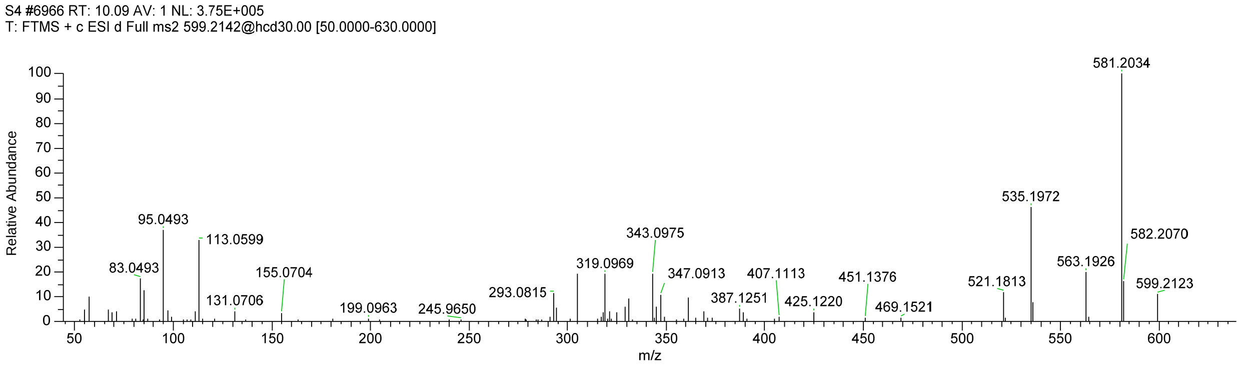


**Figure S45.** MS/MS spectrum of ion at *m/z* 599.2142 [(**18a**+H)$^+$].

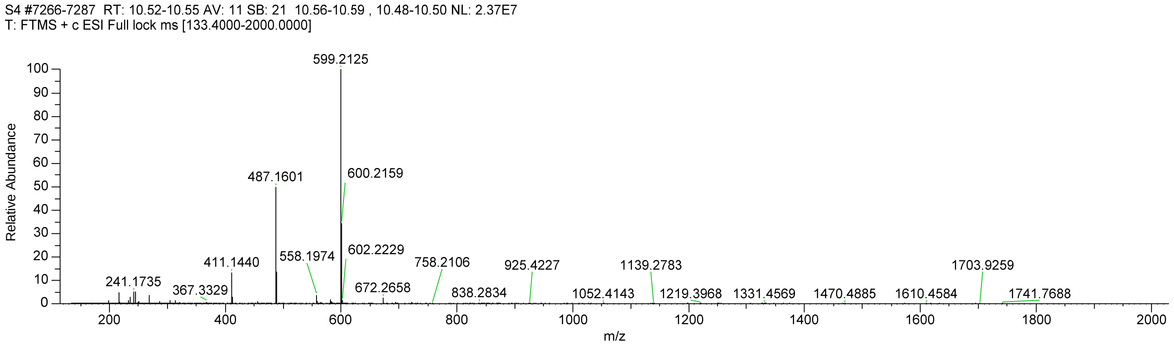


**Figure S46.** MS spectrum of compound **18b**.

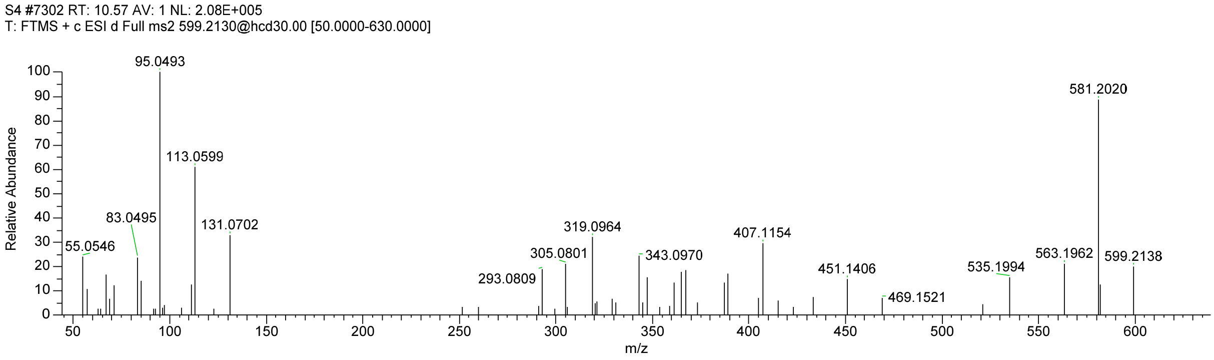


**Figure S47.** MS/MS spectrum of ion at *m/z* 599.2125 [(**18b**+H)$^+$].

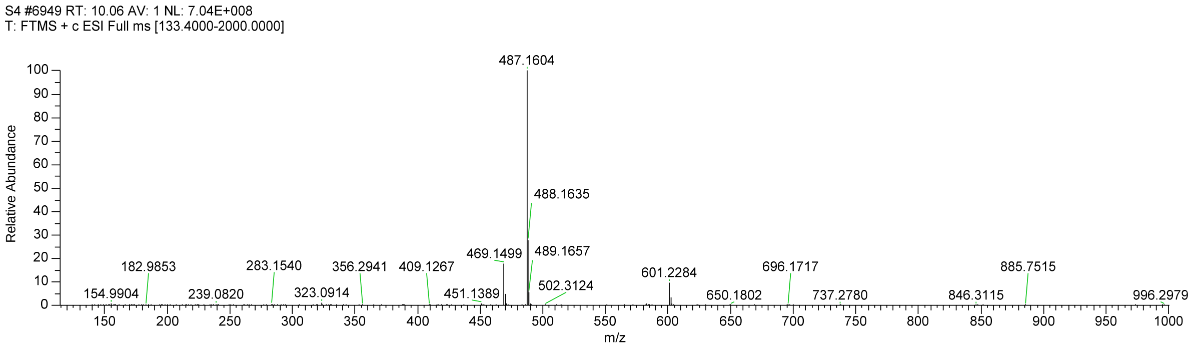


**Figure S48.** MS spectrum of compound **19a**.

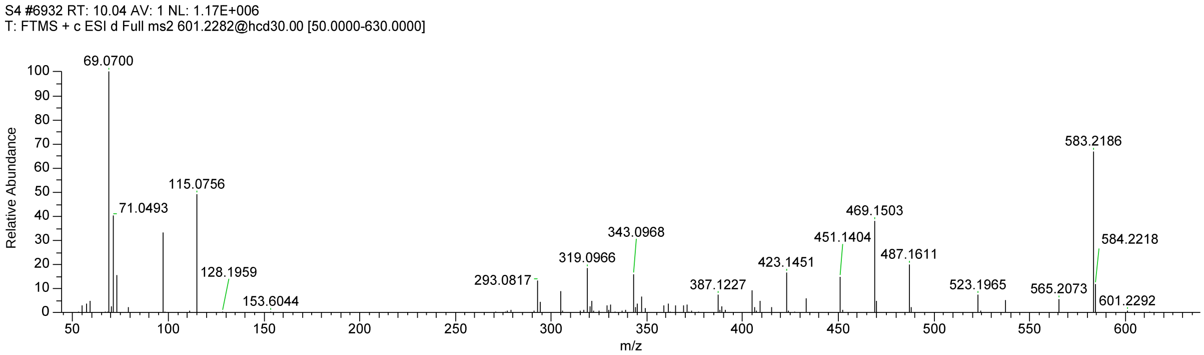


**Figure S49.** MS/MS spectrum of ion at *m/z* 601.2284 [(**19a**+H)$^+$].

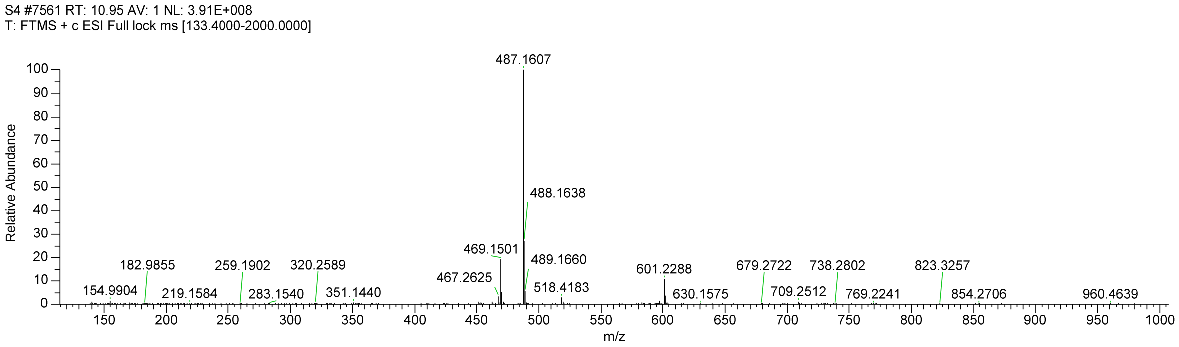


**Figure S50.** MS spectrum of compound **19b**.

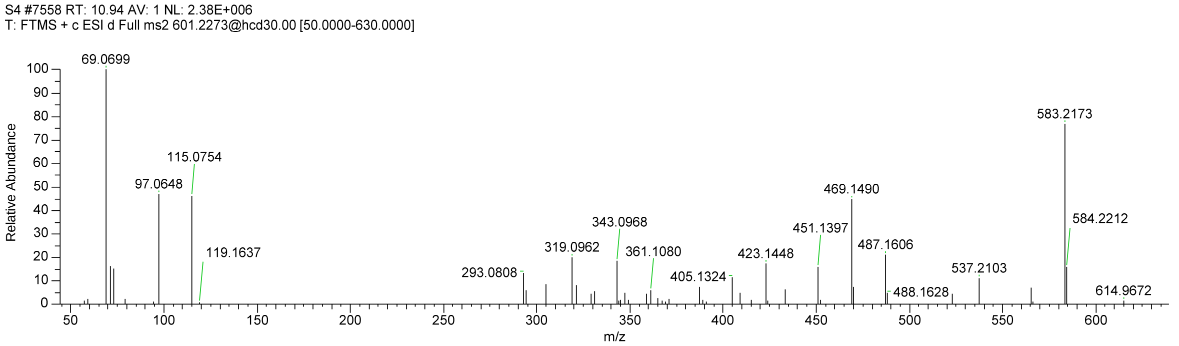


**Figure S51.** MS/MS spectrum of ion at *m/z* 601.2288 [(**19b**+H)$^+$].

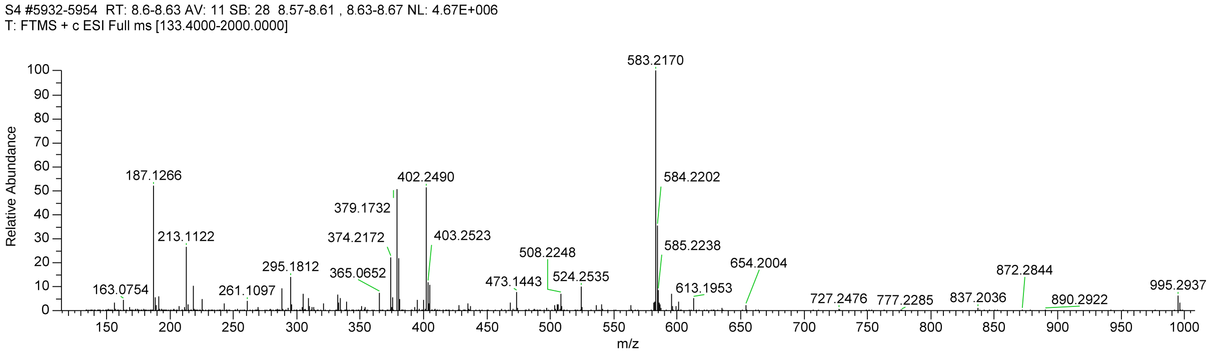


**Figure S52.** MS spectrum of compound **20**.

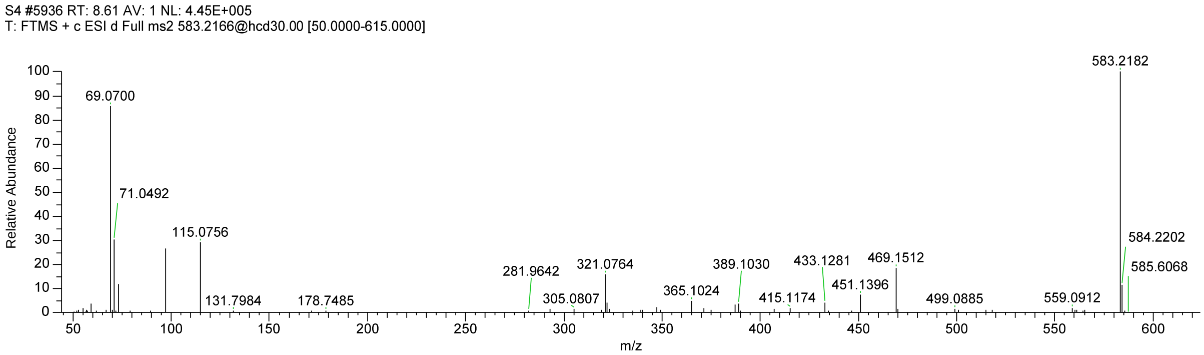


**Figure S53.** MS/MS spectrum of ion at *m/z* 583.2170 [(**20**+H)$^+$].

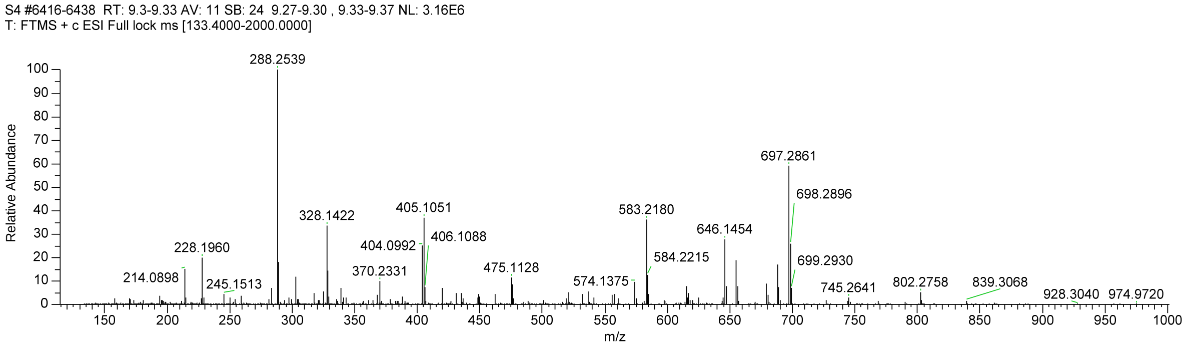


**Figure S54.** MS spectrum of compound **21**.

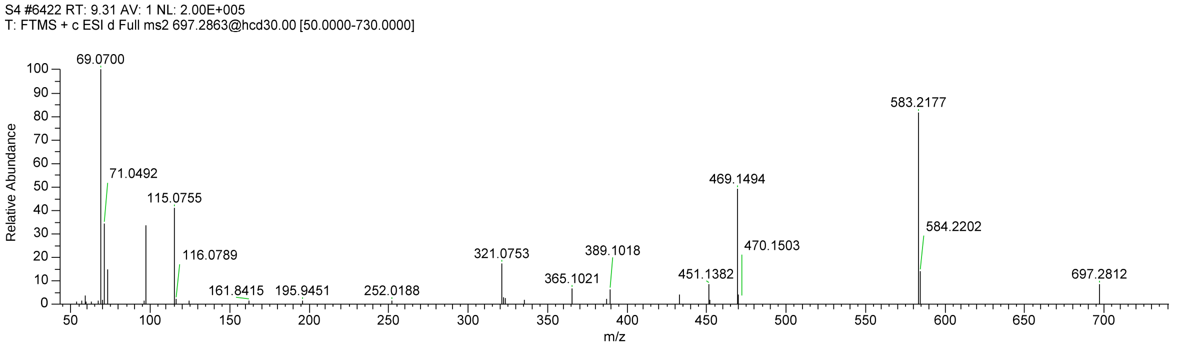


**Figure S55.** MS/MS spectrum of ion at *m/z* 697.2861 [(**21**+H)$^+$].

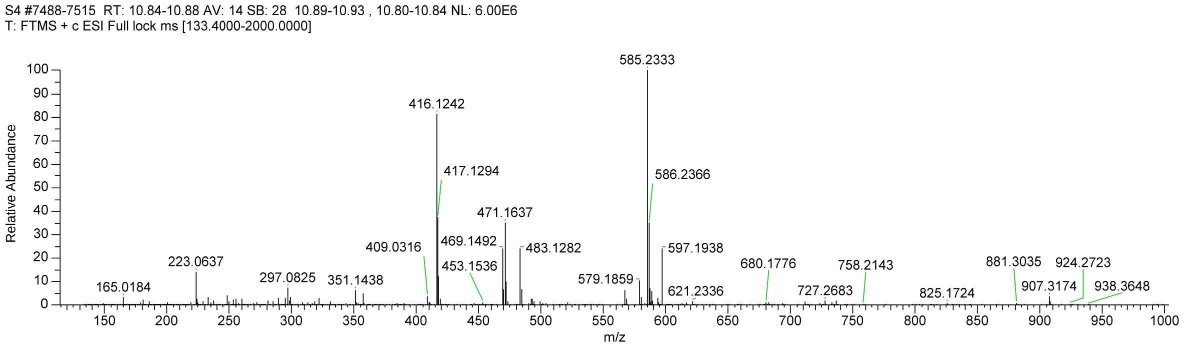


**Figure S56.** MS spectrum of compound **22**.

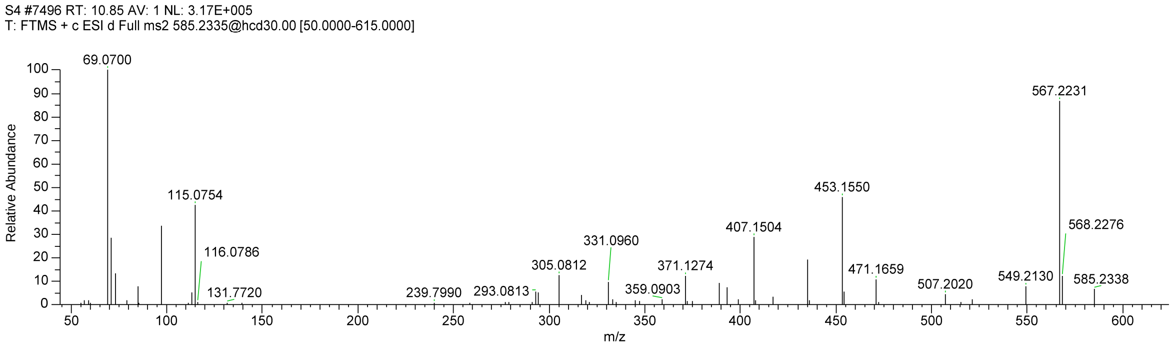


**Figure S57.** MS/MS spectrum of ion at *m/z* 585.2333 [(**22**+H)$^+$].

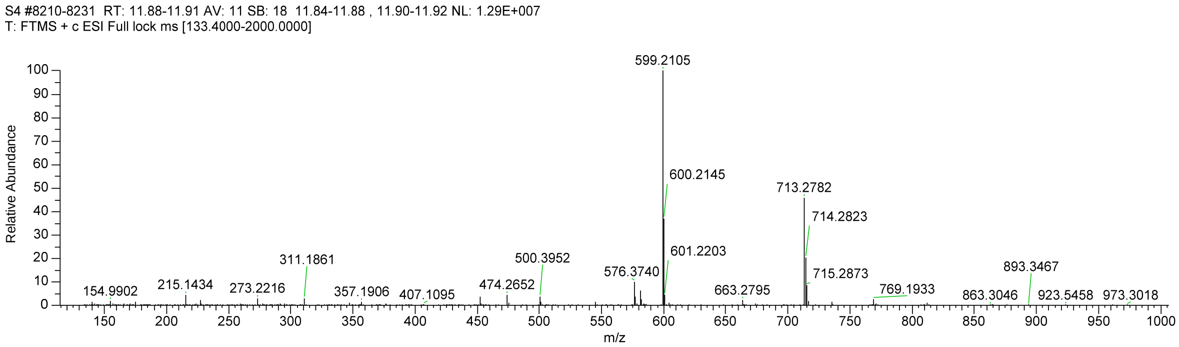


**Figure S58.** MS spectrum of compound **23**.

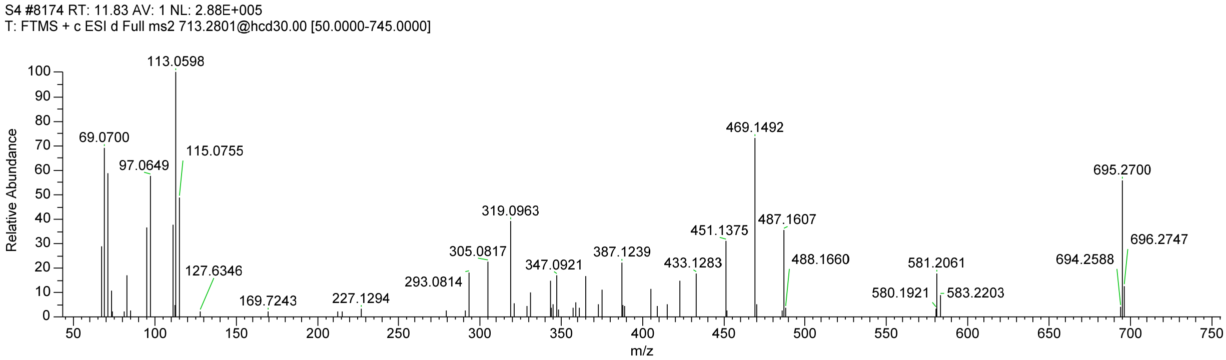


**Figure S59.** MS/MS spectrum of ion at *m/z* 713.2782 [(**23**+H)$^+$].

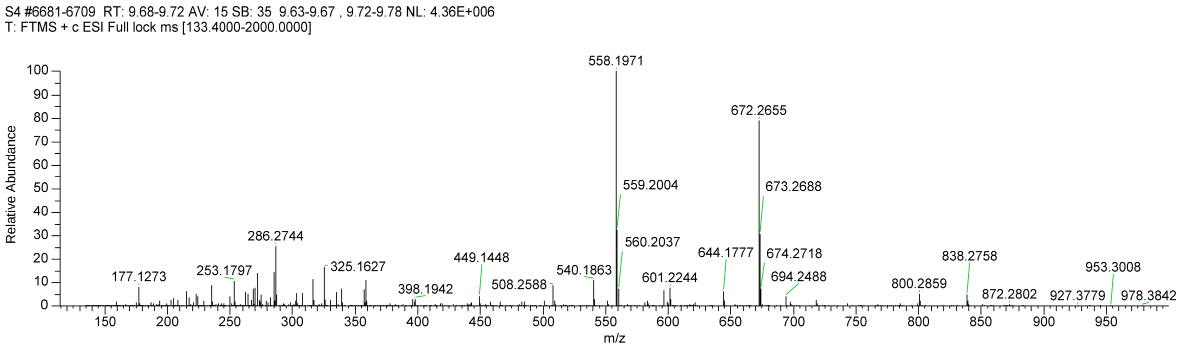


**Figure S60.** MS spectrum of compound **24**.

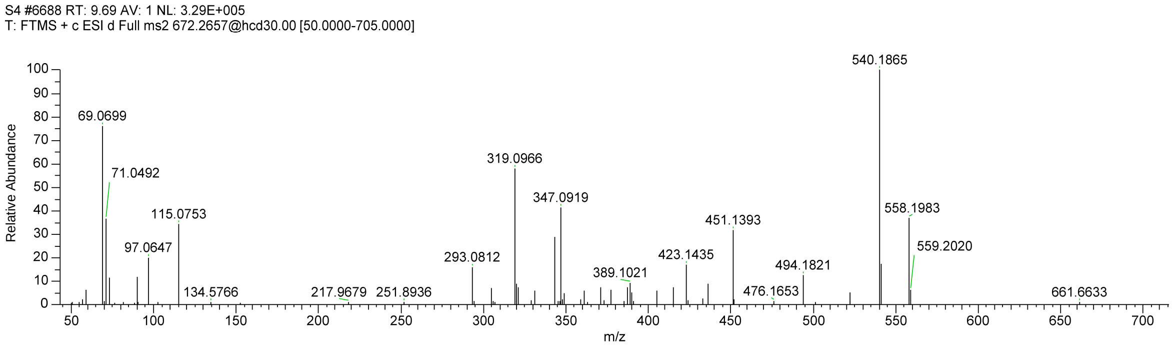


**Figure S61.** MS/MS spectrum of ion at *m/z* 672.2655 [(**24**+H)$^+$]

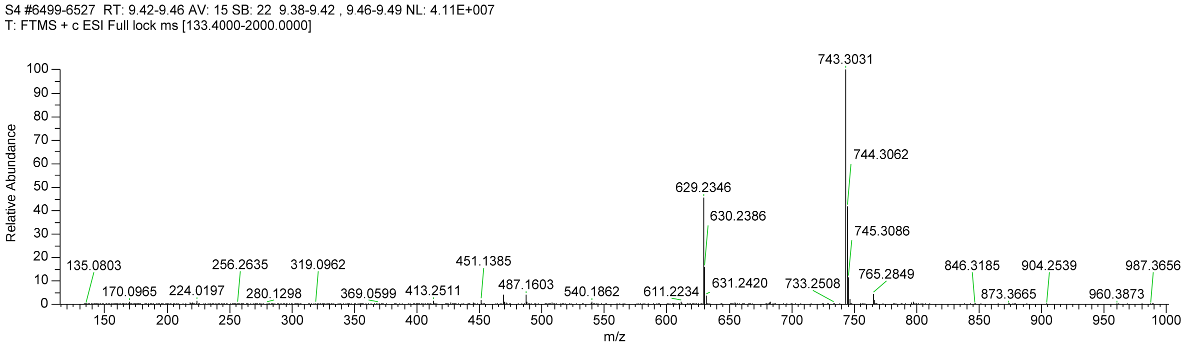


**Figure S62.** MS spectrum of compound **25**.

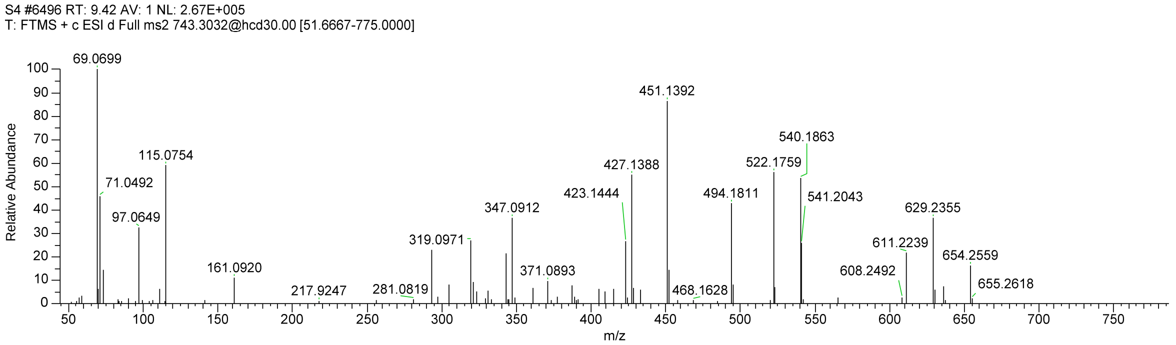


**Figure S63.** MS/MS spectrum of ion at *m/z* 743.3031 [(**25**+H)$^+$].

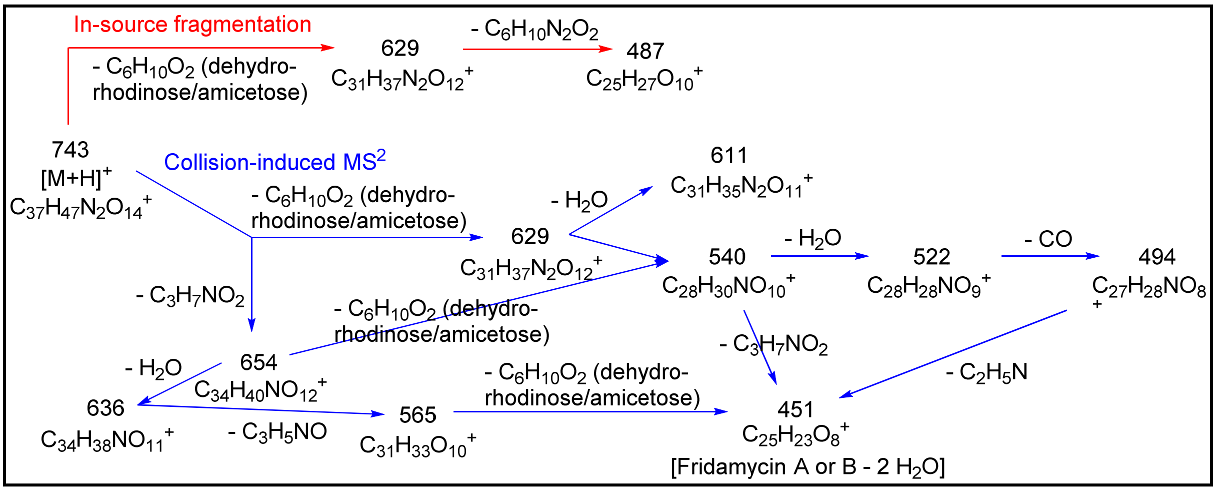


**Figure S64.** Fragmentation scheme for compound **25** from [M+H]$^+$ to the aglycone ion.

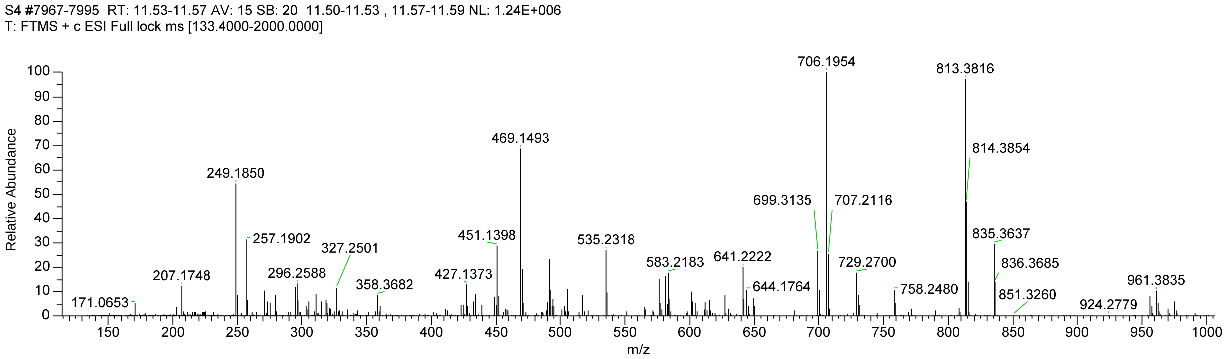


**Figure S65.** MS spectrum of compound **26**.

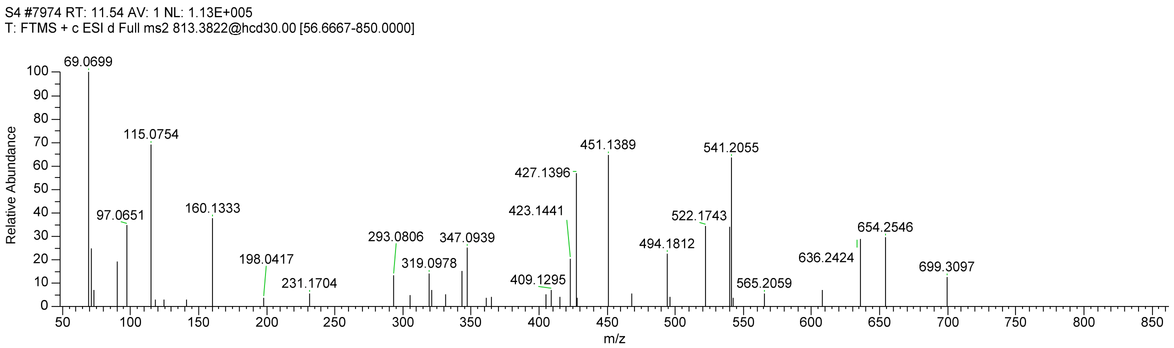


**Figure S66.** MS/MS spectrum of ion at *m/z* 813.3816 [(**26**+H)$^+$].

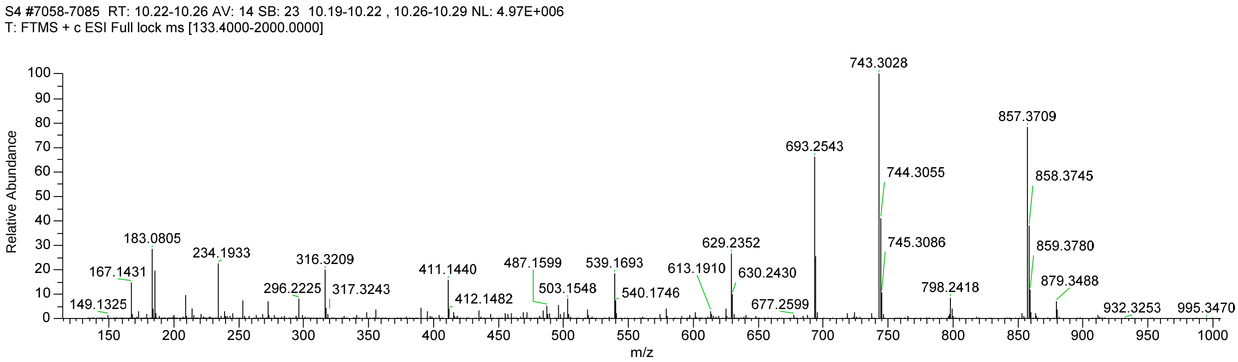


**Figure S67.** MS spectrum of compound **27**.

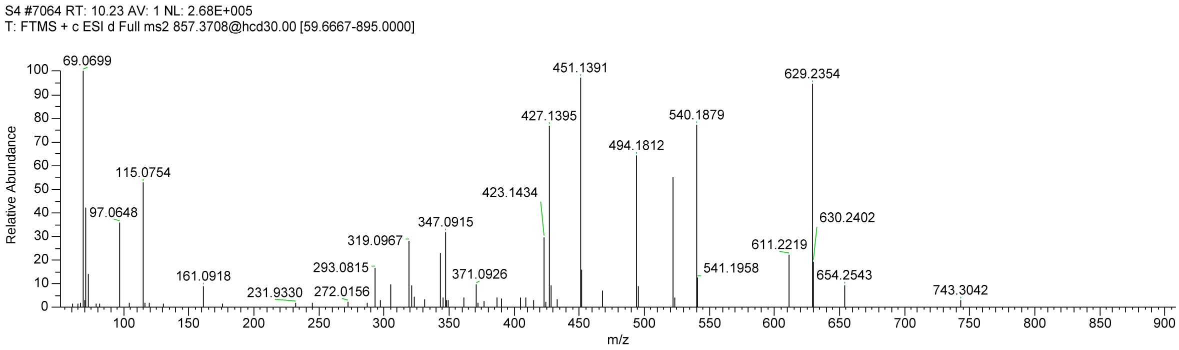


**Figure S68.** MS/MS spectrum of ion at *m/z* 857.3709 [(**27+H**)$^+$].

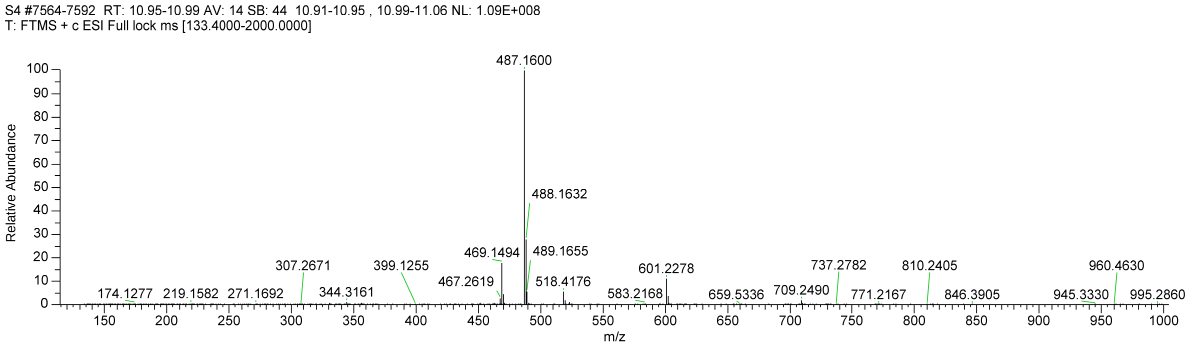


**Figure S69.** MS spectrum of compound **28**.

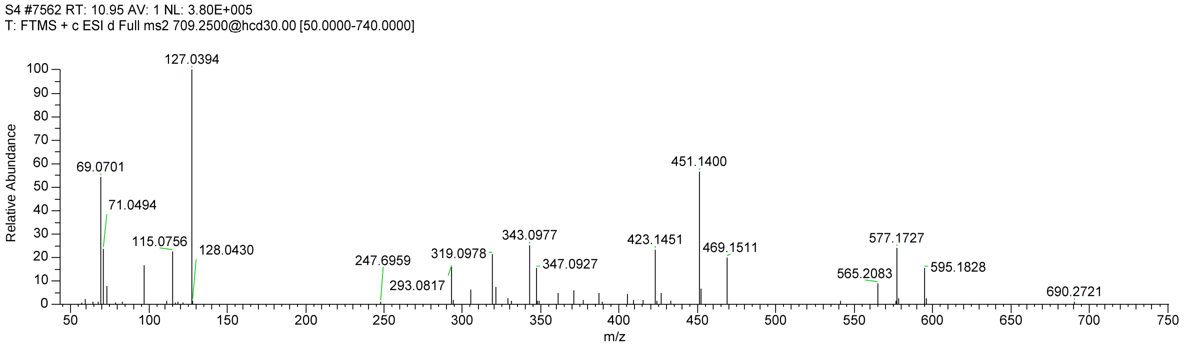


**Figure S70.** MS/MS spectrum of ion at *m/z* 709.2490 [(**28**+H)$^+$].

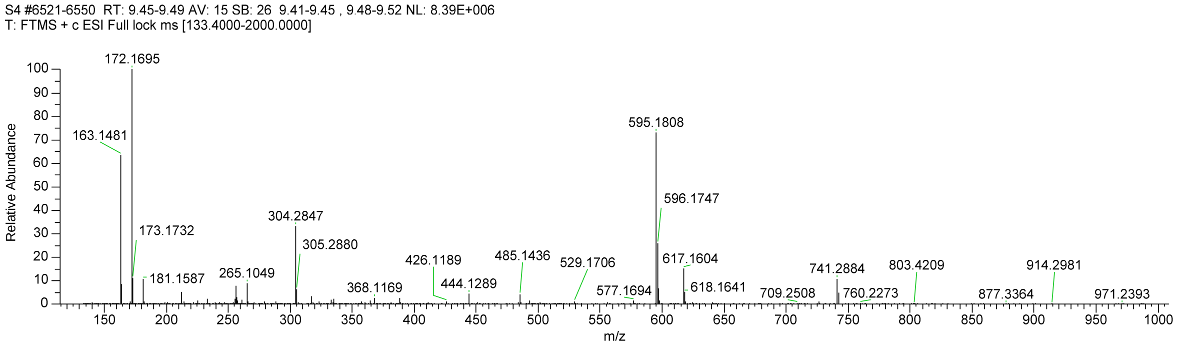


**Figure S71.** MS spectrum of compound **29**.

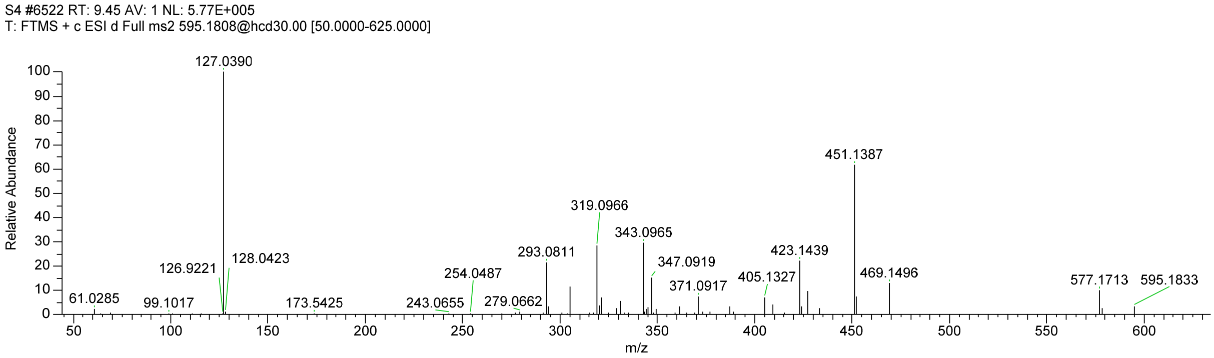


**Figure S72.** MS/MS spectrum of ion at *m/z* 595.1808 [(**29**+H)$^+$].

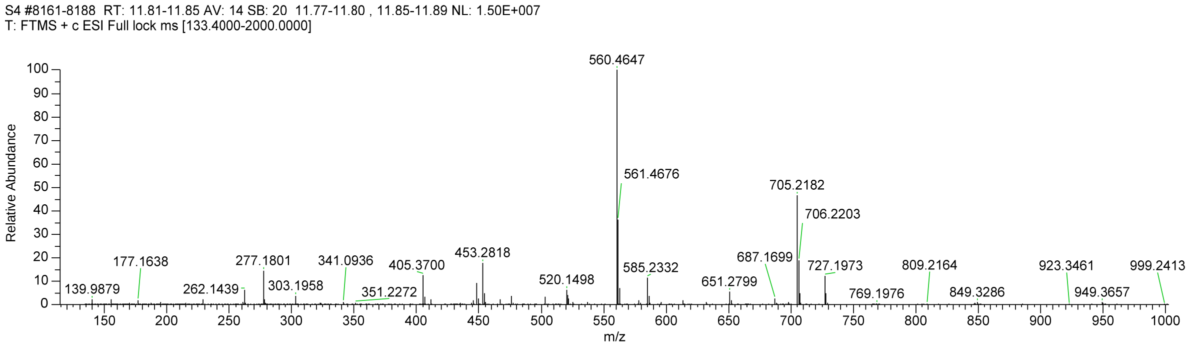


**Figure S73.** MS spectrum of compound **30**.

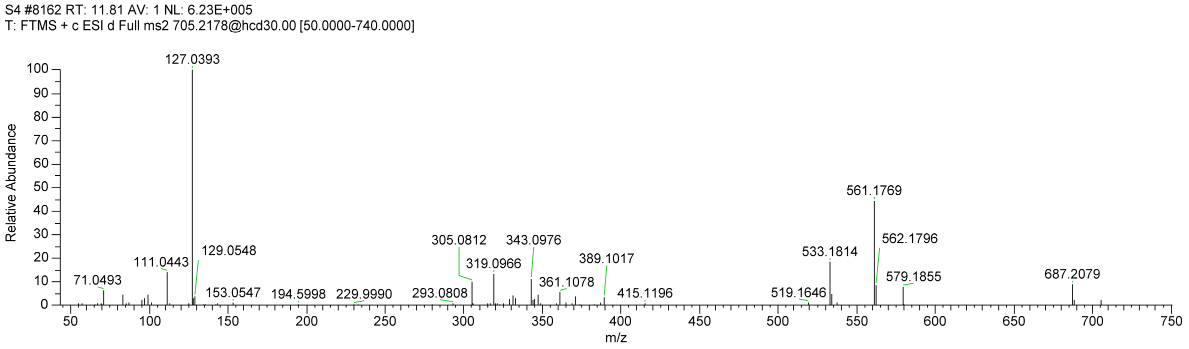


**Figure S74.** MS/MS spectrum of ion at *m/z* 705.2182 [(**30**+H)$^{+}$].

**Table S2.** Summary of antiSMASH 6.0 secondary metabolites predicted from the RO-S4 strain

| Cluster no. | Metabolite class | Most similar known biosynthetic cluster (%) | Contig no. | Span [nt] | |
|---|---|---|---|---|---|
| | | | | From | To |
| **Cluster 1** | Siderophore | Desferrioxamin B/ desferrioxamine E (83%) | ROS4_1 | 214,500 | 226,272 |
| **Cluster 2** | T2PKS, Oligosaccharide Phenazine, Siderophore | grincamycin (97%) | ROS4_2 | 140,051 | 226,533 |
| **Cluster 3** | Ectoine | ectoine (100%) | ROS4_4 | 76,930 | 87,328 |
| **Custer 4** | Lanthipeptide-class-i | kanamycin (1%) | ROS4_5 | 124,831 | 150,105 |
| **Cluster 5** | Terpene | geosmin (100%) | ROS4_ 6 | 126,687 | 148,843 |
| **Cluster 6** | T2PKS | spore pigment (83%) | ROS4_10 | 11,361 | 83,870 |
| **Cluster 7** | T3PKS | alkylresorcinol (100%) | ROS4_11 | 114,736 | 155,884 |
| **Cluster 8** | Terpene | carotenoid (54%) | ROS4_16 | 35,638 | 59,717 |
| **Cluster 9** | T1PKS | ashimides (12%) | ROS4_20 (1) | 259 | 24,193 |
| **Cluster 10** | RiPP-like, blactam, Terpene | Unknown | ROS4_20 (2) | 24,913 | 58,437 |
| **Cluster 11** | T1PKS | enduracidin (12%) | ROS4_20 (3) | 71,642 | 115,844 |
| **Cluster 12** | Phenazine | fredericamycin A (6%) | ROS4_21 | 49,206 | 69,667 |
| **Cluster 13** | Terpene | albaflavenone (100%) | ROS4_26 | 49,303 | 70,316 |
| **Cluster 14** | Terpene | hopene (92%) | ROS4_29 | 68,654 | 86,555 |
| **Cluster 15** | NRPS, T1PKS, other | polyoxypeptin (48%) | ROS4_33 | 1 | 82,128 |
| **Cluster 16** | Lanthipeptide-class-i | Unknown | ROS4_37 | 21,339 | 45,849 |
| **Cluster 17** | NRPS, T1PKS | antimycin (100%) | ROS4_41 | 1 | 47,509 |
| **Cluster 18** | RiPP-like | informatipeptin (42%) | ROS4_70 | 12,446 | 22,661 |
| **Cluster 19** | Butyrolactone | griseoviridin / fijimycin A (11%) | ROS4_89 | 1 | 6,885 |

**Table S3**. Deduced functions of the ORFs in the ROS4_2 cluster '*Streptomyces* sp. RO-S4 strain and their BLASTs. We have considered only the sequences that showed a homology >50%

| ORF | Gene annotation | Putative function | Length (nt) | Most similar protein/ Identity[a] | Strain | Accession Number |
|---|---|---|---|---|---|---|
| **1** | LRR80_00441 | Carboxymuconolactone decarboxylase | 1302 | forQ, 80.93%<br>Homolog in rubrolone, 82.04% | *Streptomyces sp. KY5*<br>*Streptomyces sp. KIB-H033* | AQP25549.1<br>AOZ61217.1 |
| **2** | LRR80_00442 | Hypothetical protein | 993 | n.a. | | |
| **3** | LRR80_00443 | Endoribonuclease | 396 | n.a. | | |
| **4** | LRR80_ 00444 | ATPase | 1380 | n.a. | | |
| **5** | LRR80_ 00445 | Phenazine biosynthesis-like protein | 858 | cnq727, 62.54% | *Streptomyces sp. CNQ-509* | AIT42117.1 |
| **6** | LRR80_ 00446 | Asparaginase | 1677 | n.a. | | |
| **7** | LRR80_00447 | Hypothetical protein unknown function | 486 | n.a. | | |
| **8** | LRR80_00448 | Putative methyltransferase | 1011 | Homolog in tetarimycin A, 60.90% | *uncultured bacterium* | AFY23025.1 |
| **9** | LRR80_00449 | 3-demethylubiquinone-9 3-methyltransferase | 447 | Homolog in SW-163C, 55.56% | *Streptomyces sp. SNA15896* | BAI63271.1 |
| **10** | LRR80_00450 | Phenazine biosynthesis protein A/B | 492 | Homolog in tetarimycin A, 74.66%<br>Homolog in streptophenazine B, 69.18% | *uncultured bacterium*<br>*Streptomyces sp. CNB091* | AFY23026.1<br>WP_018960032.1 |
| **11** | LRR80_00451 | Asparagine synthase | 1857 | PieD, 61.39%<br>ttmN, 60.84% | *Streptomyces sp. SCSIO 03032*<br>*Streptomyces afghaniensis* | AHE80999.1<br>ALJ49932.1 |
| **12** | LRR80_00452 | Pyridoxamine 5'-phosphate oxidase | 621 | LphzG, 53.93% | *Streptomyces lomondensis* | AKC91617.1 |
| **13** | LRR80_00453 | Hypothetical protein | 348 | Homolog in tetarimycin A ,61.05% | *uncultured bacterium* | AFY23027.1 |

| | | | | | | |
|---|---|---|---|---|---|---|
| **14** | LRR80_00454 | Isochorismate synthase | 1905 | stnM1, 62.62%<br>Homolog in diazaquinomycin A, 61.28% | *Streptomyces flocculus*<br>*Streptomyces sp. F001* | AFW04564.1<br>RZB16709.1 |
| **15** | LRR80_00455 | Isochorismatase | 624 | stnM2, 67.80%<br>Homolog, 68.14% | *Streptomyces flocculus*<br>*uncultured bacterium* | AFW04565.1<br>AFY23028.1 |
| **16** | LRR80_00456 | 3-deoxy-7-phosphoheptulonate synthase activity | 1125 | stnN, 64.72%<br>StrepF001_25935, 63.29%<br>epaO, 63.97% | *Streptomyces flocculus*<br>*Streptomyces sp. F001*<br>*Kitasatospora sp. HKI 714* | AFW04567.1<br>RZB16706.1<br>AHW81474.1 |
| **17** | LRR80_00457 | Hypothetical protein | 648 | n.a. | | |
| **18** | LRR80_00458 | Hypothetical protein | 375 | n.a. | | |
| **19** | LRR80_00459 | Regulatory protein | 366 | Homolog in bleomycin, 84.69% | *Streptomyces verticillus* | AAG02351.1 |
| **20** | LRR80_00460 | Hypothetical protein | 2415 | n.a. | | |
| **21** | LRR80_00461 | SNF2 Helicase protein | 2868 | Homolog in kosinostatin, 58.17% | *Micromonospora sp. TP-A0468* | AFJ52712.1 |
| **22** | LRR80_00462 | Hypothetical protein | 606 | n.a. | | |
| **23** | LRR80_00463 | Putative glucokinase | 1149 | n.a. | | |
| **24** | LRR80_00464 | Regulatory protein | 738 | n.a. | | |
| **25** | LRR80_00465 | Oxidoreductase | 1023 | n.a. | | |
| **26** | LRR80_00466 | Thioesterase superfamily protein | 435 | n.a. | | |
| **27** | LRR80_00467 | LuxR family DNA-binding response regulator | 675 | lipReg1, 85.00% | *Kitasatospora aureofaciens* | ABB05090.1 |
| **28** | LRR80_00468 | Histidine kinase | 1218 | lipReg25, 75.93% | *Kitasatospora aureofaciens* | ABB05091.1 |
| **29** | LRR80_00469 | Hypothetical protein | 204 | n.a. | | |
| **30** | LRR80_00470 | Ribonuclease | 429 | Pau53, 79.21%<br>lipX1, 66.90% | *Streptomyces Paulus*<br>*Kitasatospora aureofaciens* | AIE54220.1<br>ABB05092.1 |
| **31** | LRR80_00471 | Dehydrogenase/reductase SDR | 753 | Homolog in ebelactone, 59.52% | *Kitasatospora aburaviensis* | SCN11974.1 |

| | | | | | | |
|---|---|---|---|---|---|---|
| **32** | LRR80_00472 | Allantoicase | 1116 | n.a. | | |
| **33** | LRR80_00473 | Amidohydrolase | 1341 | PauY52, 77.53% | *Streptomyces sp. YN86* | AIE54272.1 |
| **34** | LRR80_00474 | IclR family transcriptional regulator | 807 | PauY51, 86.96% | *Streptomyces sp. YN86* | AIE54271.1 |
| **35** | LRR80_00475 | Hypothetical protein | 312 | PauY50, 56.38% | *Streptomyces sp. YN86* | AIE54270.1 |
| **36** | LRR80_00476 | NTP transferase | 600 | Orf-3, 86.43% | *Streptomyces lusitanus* | AGO50601.1 |
| **37** | LRR80_00477 | Malate synthase | 1626 | Orf-2, 94.27%<br>PauY45, 84.62% | *Streptomyces lusitanus*<br>*Streptomyces sp. YN86* | AGO50602.1<br>AIE54265.1 |
| **38** | LRR80_00478 | SelT/SelW/SelH family protein | 279 | n.a. | | |
| **39** | LRR80_00479 | Hypothetical protein | 924 | Orf-1, 93.81% | *Streptomyces lusitanus* | AGO50603.1 |
| **40** | LRR80_00480 | NADH: flavin oxidoreductase / NADH oxidase | 1119 | GcnA, 95.70%<br>spr3, 85.75 % | *Streptomyces lusitanus*<br>*Streptomyces sp. TK08046* | AGO50604.1<br>BAV16992.1 |
| **41** | LRR80_00481 | Drug resistance transporter | 1569 | GcnB, 92.87%<br>sqnB, 83.23% | *Streptomyces lusitanus*<br>*Streptomyces sp.* | AGO50605.1<br>ARO44650.1 |
| **42** | LRR80_00482 | NADPH-dependent FMN reductase | 600 | GcnC, 94.47%<br>sqnC, 85.43%<br>spr5, 85.93% | *Streptomyces lusitanus*<br>*Streptomyces sp.*<br>*Streptomyces sp. TK08046* | AGO50606.1<br>ARO44679.1<br>BAV16994.1 |
| **43** | LRR80_00483 | Regulatory proteins | 738 | GcnD, 94.69%<br>sqnD, 83.67 % | *Streptomyces lusitanus*<br>*Streptomyces sp.* | AGO50607.1<br>ARO44676.1 |
| **44** | LRR80_00484 | Hypothetical protein | 843 | GcnE, 80.71%<br>sprA, 71.73% | *Streptomyces lusitanus*<br>*Streptomyces sp. TK08046* | AGO50608.1<br>BAV16996.1 |

| | | | | | | |
|---|---|---|---|---|---|---|
| **45** | LRR80_00485 | Monooxygenase FAD-binding | 1488 | GcnF, 95.55% | *Streptomyces lusitanus* | AGO50609.1 |
| | | | | UrdE, 91.50% | *Streptomyces fradiae* | CAA60567.1 |
| | | | | sqnF, 89.68% | *Streptomyces sp.* | ARO44651.1 |
| | | | | schP10 86.76% | *Streptomyces sp. SCC 2136* | CAH10119.1 |
| | | | | PgaE, 75.82% | *Streptomyces sp. PGA64* | AAK57522.1 |
| | | | | lanE, 73.17% | *Streptomyces cyanogenus* | AAD13534.1 |
| | | | | bexE, 73.12% | *Amycolatopsis orientalis subsp. vinearia* | ADI71441.1 |
| | | | | ovmOI, 69.92% | *Streptomyces antibioticus* | ARK36153.1 |
| | | | | kinO2, 61.72% | *Streptomyces murayamaensis* | AAO65352.1 |
| **46** | LRR80_00486 | Polyketide synthesis cyclase | 327 | sprC, 96.26% | *Streptomyces sp. TK08046* | BAV16998.1 |
| | | | | sqnBB, 90.74% | *Streptomyces sp.* | ARO44683.1 |
| | | | | schP9, 90.65% | *Streptomyces sp. SCC 2136* | CAH10118.1 |
| **47** | LRR80_00487 | Beta-ketoacyl synthase | 1281 | GcnH, 98.12% | *Streptomyces lusitanus* | AGO50610.1 |
| | | | | sprD, 93.66% | *Streptomyces sp. TK08046* | BAV16999.1 |
| | | | | sqnH, 91.61% | *Streptomyces sp.* | ARO44655.1 |
| | | | | SchP8, 89.86% | *Streptomyces sp. SCC 2136* | CAH10117.1 |
| | | | | UrdA, 92.49% | *Streptomyces fradiae* | CAA60569.1 |
| **48** | LRR80_00488 | CLF Beta-ketoacyl synthase | 1221 | GcnI, 95.56% | *Streptomyces lusitanus* | AGO50611.1 |
| | | | | sprE, 90.42% | *Streptomyces sp. TK08046* | BAV17000.1 |
| | | | | sqnI, 86.70% | *Streptomyces sp.* | ARO44656.1 |
| | | | | OvmK, 74.57% | *Streptomyces antibioticus* | ARK36156.1 |
| | | | | URdB, 86.70% | *Streptomyces fradiae* | CAA60570 |
| **49** | LRR80_00489 | Acyl carrier protein | 270 | GcnJ, 98.88% | *Streptomyces lusitanus* | AGO50612.1 |
| | | | | sqnJ, 85.39% | *Streptomyces sp.* | ARO44684.1 |
| | | | | sprF, 84.27% | *Streptomyces sp. TK08046* | BAV17001.1 |
| **50** | LRR80_00490 | Dehydrogenase/reductase SDR | 786 | GcnK, 93.49 % | *Streptomyces lusitanus* | AGO50613.1 |
| | | | | sprG, 94.25% | *Streptomyces sp. TK08046* | BAV17002.1 |
| | | | | sqnK, 91.95% | *Streptomyces sp.* | ARO44671.1 |
| | | | | schP5, 88.63% | *Streptomyces sp. SCC 2136* | CAH10114.1 |

| | | | | | | |
|---|---|---|---|---|---|---|
| **51** | LRR80_00491 | Cyclase/Dehydrase | 936 | GcnL, 93.89%<br>sprH, 91.00%<br>UrdL, 86.17%<br>PgaL, 79.48%<br>May12, 77.17% | *Streptomyces lusitanus*<br>*Streptomyces sp. TK08046*<br>*Streptomyces fradiae*<br>*Streptomyces sp. PGA64*<br>*Streptomyces sp.* | AGO50614.1<br>BAV17003.1<br>AAF00205.1<br>AAK57529.1<br>AVO00811.1 |
| **52** | LRR80_00492 | Monooxygenase FAD-binding | 1989 | GcnM, 90.63%<br>SprI, 87.01%<br>UrdM, 79.82%<br>sqnM, 82.86%<br>schP3, 69.55%<br>Homolog in Baikalomycyin, 74.66%<br>pgaM, 75.00%<br>Homolog in Fridamycin A, 57.43%<br>ovmOII, 60.21%<br>bexM, 65.65%<br>SPW_4233, 64.90% | *Streptomyces lusitanus*<br>*Streptomyces sp. TK08046*<br>*Streptomyces fradiae*<br>*Streptomyces sp.*<br>*Streptomyces sp. SCC 2136*<br>*Streptomyces sp. IB201691-2A2*<br>*Streptomyces sp. PGA64*<br>*Actinomadura sp. RB99*<br>*Streptomyces antibioticus*<br>*Amycolatopsis orientalis subsp. Vinearia*<br>*Streptomyces sp. W007* | AGO50615.1<br>BAV17004.1<br>AFU51427.1<br>ARO44645.1<br>CAH10112.1<br>TRO56982.1<br>AAK57530.1<br>MBD2892419.1<br>CAG14970.1<br>ADI71448.1<br>EHM27510.1 |
| **53** | LRR80_00493 | Drug resistance transporter, EmrB/QacA | 1239 | GcnN, 90.26%<br>SprJ, 83.21%<br>saqN, 81.04% | *Streptomyces lusitanus*<br>*Streptomyces sp. TK08046*<br>*Streptomyces sp.* | AGO50616.1<br>BAV17005.1<br>ARO44657.1 |
| **54** | LRR80_00494 | Glycosyltransferase | 1293 | GcnG1, 96.98%<br>sqnG1, 88.14%<br>sprGT1, 86.74%<br>SchS10, 81.63% | *Streptomyces lusitanus*<br>*Streptomyces sp.*<br>*Streptomyces sp. TK08046*<br>*Streptomyces sp. SCC 2136* | AGO50617.1<br>ARO44654.1<br>BAV17006.1<br>CAH10110.1 |
| **55** | LRR80_00495 | Glycosyltransferase | 1197 | GcnG2, 92.46 %<br>sprGT2, 83.92%<br>sqnG2, 81.91 %<br>SchS9, 74.74% | *Streptomyces lusitanus*<br>*Streptomyces sp. TK08046*<br>*Streptomyces sp.*<br>*Streptomyces sp. SCC 2136* | AGO50618.1<br>BAV17007.1<br>ARO44658.1<br>CAH10109.1 |
| **56** | LRR80_00496 | dTDP-4-dehydrorhamnose 3,5-epimerase | 582 | GcnS1, 97.56%<br>sprK, 89.84%<br>sqnS1, 92.02% | *Streptomyces lusitanus*<br>*Streptomyces sp. TK08046*<br>*Streptomyces sp.* | AGO50619.1<br>BAV17008.1<br>ARO44680.1 |

| | | | | | | |
|---|---|---|---|---|---|---|
| **57** | LRR80_00497 | Glycosyltransferase | 1131 | GcnG3, 93.62% | *Streptomyces lusitanus* | AGO50620.1 |
| | | | | sprGT3, 89.89% | *Streptomyces sp. TK08046* | BAV17009.1 |
| | | | | sqnG3, 86.44% | *Streptomyces sp.* | ARO44660.1 |
| | | | | schS7, 78.13% | *Streptomyces sp. SCC 2136* | CAF31363.2 |
| | | | | UrdGT2, 77.81% | *Streptomyces fradiae* | AAF00209.1 |
| **58** | LRR80_00498 | Nucleotidyl transferase | 1068 | GcnS2, 95.21% | *Streptomyces lusitanus* | AGO50621.1 |
| | | | | sprL, 92.11% | *Streptomyces sp. TK08046* | BAV17010.1 |
| | | | | sqnS2, 89.86% | *Streptomyces sp.* | ARO44662.1 |
| **59** | LRR80_00499 | NAD dependent epimerase/dehydratase | 984 | GcnS3, 98.17% | *Streptomyces lusitanus* | AGO50622.1 |
| | | | | sprM, 93.56% | *Streptomyces sp. TK08046* | BAV17011.1 |
| | | | | sqnS3, 92.97% | *Streptomyces sp.* | ARO44664.1 |
| | | | | schS5, 88.65% | *Streptomyces sp. SCC 2136* | CAF31365.1 |
| **60** | LRR80_00500 | NAD dependent epimerase/dehydratase | 1059 | GcnS4, 88.43% | *Streptomyces lusitanus* | AGO50623 |
| | | | | sprN, 73.02% | *Streptomyces sp. TK08046* | BAV17012.1 |
| | | | | sqnS4, 79.08% | *Streptomyces sp.* | ARO44667.1 |
| | | | | schS4, 61.76% | *Streptomyces sp. SCC 2136* | CAF31366.1 |
| **61** | LRR80_00501 | DegT/DnrJ/EryC1/StrS aminotransferase | 1311 | GcnS5, 97.24% | *Streptomyces lusitanus* | AGO50624.1 |
| | | | | sprO, 93.12% | *Streptomyces sp. TK08046* | BAV17013.1 |
| | | | | sqnS5, 92.40% | *Streptomyces sp.* | ARO44653.1 |
| | | | | urdQ, 88.71% | *Streptomyces fradiae* | AAF72550.1 |
| | | | | schS3, 88.94% | *Streptomyces sp. SCC 2136* | CAF31367.1 |
| | | | | lanQ, 83.41% | *Streptomyces cyanogenus* | AAD13547.1 |
| | | | | aur1Q, 82.41% | *Streptomyces aureofaciens* | ACK77744.1 |
| | | | | saqQ, 80.46% | *Micromonospora sp. Tu 6368* | ACP19374.1 |
| | | | | lct46, 81.92% | *Streptomyces rishiriensis* | ABX71129.1 |

| | | | | | | |
|---|---|---|---|---|---|---|
| **62** | LRR80_00502 | NAD dependent epimerase/dehydratase | 765 | GcnS6, 92.91%<br>sprP, 88.98%<br>urdR, 78.86%<br>sqnS6, 84.25%<br>lanR, 73.06%<br>PgaR, 70.66% | *Streptomyces lusitanus*<br>*Streptomyces sp. TK08046*<br>*Streptomyces fradiae*<br>*Streptomyces sp.*<br>*Streptomyces cyanogenus*<br>*Streptomyces sp. PGA64* | AGO50625.1<br>BAV17014.1<br>AAF72551.1<br>ARO44673.1<br>AAD13548.1<br>AHW57781.1 |
| **63** | LRR80_00503 | Hypothetical protein | 954 | GcnO, 86.75%<br>sqnO, 74.92%<br>sprQ, 77.70% | *Streptomyces lusitanus*<br>*Streptomyces sp.*<br>*Streptomyces sp. TK08046* | AGO50626.1<br>ARO44665.1<br>BAV17015.1 |
| **64** | LRR80_00504 | NDP-hexose 2,3-dehydratase | 1404 | GcnS7, 94.00%<br>sprR, 87.37%<br>sqnS7, 85.81%<br>schS2, 84.29%<br>urdS, 78.52%<br>lanS, 74.50% | *Streptomyces lusitanus*<br>*Streptomyces sp. TK08046*<br>*Streptomyces sp.*<br>*Streptomyces sp. SCC 2136*<br>*Streptomyces fradiae*<br>*Streptomyces cyanogenus* | AGO50627.1<br>BAV17016.1<br>ARO44652.1<br>CAF31368.1<br>AAF72552.1<br>AAD13549.1 |
| **65** | LRR80_00505 | Oxidoreductase | 960 | GcnS8, 89.34 %<br>sqnS8, 83.65%<br>sprS, 86.79%<br>schS1, 76.85% | *Streptomyces lusitanus*<br>*Streptomyces sp.*<br>*Streptomyces sp. TK08046*<br>*Streptomyces sp. SCC 2136* | AGO50628.1<br>ARO44666.1<br>BAV17017.1<br>CAF31369.2 |
| **66** | LRR80_00506 | Acetyl-CoA carboxylase, carboxyl transferase | 1536 | GcnP, 97.85%<br>sqnP, 95.67%<br>sprU, 96.06%<br>Homolog in rubrolone A, 94.09% | *Streptomyces lusitanus*<br>*Streptomyces sp.*<br>*Streptomyces sp. TK08046*<br>*Streptomyces sp. KIB-H033* | AGO50629.1<br>ARO44649.1<br>BAV17019.1<br>AOZ61215.1 |
| **67** | LRR80_00507 | Acyl-CoA carboxylase epsilon subunit | 192 | ChlI, 62.07% | *Streptomyces antibioticus* | AAZ77683.1 |
| **68** | LRR80_00508 | FAD linked oxidase protein | 1590 | GcnQ, 95.09%<br>sqnQ, 86.96%<br>sprY, 84.69%<br>schA26, 72.18% | *Streptomyces lusitanus*<br>*Streptomyces sp.*<br>*Streptomyces sp. TK08046*<br>*Streptomyces sp. SCC 2136* | AGO50630.1<br>ARO44647.1<br>BAV17021.1<br>CAH10126.1 |

| | | | | | | |
|---|---|---|---|---|---|---|
| **69** | LRR80_00509 | Hypothetical protein | 438 | sqnEE, 84.03%<br>sprZ, 82.64% | *Streptomyces sp.*<br>*Streptomyces sp. TK08046* | ARO44681.1<br>BAV17022.1 |
| **70** | LRR80_00510 | Response regulator | 711 | GcnR, 95.34%<br>sprR3, 87.66%<br>sqnR, 86.86% | *Streptomyces lusitanus*<br>*Streptomyces sp. TK08046*<br>*Streptomyces sp.* | AGO50631.1<br>BAV17023.1<br>ARO44672.1 |
| **71** | LRR80_00511 | Oxidoreductase | 354 | SCO6269, 78.95%<br>schA18, 76.72%<br>sqnU, 71.55% | *Streptomyces coelicolor A3(2)*<br>*Streptomyces sp. SCC 2136*<br>*Streptomyces sp.* | CAB60188.1<br>CAF31373.1<br>ARO44661.1 |
| **72** | LRR80_00512 | Hypothetical protein | 246 | n.a. | | |
| **73** | LRR80_00513 | Integral membrane protein TerC family | 954 | Orf+1, 91.17% | *Streptomyces lusitanus* | AGO50634.1 |
| **74** | LRR80_00514 | Siderophore: IucA_IucC | 1623 | n.a. | | |
| **75** | LRR80_00515 | Siderophore: IucA_IucC | 1506 | n.a. | | |
| **76** | LRR80_00516 | Hypothetical protein | 1272 | n.a. | | |
| **77** | LRR80_00517 | Hypothetical protein | 801 | n.a. | | |
| **78** | LRR80_00518 | Hypothetical protein unknown function | 531 | arpX, 73.71% | *Streptomyces argillaceus* | SCO70312.1 |
| **79** | LRR80_00519 | YciI family protein | 549 | n.a. | | |
| **80** | LRR80_00520 | Transcriptional regulator | 627 | n.a. | | |
| **81** | LRR80_00521 | Aminotransferase | 1209 | n.a. | | |

[a] Identity in amino acid between predicted gene product of *Streptomyces* sp. RO-S4 and its nearest homologue.

n.a.: not annotated

```
gnl|LBBM|ROS4_2 [gcode=11] [organism=Streptomyces sp.] [strain=RO-S4]
1989 nucleotides in sequence
Mean G+C content = 72.8%

1.



               g
             t-a
             g-c
             c-g
             g-c
             g-c
             g-c
             t.t
             g-c     aa
            a   gccgc  a
    cct    c    +! !!  g
   g   gccg     tgacg  c
   g   !!!!     g    ct
   c   cggc      g
    ccg          g
            g-cc g
            g-c g
            g-c
            c-g
            g-c
           t   a
           t   c
            ggc


    tRNA-Ala(ggc)
    76 bases, %GC = 80.3
    Sequence c[568,643]
```

**Figure S75.** tRNAala predicted by Aragorn in the complementary strand of the LRR80_00492 ORF region.

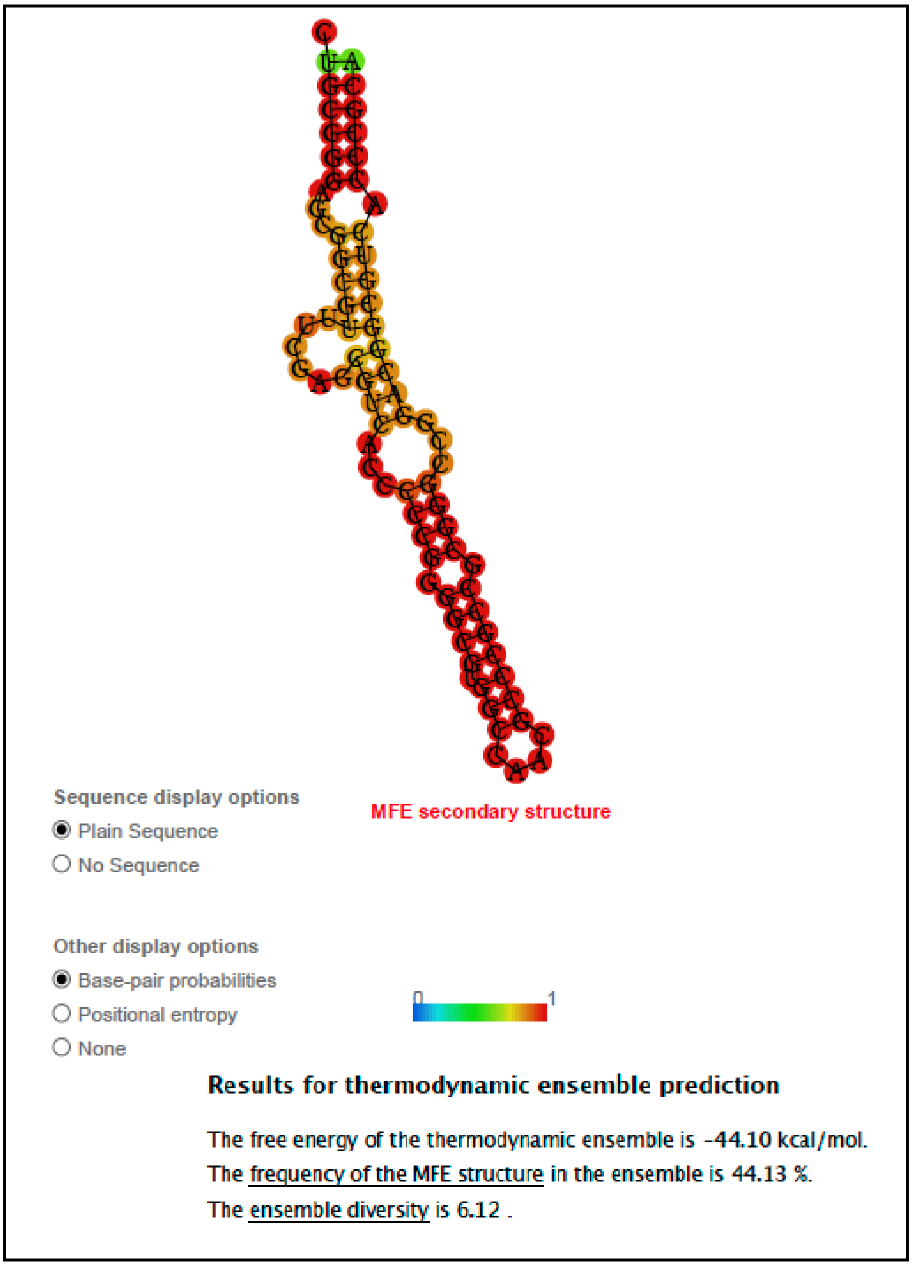


**Figure S76.** RNA-fold predicted RNA secondary structure of the region predicted by Aragorn to code (in the complementary strand) a tRNAala in the LRR80_00492 ORF region gene.

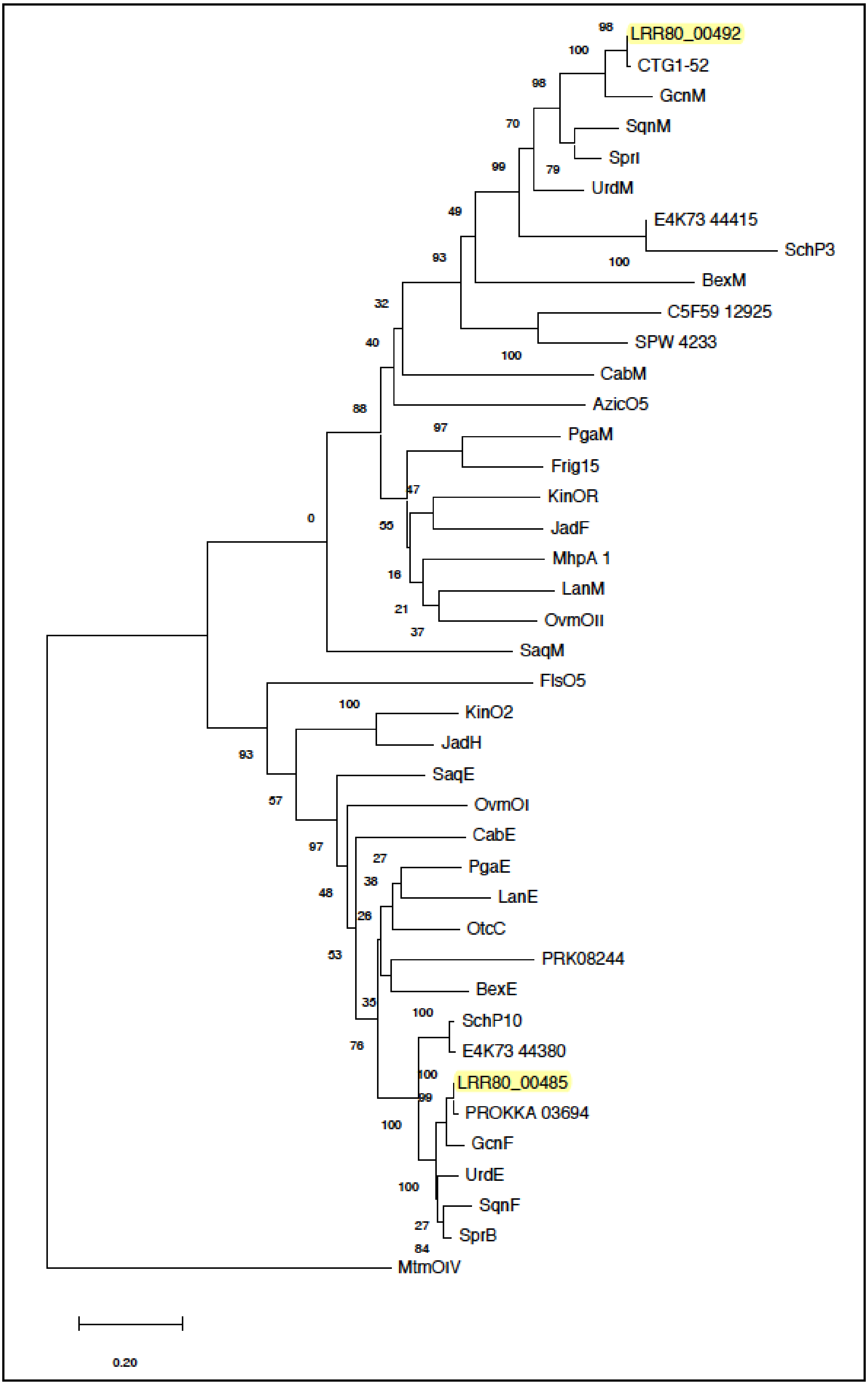
LRR80_00492
CTG1-52
GcnM
SqnM
SprI
UrdM
E4K73 44415
SchP3
BexM
C5F59 12925
SPW 4233
CabM
AzicO5
PgaM
Frig15
KinOR
JadF
MhpA 1
LanM
OvmOII
SaqM
FlsO5
KinO2
JadH
SaqE
OvmOI
CabE
PgaE
LanE
OtcC
PRK08244
BexE
SchP10
E4K73 44380
LRR80_00485
PROKKA 03694
GcnF
UrdE
SqnF
SprB
MtmOIV
0.20

**Figure S77.** Phylogenetic reconstruction (Neighbor Joining) of FAD dependent oxidases as described in the text. Where:

**PcpB** Pentachlorophenol 4-monooxygenase [*Sphingobium chlorophenolicum* L-1] AAF15368
**RdmE** Aklavinone C11-hydroxylase [*Streptomyces purpurascens* ATCC 25489] AAA83424
**TcmG** Tetracenomycin A2 hydroxylase [*Streptomyces glaucescens*] AAA67511
**DnrF** Aklavinone C-11 hydroxylase [*Streptomyces peucetius subsp. caesius* ATCC 27952] AAA62496
**LRR80_00492** Oxidase-Reductase (This study) [*Streptomyces* sp. S4]
**CTG1-52** Oxidase-Reductase (Grincamycins) [*Streptomyces* sp. CZN-748]
**GcnM** Oxygenase-reductase (Grincamycins) [*Streptomyces lusitanus* SCSIO LR32] AGO50615
**UrdM** Oxygenase-reductase (Urdamycin) [*Streptomyces fradiae* Tu 2717] AFU51427
**SqnM** Monooxygenase (Saquayamycin) [*Streptomyces* sp. KY40-1] ARO44645
**SprI** Oxygenase-reductase (Saprolmycin) [*Streptomyces* sp. TK08046] BAV17004
**C5F59_12925** SDR family oxidoreductase (Lugdunomycin) [*Streptomyces* sp. QL37] PPQ57494
**E4K73_44415** SDR family oxidoreductase (Baikalomycin) [*Streptomyces* sp. IB201691-2A2] WP_143644098
**SPW_4233** monooxygenase FAD-binding (Kiamycin) [*Streptomyces* sp. W007] EHM27510
**BexM** oxygenase-reductase (BE-7585A) [*Amycolatopsis orientalis subsp. vinearia*] ADI71448
**SchP3** oxidase-reductase (Sch 47554) [*Streptomyces* sp. SCC 2136] CAH10112
**PgaM** two-domain mono-oxygenase/angucyclinone (Gaudimycins) [*Streptomyces* sp. PGA64] AAK57530
**CabM** monooxygenase FAD-binding (Gaudimycin-B) [*Streptomyces* sp H021] DD106591
**MhpA_1** 3-(3-hydroxy-phenyl) propionate/3-hydroxycinnamic acid hydroxylase (Fridamycin-A) [*Actinomadura* sp. RB99] MBD2892419
**Frig15** oxygenase-reductase (Frigocyclinone) [*Streptomyces griseus* NTK97] QDG00821
**SaqM** putative oxygenase (Saquayamycin Z) [*Micromonospora* sp. Tu 6368] ACP19358
**LanM** oxygenase homolog (Landomycin) [*Streptomyces cyanogenus* S136] AAD13541
**MtmOIV** oxygenase (Mithramycin) [*Streptomyces argillaceus* ATCC 12956] CAK50794
**OvmOII** oxygenase (Oviedomycin) [*Streptomyces antibioticus* ATCC 11891] CAG14970
**AzicO5** oxygenase (Azicemicin) [*Kibdelosporangium* sp. MJ126-NF4] ADB02848
**FlsO5** polyketide oxygenase CabE (Fluostatin) [*Micromonospora rosaria* SCSIO N160] ALJ9986
**KinOR** oxygenase reductase-like protein (Kinamycin) [*Streptomyces murayamaensis*] AAO65351
**LRR80**_00485 monooxygenase FAD-binding (This study) [*Sreptomyces* sp.S4]
**PROKKA_03694** monooxygenase FAD-binding (Grincamycins) [*Streptomyces* sp. CZN-748]
**GcnF** Oxidoreductase [Streptomyces lusitanus SCSIO LR32] AGO50609
**SaqE** Putative oxygenase (Saquayamycin Z) [*Micromonospora* sp. Tu 6368] ACP19351
**SqnF** monooxygenase (Saquayamycin) [*Streptomyces* sp. KY40-1] ARO44651
**SprB** Putative oxygenase (Saprolmycin) [*Streptomyces* sp. TK08046] BAV16997
**SchP10** putative oxygenase (Sch 47554 and Sch 47555) [*Streptomyces* sp. SCC 2136] CAH10119
**PRK08244** FAD-dependent monooxygenase (Lugdunomycin) [*Streptomyces* sp. QL37] WP_104785789
**E4K73_44380** monooxygenase (Baikalomycins) [*Streptomyces* sp. IB201691-2A2] TRO56975
**PgaE** prejadomycin C12- and C12b- hydroxylase (Prejadomycin) [*Streptomyces* sp. PGA64] AAK57522
**OtcC** Anhydrotetracycline monooxygenase (Fridamycin-A) [*Actinomadura* sp. RB99] MBD2892418
**LanE** oxygenase homolog (Landomycin) [*Streptomyces cyanogenus* S136] AAD13534
**MtmOIV** oxygenase (Mithramycin) [*Streptomyces argillaceus* ATCC 12956] CAK50794
**OvmOI** oxygenase (Oviedomycin) [*Streptomyces antibioticus* NRRL 3238] ARK36153
**KinO2** oxygenase-like protein (Kinamycin) [*Streptomyces murayamaensis*] AAO65352
**BexE** putative oxygenase (BE-7585A) [*Amycolatopsis orientalis subsp. vinearia*] ADI71441
**CabE** monooxygenase FAD-binding (Gaudimycin-B) [*Streptomyces* sp H021] DD106591

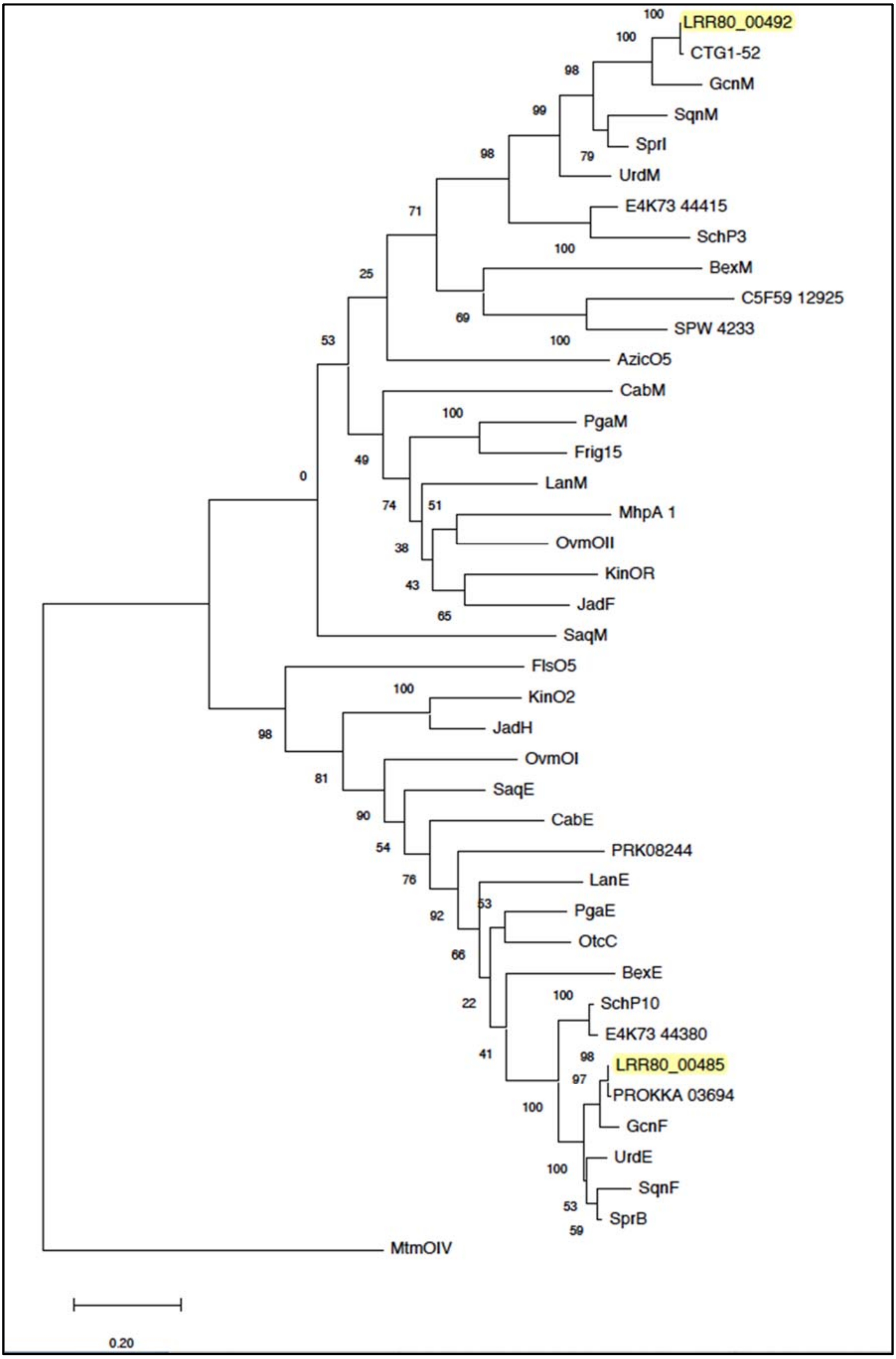
LRR80_00492
CTG1-52
GcnM
SqnM
SprI
UrdM
E4K73 44415
SchP3
BexM
C5F59 12925
SPW 4233
AzicO5
CabM
PgaM
Frig15
LanM
MhpA 1
OvmOII
KinOR
JadF
SaqM
FlsO5
KinO2
JadH
OvmOI
SaqE
CabE
PRK08244
LanE
PgaE
OtcC
BexE
SchP10
E4K73 44380
LRR80_00485
PROKKA 03694
GcnF
UrdE
SqnF
SprB
MtmOIV
0.20

**Figure S78.** Phylogenetic reconstruction (Maximum Likelihood) of FAD dependent oxidases as described in the text.

**PcpB** Pentachlorophenol 4-monooxygenase [*Sphingobium chlorophenolicum* L-1] AAF15368
**RdmE** Aklavinone C11-hydroxylase [*Streptomyces purpurascens* ATCC 25489] AAA83424
**TcmG** Tetracenomycin A2 hydroxylase [*Streptomyces glaucescens*] AAA67511
**DnrF** Aklavinone C-11 hydroxylase [*Streptomyces peucetius subsp. caesius* ATCC 27952] AAA62496
**LRR80_00492** Oxidase-Reductase (This study) [*Streptomyces* sp. RO-S4]
**CTG1-52** Oxidase-Reductase (Grincamycins) [*Streptomyces* sp. CNZ-748]
**GcnM** Oxygenase-reductase (Grincamycins) [*Streptomyces lusitanus* SCSIO LR32] AGO50615
**UrdM** Oxygenase-reductase (Urdamycin) [*Streptomyces fradiae* Tu 2717] AFU51427
**SqnM** Monooxygenase (Saquayamycin) [*Streptomyces* sp. KY40-1] ARO44645
**SprI** Oxygenase-reductase (Saprolmycin) [*Streptomyces* sp. TK08046] BAV17004
**C5F59_12925** SDR family oxidoreductase (Lugdunomycin) [*Streptomyces* sp. QL37] PPQ57494
**E4K73_44415** SDR family oxidoreductase (Baikalomycin) [*Streptomyces* sp. IB201691-2A2] WP_143644098
**SPW_4233** monooxygenase FAD-binding (Kiamycin) [*Streptomyces* sp. W007] EHM27510
**BexM** oxygenase-reductase (BE-7585A) [*Amycolatopsis orientalis subsp. vinearia*] ADI71448
**SchP3** oxidase-reductase (Sch 47554) [*Streptomyces* sp. SCC 2136] CAH10112
**PgaM** two-domain mono-oxygenase/angucyclinone (Gaudimycins) [*Streptomyces* sp. PGA64] AAK57530
**CabM** monooxygenase FAD-binding (Gaudimycin-B) [*Streptomyces* sp H021] DD106591
**MhpA_1** 3-(3-hydroxy-phenyl) propionate/3-hydroxycinnamic acid hydroxylase (Fridamycin-A) [*Actinomadura* sp. RB99] MBD2892419
**Frig15** oxygenase-reductase (Frigocyclinone) [*Streptomyces* griseus NTK97] QDG00821
**SaqM** putative oxygenase (Saquayamycin Z) [*Micromonospora* sp. Tu 6368] ACP19358
**LanM** oxygenase homolog (Landomycin) [*Streptomyces cyanogenus* S136] AAD13541
**MtmOIV** oxygenase (Mithramycin) [*Streptomyces argillaceus* ATCC 12956] CAK50794
**OvmOII** oxygenase (Oviedomycin) [*Streptomyces antibioticus* ATCC 11891] CAG14970
**AzicO5** oxygenase (Azicemicin) [*Kibdelosporangium* sp. MJ126-NF4] ADB02848
**FlsO5** polyketide oxygenase CabE (Fluostatin) [*Micromonospora rosaria* SCSIO N160] ALJ9986
**KinOR** oxygenase reductase-like protein (Kinamycin) [*Streptomyces murayamaensis*] AAO65351
**LRR80_00485** monooxygenase FAD-binding (This study) [*Streptomyces* sp. RO-S4]
**PROKKA_03694** monooxygenase FAD-binding (Grincamycins) [*Streptomyces* sp. CNZ-748]
**GcnF** Oxidoreductase [*Streptomyces lusitanus* SCSIO LR32] AGO50609
**SaqE** Putative oxygenase (Saquayamycin Z) [*Micromonospora* sp. Tu 6368] ACP19351
**SqnF** monooxygenase (Saquayamycin) [*Streptomyces* sp. KY40-1] ARO44651
**SprB** Putative oxygenase (Saprolmycin) [*Streptomyces* sp. TK08046] BAV16997
**SchP10** putative oxygenase (Sch 47554 and Sch 47555) [*Streptomyces* sp. SCC 2136] CAH10119
**PRK08244** FAD-dependent monooxygenase (Lugdunomycin) [*Streptomyces* sp. QL37] WP_104785789
**E4K73_44380** monooxygenase (Baikalomycins) [*Streptomyces* sp. IB201691-2A2] TRO56975
**PgaE** prejadomycin C12- and C12b- hydroxylase (Prejadomycin) [*Streptomyces* sp. PGA64] AAK57522
**OtcC** Anhydrotetracycline monooxygenase (Fridamycin-A) [*Actinomadura* sp. RB99] MBD2892418
**LanE** oxygenase homolog (Landomycin) [*Streptomyces cyanogenus* S136] AAD13534
**MtmOIV** oxygenase (Mithramycin) [*Streptomyces argillaceus* ATCC 12956] CAK50794
**OvmOI** oxygenase (Oviedomycin) [*Streptomyces antibioticus* NRRL 3238] ARK36153
**KinO2** oxygenase-like protein (Kinamycin) [*Streptomyces murayamaensis*] AAO65352
**BexE** putative oxygenase (BE-7585A) [*Amycolatopsis orientalis subsp. vinearia*] ADI71441
**CabE** monooxygenase FAD-binding (Gaudimycin-B) [*Streptomyces* sp H021] DD106591
